# Novelty and emergent patterns in sperm: morphological diversity and evolution of spermatozoa and sperm conjugation in ground beetles (Coleoptera: Carabidae)

**DOI:** 10.1101/809863

**Authors:** R. Antonio Gomez, David R. Maddison

## Abstract

1.

The beetle family Carabidae, with about 40,000 species, exhibits enough diversity in sperm structure and behavior to be an excellent model system for studying patterns and processes of sperm evolution. We explore their potential, documenting sperm form in 177 species of ground beetles and collecting data on 1 qualitative and 7 quantitative sperm phenotypic traits. Our sampling captures 61% of the tribal-level diversity of ground beetles. These data highlight the notable morphological diversity of sperm in ground beetles and suggest that sperm in the group have dynamic evolutionary histories with much morphological innovation and convergence. Sperm vary among species in total length from 48-3,400μm and in length and width of the sperm head. Most ground beetles make filamentous sperm with visually indistinct heads, but some or all studied members of the genus *Omophron,* genus *Trachypachus,* and tribe Dyschiriini make broad-headed sperm that show morphological differences between species. Most ground beetles package their sperm into groups of sperm, termed conjugates, and ground beetles show variation in conjugate form and in the number and arrangement of sperm in a conjugate. Most ground beetles make sperm conjugates by embedding their sperm in a non-cellular rod or spermatostyle, but some Trechitae make conjugates without a spermatostyle. The spermatostyle is remarkably variable among species and varies in length from 17-41,000μm. Several unrelated groups of ground beetles make only singleton sperm, including Nebriinae, Cicindelinae, many Trechinae, and the tribe Paussini. Given current views about ground beetle relationships, we propose preliminary hypotheses on ground beetle sperm diversification. We hypothesize that spermatostyle and conjugate traits evolve faster than sperm traits and that head width evolves more slowly than head length and sperm length. We propose that conjugation with a spermatostyle evolved early within the history of Carabidae and that it has been lost independently at least three times.

**Research highlights:**

- Ground beetle sperm is morphologically diverse.
- Most species make sperm conjugates with a spermatostyle, and there is variation in sperm, spermatostyles, and conjugates.
- Sperm have dynamic evolutionary histories.

## 2. Introduction

Animal sperm are among the most morphologically diverse cell type known. Although sperm have been described from thousands of species, patterns in sperm evolution remain largely unexplored (Birkhead and Montgomerie, 2009). Despite their few constituent parts, almost every part of a sperm cell that could be altered has been altered over evolutionary time, including the loss of cellular structures typical of sperm such as flagella and nuclei (see review by Pitnick et al., 2009a). Sperm live particularly odd “lives”, being launched away from the soma to face a variety of challenges unique among animal cells (Sivinski, 1984); variation in the environments sperm encounter is thought to account for their diversity of form.

There is also variation in how sperm travel upon leaving the male soma; some travel as singletons, but others travel in groups (Fig. 1), called conjugates. In sperm conjugates, two or more sperm cells join or are joined together for motility or transport through the female reproductive tract (see review by Higginson and Pitnick, 2011). Individual sperm in a conjugate frequently swim in a highly coordinated fashion (Taggart et al., 1993), and there is some evidence that an individual sperm’s form can be adaptive for conjugation (Immler et al., 2007; Taggart et al., 1993). Sperm conjugation is thought to be present in a small fraction of animal species but be taxonomically widespread. Given current phylogenetic hypotheses for relationships among animals (Hinchliff et al., 2015), it is likely that conjugation has evolved multiple times independently (Higginson and Pitnick, 2011).

**Figure 1.**
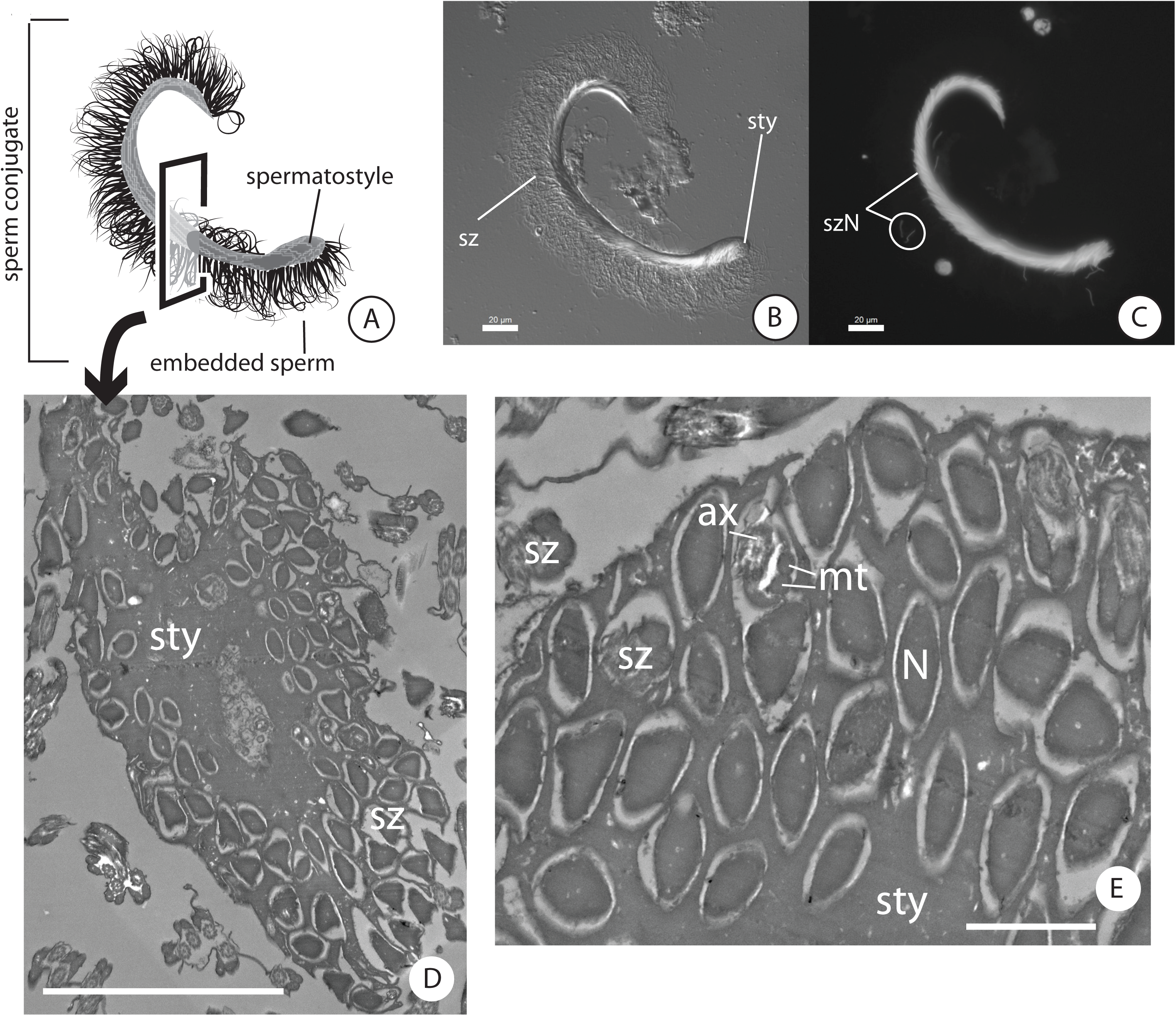
Many carabid beetles make sperm conjugates by pairing their sperm to a non-cellular structure or spermatostyle. (A) illustration of a *Scaphinotus marginatus* sperm conjugate (B, C). (B) DIC light microscope image of same. (C) fluorescence microscope image of DAPI-stained sperm heads. (D) TEM of a cross section through a *S. marginatus* sperm conjugate. (E) closeup of D showing details of indiviual sperm, ax = axoneme, mt = mitochondrial derivatives, sty = spermatostyle, sz = spermatozoa, szN = sperm nuclei. Scale bars: 1 μm (E), 5 μm (D).

In numerous animal clades, the striking variation of animal sperm form and function suggests that sperm are evolving rapidly and divergently. Rapid morphological divergence is common to reproductive traits and is a core prediction of evolution by sexual selection (Arnqvist and Rowe, 2005; Darwin, 1871; Eberhard, 1985; 1996; Holman and Snook, 2006; Hosken and Stockley, 2004; Miller and Pitnick, 2002; Parker, 1970; 1979; 2005; Pitnick and Hosken, 2010; Thornhill and Alcock, 1983). Although post-mating sexual selection is widely considered to be the mechanism driving sperm morphological variation, the adaptive function of most sperm traits is not known (Lüpold and Pitnick, 2018). Some sperm traits are recognized as exaggerated ornaments evolving under female choice or male persistence traits in sexual conflicts (e.g., Lupöld et al., 2016; Schärer et al., 2011).

The joining of sperm into groups of cells poses interesting broad scale evolutionary questions regarding how individual traits and group traits coevolve (Higginson and Pitnick, 2011). Do changes in sperm form drive changes in sperm conjugates such as the number of sperm in a group? Do the different components of a sperm conjugate show variation between species and if so, do they evolve at different rates? How do conjugates evolve? Is it the parts that change, or the arrangement of parts? Do females make decisions based upon a male’s sperm as well as his sperm conjugate? Do sperm cooperate (Fisher et al., 2014; Immler et al., 2007; Moore et al., 2002; Pizzari and Foster, 2008)? Although it is too early to draw conclusions about general processes from the available literature on the topic, early signs indicate that evolution of sperm conjugation is a fertile topic of investigation (e.g., Ferraguti et al., 1989; Fisher et al., 2014; Higginson et al., 2012a,b; Immler et al., 2007; Moore et al., 2002; Sasakawa, 2007).

Ground beetles (family Carabidae) are a large clade suitable as a study system for understanding the evolutionary patterns and processes of sexual trait evolution, as previous studies hint at diverse sperm forms. Carabid beetles are an old, varied family of terrestrial insects with nearly 40,000 described species (Lorenz, 2005; 2018). They reproduce sexually and have internal fertilization (Crowson, 1981). During copulation, males inseminate females, and females store sperm prior to fertilization (Crowson, 1981). Female reproductive tracts are morphologically diverse across the family, but all are of the “cul-de-sac” type with one duct leading to and away from the sperm storage organ (Liebherr and Will, 1998). Previous studies report variation in sperm across the species that have been studied (Supporting Information Table S1 and references therein). Ground beetle sperm vary in length from 68μm to 700μm (Takami and Sota, 2007; Sasakawa, 2009), and both sperm dimorphism and sperm conjugation are known to occur in the group (Supporting Information Table S1).

Although carabid beetles are a promising group in which to study sperm phenotypic evolution, essential data are lacking for most of the group’s diversity (Supporting Information Table S1). For example, most of the data (54 of the 69 studied species) come from only two genera, *Carabus* and *Pterostichus,* which are on widely separated branches of the tree of Carabidae. The near relatives of carabids, the diving beetles (Dytiscidae), are advancing as a system for studying sexual trait evolution (see review by Miller and Bergsten, 2014 and references therein). Diving beetles are known for their complex female reproductive tracts, diverse sperm forms with sperm length ranging from 128μm to 4493μm, three different qualitative types of sperm conjugation, and several, independently derived instances of dimorphism and/or conjugates that include more than one sperm morph (Higginson et al., 2012a,b). Carabid beetles are ten times as diverse as diving beetles, and if their sperm are variable like their near relatives, carabids are likely to provide numerous opportunities for studying the evolution of complex sperm traits.

The primary goal of the present study is to document sperm morphological diversity in ground beetles, making an effort to sample broadly across this diverse radiation of terrestrial insects by gathering data from as many lineages as possible within the family. We examine patterns and trends in sperm evolution in light of our results.

## 3. Materials & Methods

### Taxon sampling

Our study focused on identifying broad-scale patterns in sperm form across carabid beetles, and we prioritized capturing morphological variation of sperm across subfamilies and tribes within Carabidae. In total we studied 177 species of carabid beetle classified in 121 genera across 61 tribes or approximately 0.44%, 5.8%, and 61% of the known global diversity of carabid species, genera, and tribes (Table 1; Lorenz, 2005; 2018). Our attempt to sample different higher-level groups of ground beetles was guided by current classification of carabid beetles (e.g., Bousquet, 2012; Lorenz, 2005; 2018), current views about carabid relationships (e.g., Arndt et al., 2005; Figs. 3-4), and recent molecular phylogenetic studies of the group (Maddison et al., 1999; 2009; 2019; Ober, 2002; Ober and Maddison, 2008). Table 1 summarizes our sampling and includes the number of specimens studied per species by sex. We attempted to study multiple specimens per species in order to understand the stability of sperm traits within a species, and we averaged about two specimens per species (range = 1-8 specimens/species; Table 1).

**Table 1.** Taxon and specimen sampling for sperm data.

|  | Tribe | Genus | Species | #<br>male | #<br>female | total<br>(n) |
| --- | --- | --- | --- | --- | --- | --- |
| Carabinae | Carabini | *Calosoma* | *C. peregrinator* | 2 | - | 2 |
| Carabinae | Carabini | *Carabus* | *C. nemoralis* | 1 | - | 1 |
| Carabinae | Carabini | *Carabus* | *C. taedatus* | 1 | - | 1 |
| Carabinae | Cychrini | *Cychrus* | *C. tuberculatus* | - | 1 | 1 |
| Carabinae | Cychrini | *Scaphinotus* | *S. marginatus* | 1 | 1 | 2 |
| Carabinae | Cychrini | *Sphaeroderus* | *S. schaumii* | 2 | - | 2 |
| Carabinae | Cychrini | *Sphaeroderus* | *S. stenostomus* | 3 | 2 | 5 |
| Elaphrinae | Elaphrini | *Elaphrus* | *E. purpuratus* | 1 | 4 | 5 |
| Elaphrinae | Elaphrini | *Blethisa* | *B. oregonensis* | 2 | - | 2 |
| Trachypachinae | Trachypachini | *Trachypachus* | *T. inermis* | 1 | - | 1 |
| Trachypachinae | Trachypachini | *Trachypachus* | *T. slevini* | - | 4 | 4 |
| Loricarinae | Loricerini | *Loricera* | *L. decempunctata* | 1 | 1 | 2 |
| Loricarinae | Loricerini | *Loricera* | *L. foveata* | - | 2 | 2 |
| Nebriinae | Nebriini | *Nebria* | *N. brevicollis* | 1 | - | 1 |
| Nebriinae | Notiophilini | *Notiophilus* | *N. sylvaticus* | 3 | - | 3 |
| Nebriinae | Opisthiini | *Opisthius* | *O. richardsoni* | 1 | 1 | 2 |
| Omophroninae | Omophronini | *Omophron* | *O. americanum* | 1 | - | 1 |
| Omophroninae | Omophronini | *Omophron* | *O. ovale* | 4 | - | 4 |
| Trechinae | Patrobini | *Patrobus* | *P. longicornis* | 3 | 1 | 4 |
| Trechinae | Patrobini | *Diplois* | *D. filicornis* | - | 1 | 1 |
| Trechinae | Anillini | *gen. nov.* | Anillini *gen. nov. sp. nov.* | 3 | 1 | 4 |
| Trechinae | Bembidiini | *Bembidion* | *B. sp. nr. transversale* | 4 | - | 4 |
| Trechinae | Bembidiini | *Bembidion* | *B. incrematum* | 2 | 1 | 3 |
| Trechinae | Bembidiini | *Bembidion* | *B. iridescens* | 1 | 1 | 2 |
| Trechinae | Bembidiini | *Bembidion* | *B. kuprianovi* #2 | 1 | - | 1 |
| Trechinae | Bembidiini | *Bembidion* | *B. obliquulum* | 1 | 1 | 2 |
| Trechinae | Bembidiini | *Bembidion* | *B. sejunctum* | 1 | - | 1 |
| Trechinae | Bembidiini | *Bembidion* | *B. zephyrum* | 3 | - | 3 |
| Trechinae | Bembidiini | *Lionepha* | *L. sp. nov.* | 3 | - | 3 |
| Trechinae | Pogonini | *Diplochaetus* | *D. planatus* | 5 | - | 5 |
| Trechinae | Tachyini | *Mioptachys* | *M. flavicauda* | 3 | 1 | 4 |
| Trechinae | Tachyini | *Tachyta* | *T. inornata* | 1 | 1 | 2 |
| Trechinae | Tachyini | *Tachyura* | *T. rapax* | - | 1 | 1 |
| Trechinae | Tachyini | *Paratachys* | *P. sp. 1* | 1 | - | 1 |
| Trechinae | Tachyini | *Paratachys* | *P. sp. 2* | 2 | - | 2 |
| Trechinae | Trechini | *Pachydesus* | *P. sp.* | 3 | 1 | 4 |
| Trechinae | Trechini | *Perileptus* | *P. sp.* | 1 | 1 | 2 |
| Trechinae | Trechini | *Trechodes* | *T. sp.* | - | 1 | 1 |
| Trechinae | Trechini | *Trechosiella* | *T. scotti* | 3 | - | 3 |
| Trechinae | Trechini | *Trechus* | *T. humboldti* | 3 | 1 | 4 |
| Broscinae | Broscini | *Zacotus* | *Z. matthewsii* | 1 | 3 | 4 |
| Broscinae | Broscini | *Broscodera* | *B. insignis* | 1 | 1 | 2 |
| Non-Harpalinae<br>Carabidae<br>*incertae sedis* | Psydrini | *Psydrus* | *P. piceus* | - | 1 | 1 |
| Non-Harpalinae<br>Carabidae<br>*incertae sedis* | Gehringiini | *Gehringia* | *G. olympica* | 2 | - | 2 |
| Non-Harpalinae<br>Carabidae<br>*incertae sedis* | Apotomini | *Apotomus* | *A. sp.* | 1 | 3 | 4 |
| Non-Harpalinae<br>Carabidae<br>*incertae sedis* | Promecognathini | *Promecognathus* | *P. laevissimus* | 1 | 1 | 2 |
| Non-Harpalinae<br>Carabidae<br>*incertae sedis* | Hiletini | *Eucamaragnathus* | *E. oxygonus* | 1 | 2 | 3 |
| Scaritinae s. l. | Clivinini | *Ardistomis* | *A. obliquata* | 3 | - | 3 |
| Scaritinae s. l. | Clivinini | *Ardistomis* | *A. schaumii* | 5 | - | 5 |
| Scaritinae s. l. | Clivinini | *Aspidoglossa* | *A. subangulata* | 2 | - | 2 |
| Scaritinae s. l. | Clivinini | *Clivina* | *C. fossor* | 3 | - | 3 |
| Scaritinae s. l. | Clivinini | *Paraclivina* | *P. bipustulata* | 2 | - | 2 |
| Scaritinae s. l. | Clivinini | *Schizogenius* | *S. litigiosus* | 2 | 1 | 3 |
| Scaritinae s. l. | Clivinini | *Semiardistomis* | *S. viridis* | 6 | - | 6 |
| Scaritinae s. l. | Dyschiriini | *Akephorus* | *A. obesus* | 2 | 3 | 5 |
| Scaritinae s. l. | Dyschiriini | *Dyschirius* | *D. dejeanii* | 1 | - | 1 |
| Scaritinae s. l. | Dyschiriini | *Dyschirius* | *D. globosus* | 5 | - | 5 |
| Scaritinae s. l. | Dyschiriini | *Dyschirius* | *D. haemorrhoidalis* | 1 | - | 1 |
| Scaritinae s. l. | Dyschiriini | *Dyschirius* | *D. pacificus* | 1 | 1 | 2 |
| Scaritinae s. l. | Dyschiriini | *Dyschirius* | *D. thoracicus* | 4 | - | 4 |
| Scaritinae s. l. | Dyschiriini | *Dyschirius* | *D. tridentatus* | 1 | - | 1 |
| Scaritinae s. l. | Dyschiriini | *Striganoviella* | *S. vanhillei* | 5 | 1 | 6 |
| Scaritinae s. l. | Pasimachini | *Pasimachus* | *P. californicus* | 2 | 1 | 3 |
| Scaritinae s. l. | Scaritini | *Haplotrachelus* | *H. atropsis* | 1 | - | 1 |
| Scaritinae s. l. | Scaritini | *Haplotrachelus* | *H. cf. latesulcatus* | 2 | - | 2 |
| Scaritinae s. l. | Scaritini | *Haplotrachelus* | *H. sp.* | 2 | - | 2 |
| Scaritinae s. l. | Scaritini | *Scarites* | *S. marinus* | 2 | 1 | 3 |
| Scaritinae s. l. | Scaritini | *Scarites* | *S. (Distichus) sp.* | 1 | - | 1 |
| Scaritinae s. l. | Scaritini | *Scarites* | *S.*<br>*(Parallelomorphus)*<br>*sp.* | 2 | - | 2 |
| Rhysodinae | Clinidiini | *Clinidium* | *C. sp. nr.*<br>*guatemalenum* | 1 | 1 | 2 |
| Rhysodinae | Omoglymmiini | *Omoglymmius* | *O. hamatus* | 3 | - | 3 |
| Cicindelinae | Amblycheilini | *Omus* | *Omus audouini* | 5 | 2 | 7 |
| Cicindelinae | Amblycheilini | *Omus* | *Omus dejeanii* | 2 | 1 | 3 |
| Cicindelinae | Cicindelini | *Brasiella* | *B. wickhami* | 1 | - | 1 |
| Cicindelinae | Cicindelini | *Cicindela* | *C. haemorrhagica* | 2 | 1 | 3 |
| Cicindelinae | Megacephalini | *Tetracha* | *T. carolina* | 2 | 1 | 3 |
| Paussinae | Metriini | *Metrius* | *M. contractus* | 2 | 1 | 3 |
| Paussinae | Ozaenini | *Goniotropis* | *G. parca* | 3 | - | 3 |
| Paussinae | Ozaenini | *Ozaena* | *O. sp.* | - | 1 | 1 |
| Paussinae | Ozaenini | *Pachyteles* | *P. sp.* | - | 1 | 1 |
| Paussinae | Paussini | *Cerapterus* | *C. sp.* | 1 | - | 1 |
| Paussinae | Paussini | *Paussus* | *P. cucullatus* | - | 2 | 2 |
| Paussinae | Paussini | *Paussus* | *P. (Bathypaussus)*<br>*sp.* | 1 | - | 1 |
| Brachininae | Brachinini | *Brachinus* | *B. elongatulus* | 4 | 3 | 7 |
| Brachininae | Brachinini | *Brachinus* | *B. ichabodopsis* | 1 | - | 1 |
| Brachininae | Brachinini | *Mastax* | *M. sp.* | 3 | 1 | 4 |
| Brachininae | Brachinini | *Pheropsophus* | *P. sp. 1* | 4 | 1 | 5 |
| Brachininae | Brachinini | *Pheropsophus* | *P. sp. 2* | 1 | - | 1 |
| Harpalinae | Abacetini | *Abacetus* | *A. sp.* | 3 | 1 | 4 |
| Harpalinae | Abacetini | *Stolonis* | *S. intercepta* | 1 | 1 | 2 |
| Harpalinae | Abacetini | *Stolonis* | *S. sp.* | 1 | - | 1 |
| Harpalinae | Anthiini | *Anthia* | *Anthia*<br>*(Termophilum) sp.* | 1 | - | 1 |
| Harpalinae | Anthiini | *Cycloloba* | *C. sp.* | 1 | - | 1 |
| Harpalinae | Catapiesini | *Catapiesis* | *C. sp.* | - | 1 | 1 |
| Harpalinae | Chlaeniini | *Chlaenius* | *C. cumatilis* | 2 | - | 2 |
| Harpalinae | Chlaeniini | *Chlaenius* | *C. glaucus* | 1 | - | 1 |
| Harpalinae | Chlaeniini | *Chlaenius* | *C. harpalinus* | 1 | 1 | 2 |
| Harpalinae | Chlaeniini | *Chlaenius* | *C. leucoscelis* | 2 | - | 2 |
| Harpalinae | Chlaeniini | *Chlaenius* | *C. prasinus* | 1 | - | 1 |
| Harpalinae | Chlaeniini | *Chlaenius* | *C. ruficauda* | 2 | - | 2 |
| Harpalinae | Chlaeniini | *Chlaenius* | *C. sericeus* | 1 | - | 1 |
| Harpalinae | Chlaeniini | *Chlaenius* | *C. tricolor* | 1 | - | 1 |
| Harpalinae | Ctenodactylini | *Leptotrachelus* | *L. sp.* | 1 | - | 1 |
| Harpalinae | Cyclosomini | *Tetragonoderus* | *T. fasciatus* | 3 | 2 | 5 |
| Harpalinae | Cyclosomini | *Tetragonoderus* | *T. sp. nr. latipennis* | 1 | - | 1 |
| Harpalinae | Dryptini | *Drypta* | *D. sp.* | 2 | 1 | 3 |
| Harpalinae | Galeritini | *Galerita* | *G. atripes* | - | 1 | 1 |
| Harpalinae | Galeritini | *Galerita* | *G. bicolor* | - | 1 | 1 |
| Harpalinae | Galeritini | *Galerita* | *G. forreri* | 1 | - | 1 |
| Harpalinae | Galeritini | *Galerita* | *G. lecontei* | 2 | - | 2 |
| Harpalinae | Graphipterini | *Graphipterus* | *G. sp.* | 1 | - | 1 |
| Harpalinae | Harpalini | *Anisodactylus* | *A. alternans* | 1 | 2 | 3 |
| Harpalinae | Harpalini | *Anisodactylus* | *A. anthracinus* | 2 | - | 2 |
| Harpalinae | Harpalini | *Anisodactylus* | *A. similis* | 1 | - | 1 |
| Harpalinae | Harpalini | *Bradycellus* | *B. sp. 1* | 1 | - | 1 |
| Harpalinae | Harpalini | *Bradycellus* | *B. sp. 2* | 1 | - | 1 |
| Harpalinae | Harpalini | *Discoderus* | *D. sp.* | 1 | - | 1 |
| Harpalinae | Harpalini | *Euryderus* | *E. grossus* | 2 | - | 2 |
| Harpalinae | Harpalini | *Harpalus* | *H. affinis* | 2 | 2 | 4 |
| Harpalinae | Harpalini | *Polpochila* | *P. erro* | 2 | - | 2 |
| Harpalinae | Harpalini | *Selenophorus* | *S. sp.* | - | 1 | 1 |
| Harpalinae | Harpalini | *Stenolophus* | *S. sp.* | 1 | - | 1 |
| Harpalinae | Harpalini | *Stenomorphus* | *S. convexior* | 3 | - | 3 |
| Harpalinae | Helluonini | *Helluomorphoides* | *H. papago* | 1 | 1 | 2 |
| Harpalinae | Helluonini | *Macrocheilus* | *M. sp.* | 1 | - | 1 |
| Harpalinae | Lachnophorini | *Ega* | *E. sallei* | 1 | - | 1 |
| Harpalinae | Lachnophorini | *Lachnophorus* | *L. elegantulus* | 2 | - | 2 |
| Harpalinae | Lachnophorini | *Lachnophorus* | *L. sp. nr. elegantulus* | 1 | - | 1 |
| Harpalinae | Lebiini | *Agra* | *A. sp. 1* | 1 | - | 1 |
| Harpalinae | Lebiini | *Agra* | *A. sp. 2* | - | 1 | 1 |
| Harpalinae | Lebiini | *Apenes* | *A. lucidula* | 1 | - | 1 |
| Harpalinae | Lebiini | *Calleida* | *C. bella* | 1 | - | 1 |
| Harpalinae | Lebiini | *Calleida* | *C. decora* | - | 1 | 1 |
| Harpalinae | Lebiini | *Calleida* | *C. jansoni* | 2 | - | 2 |
| Harpalinae | Lebiini | *Cymindis* | *C. punctifera* | 1 | - | 1 |
| Harpalinae | Lebiini | *Cymindis* | *C. punctigera* | 1 | - | 1 |
| Harpalinae | Lebiini | *Cymindis* | *C. basipunctata-group sp.* | 1 | - | 1 |
| Harpalinae | Lebiini | *Lebia* | *L. deceptrix* | 1 | 1 | 2 |
| Harpalinae | Lebiini | *Lebia* | *L. subgrandis* | 1 | - | 1 |
| Harpalinae | Lebiini | *Lebia* | *L. viridis* | 1 | 1 | 2 |
| Harpalinae | Lebiini | *Phloeoxena* | *P. nigricollis* | 1 | - | 1 |
| Harpalinae | Lebiini | *Stenognathus* | *S. quadricollis* | - | 1 | 1 |
| Harpalinae | Lebiini | *Syntomus* | *S. americanus* | 1 | 1 | 2 |
| Harpalinae | Lebiini | *Thyreopterus* | *T. flavosignatus* | 1 | - | 1 |
| Harpalinae | Licinini | *Badister* | *B. ferrugineus* | 1 | - | 1 |
| Harpalinae | Licinini | *Dicaelus* | *D. suffusus* | 1 | 2 | 3 |
| Harpalinae | Licinini | *Diplocheila* | *D. nupera* | 1 | - | 1 |
| Harpalinae | Morionini | *Morion* | *M. sp.* | 1 | - | 1 |
| Harpalinae | Odacanthini | *Colliuris* | *C. pennsylvanica* | 3 | 1 | 4 |
| Harpalinae | Oodini | *Anatrichis* | *A. minuta* | 1 | - | 1 |
| Harpalinae | Oodini | *Oodes* | *O. fluvialis* | 8 | - | 8 |
| Harpalinae | Oodini | *Stenocrepis* | *S. elegans* | 1 | - | 1 |
| Harpalinae | Panagaeini | *Panagaeus* | *P. sallei* | 3 | - | 3 |
| Harpalinae | Peleciini | *Disphaericus* | *D. sp.* | 1 | 1 | 2 |
| Harpalinae | Pentagonicini | *Pentagonica* | *P. sp.* | 2 | - | 2 |
| Harpalinae | Perigonini | *Perigona* | *P. nigriceps* | 1 | - | 1 |
| Harpalinae | Platynini | *Agonum* | *A. piceolum* | 4 | 1 | 5 |
| Harpalinae | Platynini | *Agonum* | *A. muelleri* | 2 | - | 2 |
| Harpalinae | Platynini | *Rhadine* | *R. dissecta*-group<br>*sp.* | 1 | - | 1 |
| Harpalinae | Platynini | *Sericoda* | *S. bembidioides* | 1 | - | 1 |
| Harpalinae | Pseudomorphini | *Pseudomorpha* | *P. sp.* | - | 1 | 1 |
| Harpalinae | Pterostichini | *Abaris* | *A. splendida* | 2 | 1 | 3 |
| Harpalinae | Pterostichini | *Cyrtomoscelis* | *C. cf. dwesana* | 3 | - | 3 |
| Harpalinae | Pterostichini | *Cyclotrachelus* | *C. dejeanellus* | 2 | 1 | 3 |
| Harpalinae | Pterostichini | *Hybothecus* | *H. flohri* | - | 1 | 1 |
| Harpalinae | Pterostichini | *Poecilus* | *P. laetulus* | 1 | 1 | 2 |
| Harpalinae | Pterostichini | *Poecilus* | *P. scitulus* | - | 1 | 1 |
| Harpalinae | Pterostichini | *Pterostichus* | *P. infernalis* | 2 | 3 | 5 |
| Harpalinae | Pterostichini | *Pterostichus* | *P. lama* | 1 | - | 1 |
| Harpalinae | Pterostichini | *Pterostichus* | *P. melanarius* | - | 1 | 1 |
| Harpalinae | Sphodrini | *Calathus* | *C. peropacus* | 2 | 1 | 3 |
| Harpalinae | Sphodrini | *Synuchus* | *S. dubius* | 3 | 2 | 5 |
| Harpalinae | Zabrini | *Amara* | *A. aenea* | 6 | 2 | 8 |
| Harpalinae | Zabrini | *Amara* | *A. farcta* | - | 1 | 1 |
| Harpalinae | Zuphiini | *Pseudaptinus* | *P. horni* | - | 1 | 1 |
| Harpalinae | Zuphiini | *Pseudaptinus* | *P. simplex* | - | 1 | 1 |
| Harpalinae | Zuphiini | *Pseudaptinus* | *P. tenuicollis* | 1 | - | 1 |

### Specimens

Our study is based on a total of 397 specimens (Tables 1, 2, S2). We collected live beetles for sperm morphology in the United States, Mexico, the Republic of South Africa, and Mozambique. We also studied additional specimens preserved in 10% neutral-buffered formalin from Germany and Guatemala (Supporting Information Table S2). 10% neutral-buffered formalin has a long history of use in sperm morphology, and recent evidence from passerine birds suggests that it does not alter the form of sperm (Schmoll et al., 2016).

We kept beetles alive in small containers separated by collection locality and species prior to dissection or preservation in neutral-buffered formalin. When possible we stored the beetles in a refrigerator or cooler to limit movement and increase longevity. Following dissection and slide preparation, we associated slides with their parent specimens with the use of unique alphanumeric codes given to each specimen. We attempted to identify all of our specimens to species with the aid of taxonomic literature or help from taxonomic specialists (see Acknowledgements). If we were unable to identify a specimen to species because it represents an undescribed species or is part of group in need of revision, the specimen was only identified to genus.

### Phylogeny

The phylogenetic relationships of most ground beetles sampled for sperm data are not well understood, which limits insights into carabid sperm evolution. We use a low-resolution phylogenetic hypothesis of ground beetles to guide our interpretation of sperm data (Figs. 3-4). This phylogenetic hypothesis is based on published phylogenies with minor contributions from traditional classifications of ground beetles. The tree’s shape is predominately derived from large-scale molecular studies of ground beetle phylogeny (Maddison et al., 1999; 2009; Ober, 2002; Ober and Maddison, 2008). Additional molecular phylogenetic studies provided support for relationships in the following clades: Carabinae (Osawa et al., 2004), Cicindelinae (Vogler and Pearson, 1996; Gough et al., 2018), Harpalini (Martínez-Navarro et al., 2005), Paussinae (Moore, 2008; Robertson and Moore, 2016), Pterostichini and allies (Will and Gill, 2008), and Trechinae (Maddison and Ober, 2011; Maddison et al., 2019).

### Use of terms in sperm conjugation

The study of sperm conjugation has been complicated by the variation in sperm conjugation across animals and the historical lack of standard terms to refer to these structures and their method of development (Higginson and Pitnick, 2011). Previous workers have referred to the sperm conjugates of carabid beetles by a variety of terms such as sperm bundles (e.g., Hodgson et al., 2013), spermatodesms (Sasakawa, 2007; Sasakawa and Toki, 2008), or spermiozeugmata or similar (e.g., Ferenz, 1986; Schubert et al., 2017). Higginson and Pitnick (2011) suggest restricting the use of terms like these to particular morphological and developmental patterns. Higginson and Pitnick (2011) identified two major types of conjugation: primary and secondary. Primary conjugates like spermatodesms result from the products of a single spermatogonium remaining grouped together following spermiogenesis (Higginson and Pitnick, 2011). Secondary conjugates like sperm bundles result from sperm becoming joined together after individualization with sperm that are not necessarily from the same cyst (Higginson and Pitnick, 2011).

Data are still lacking regarding whether the sperm conjugates of carabid beetles are primary or secondary conjugates. Evidence from beetles in the closely related families Haliplidae and Gyrinidae that make conjugates that look similar to those in many carabids (Breland and Simmons, 1970; Higginson and Pitnick, 2011) suggests that they are spermatodesms. Schubert et al., (2017) studied the reproductive tract and spermatogenesis in a carabid beetle, *Limodromus assimilis,* and came to a different conclusion. They examined various sections of the male internal tract in this beetle and found that sperm individualize prior to becoming joined together with a hyaline rod (Schubert et al., 2017). It is still unclear whether the sperm conjugates of *L. assimilis* are composed of sperm derived from a single spermatogonium, which is necessary for it to be considered to be primary conjugation. The form of the male internal tract of *L. assimilis* is also highly similar to whirligig beetles in the genus *Dineutus,* which are known to make spermatodesms (Pitnick, unpublished data). Because of the uncertainty in conjugate type in carabid beetles studied thus far and the lack of data for the overwhelming majority of species in the family, we choose to refer to these multi-sperm forms by the neutral term conjugate (Fig. 1).

We classified the variation in conjugation we observed into different qualitative discrete types (Figs. 2-3). If we did not observe any physical association between two or more sperm, we considered those species to lack conjugation. The conjugates of species that make sperm that are physically associated via their heads with a hyaline rod, or spermatostyle, with unbounded flagella were considered rod conjugates. Conjugates characterized by sperm with flagella that are bounded to a spermatostyle were considered sheet conjugates following Sasakawa, (2007). Those with sperm joined together via their heads and cementing material but without a spermatostyle were considered aggregate conjugates (Higginson et al., 2012a). In some rare cases, we observed species that make singleton sperm bounded to a spermatostyle with a 1-to-1 match between sperm and spermatostyle. We did not consider this to be an example of sperm conjugation. Conjugates that form as a result of sperm grappling onto one another in a seemingly imprecise location were considered mechanical conjugates reminiscent of the sperm trains of muroid rodents (Higginson and Pitnick, 2011).

**Figure 2.**
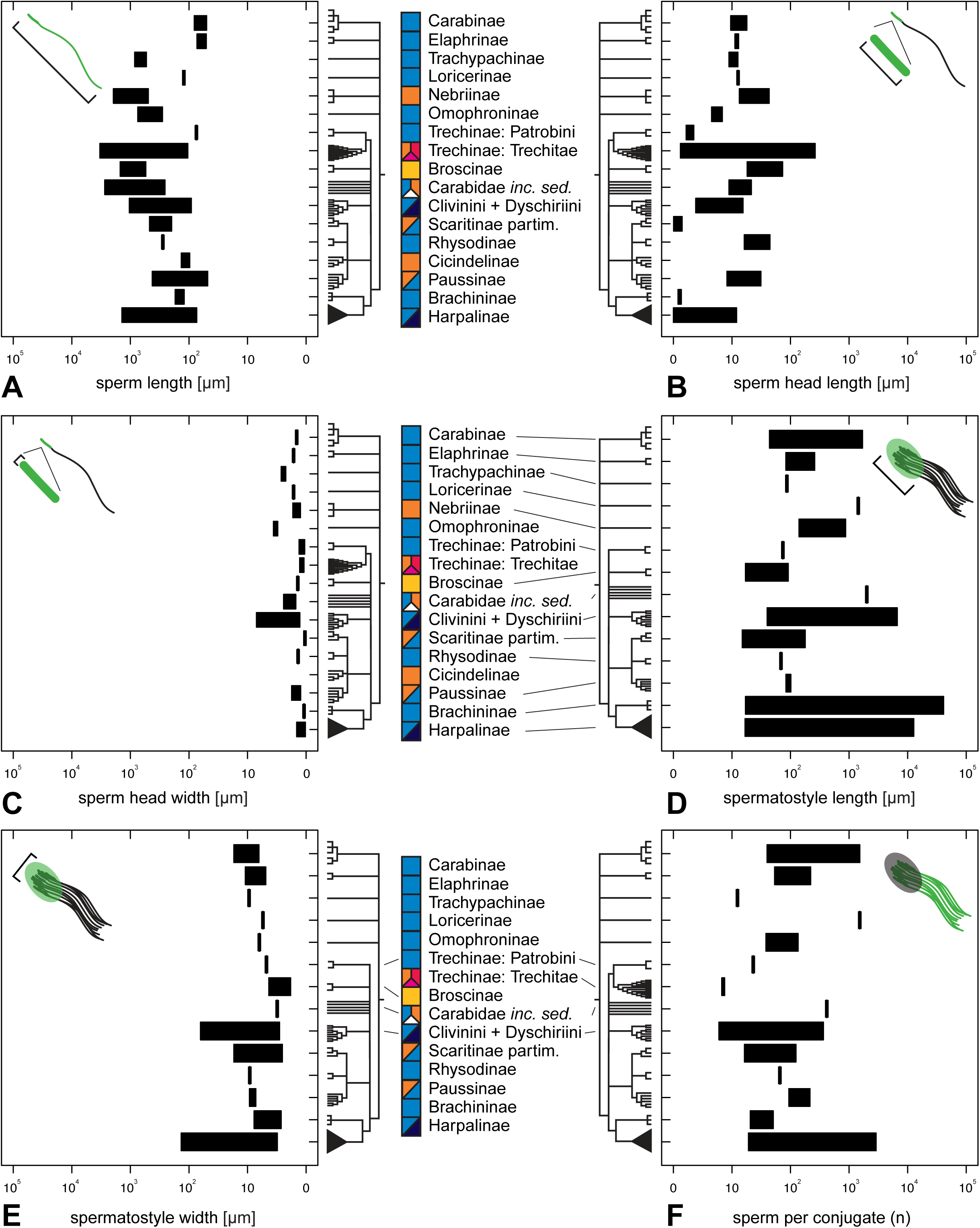
Variation in studied sperm quantitative traits across major taxonomic groupings of ground beetles (on logarithmic scale; lengths in μm). Colored boxes beside taxon names refer to qualitative conjugate types (see text and Fig. 3). (A) sperm length. (B) sperm head length. (C) sperm head width. (D) spermatostyle length. (E) spermatostyle width. (F) number of sperm included in a conjugate. Note that there are fewer rows in plots D-F because some ground beetles do not make a spermatostyle and/or they lack sperm conjugation. In order to avoid negative transformed values for small-headed sperm in plots B and C, we adjusted all values by 1.2μm prior to log transformation.

**Figure 3.**
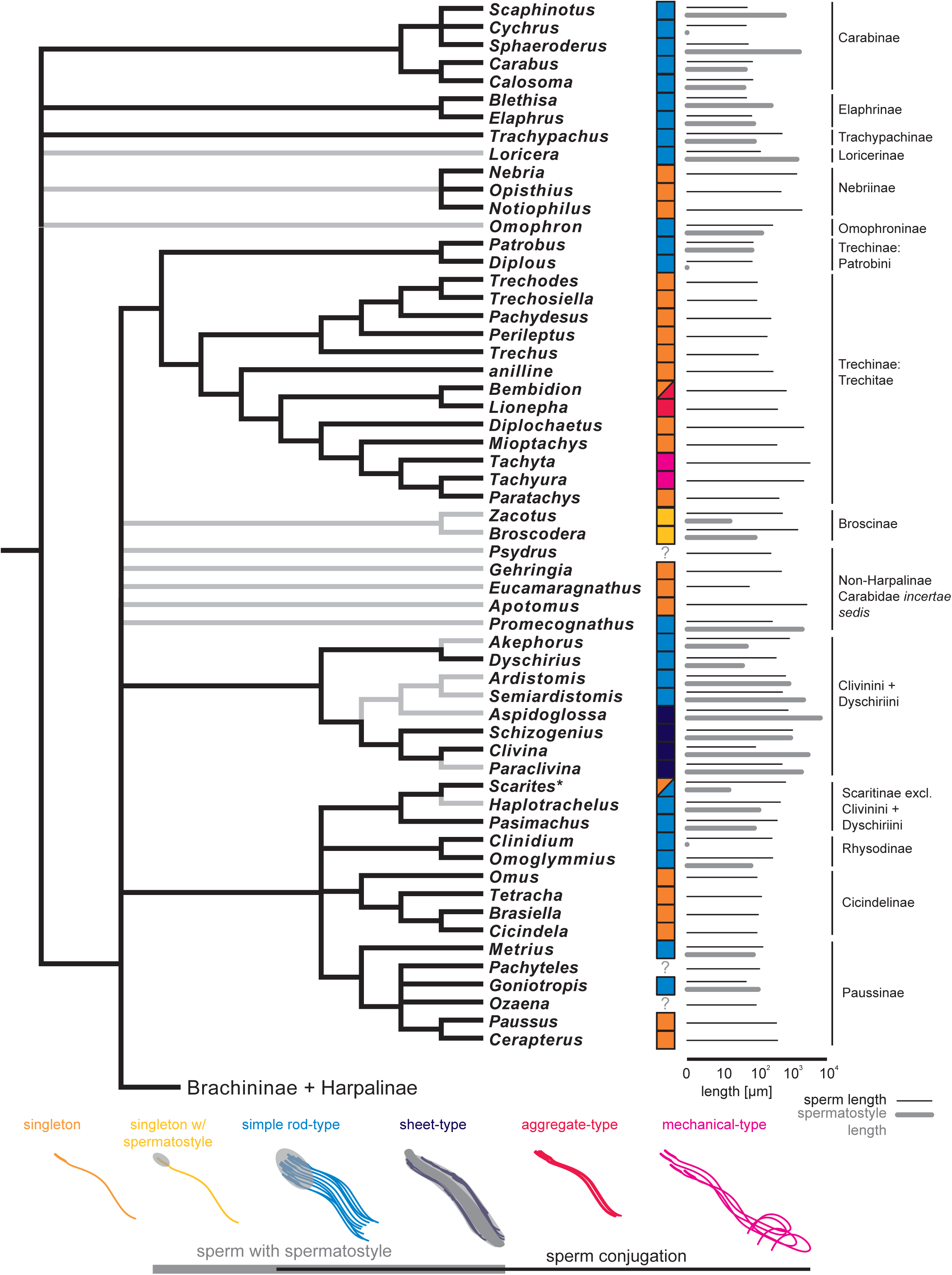
A genus-level phylogenetic visualization of ground beetle sperm data. Colored boxes refer to different qualitative types of sperm conjugation, which are described in the text (see methods) and illustrated below the tree. The tree is not derived from any one particular phylogenetic analysis but is meant to summarize current understanding of ground beetle phylogeny (see methods). Sperm length and spermatostyle length are illustrated in μm on a logarithmic scale to the right of the tree. Grey circles in place of gray bars indicate that a spermatostyle was observed but was not measured. When reporting sperm length and spermatostyle length, we chose one species per genus. In cases where we studied more than one species per genus, we chose one species arbitrarily. The asterisk beside *Scarites* refers to the fact that one species in the genus makes two distinct sperm forms, one of which is singleton and another that is involved in conjugation (Sasakawa 2009). Branches colored black in the tree are supported by molecular phylogenetic studies. Branches colored gray refer to low-resolution placements of taxa, which have not been previously sampled or whose placement is contentious

### Sperm and tissue preparation for light microscopy

Our survey largely focused on the form of mature spermatozoa of different species of carabid beetles. We dissected both males and females and extracted sperm from either the seminal vesicle of males or the sperm storage organ (spermatheca) of females, respectively. Our sampling (Table 1) is biased towards male beetles because aspects of our sampling were largely opportunistic, the probability of collecting mature sperm is high in males whereas in females it requires their having been inseminated, and we found consistent evidence that sperm, particularly sperm conjugates, undergo changes in the female reproductive tract, posing challenges for documenting sperm form prior to their exposure to the environment of the female reproductive tract.

Our sperm preparation methods largely followed those of Higginson et al., (2012a; 2015). We removed the external and internal genitalia from live beetles or, rarely, beetles preserved in 10% neutral buffered formalin, and placed them in a small drop of IX Phosphate Buffered Saline (PBS) prior to further dissection. For small­bodied beetles (5mm and smaller), we frequently removed the entire abdomen and placed it in 1X PBS prior to isolating portions of the reproductive tract and collecting sperm. We reassociated dissected tissues with the specimen either by placing them on the slide alongside the sperm or, more commonly, by placing them in a micro vial with glycerin stored beneath the pinned specimen.

For males, we first isolated the accessory glands and testes from the aedeagus. We then used an insect pin and fine forceps to gently loosen the testes and male internal tract. We identified the vas deferens where it meets the accessory glands and severed a portion of it. We transferred the severed portion of the male’s tract to a small drop of 1X PBS on a clean gelatin-coated or charged slide. We gently shook the tissue to release sperm into the saline or held the tissue with forceps and ran fine scissors along the length of the tract to extract sperm. For females, we generally isolated the spermatheca and its subtending duct from the bursa copulatrix (Liebherr and Will, 1998). We transferred the severed tract to a drop of 1X PBS on a clean subbed slide. We gently shook the tissue to release sperm or made a longitudinal incision along the outer wall of the spermatheca and compressed it to release stored sperm. After collecting sperm, the slides were allowed to air dry and were stored in slide boxes prior to fixation, staining, and mounting.

The majority of our sampling is based on sperm preparations made using a portion of the male’s seminal vesicle or female’s spermatheca. In a few cases, however, we also made observations from slides of testes, additional female reproductive tract structures, and spermatophores by placing the tissues or spermatophores in saline on a subbed slide and allowing the slide to air dry.

Once dry, we fixed and stained the slides using two different protocols. For 21 of our earliest samples (up to specimen RAGspcmn0000000134), we simultaneously fixed and stained sperm using SpermBlue and the manufacturer’s standard protocol (van der Horst and Maree, 2010) followed by mounting in Euparal. Sperm heads in carabid beetle sperm were not easily visible with SpermBlue and brightfield microscopy, and we switched to viewing heads using DAPI and fluorescence. For DAPI staining, we first placed slides in Coplin jars with a 3:1 mixture of methanol and acetic acid for 1 minute. After fixation, we rinsed the slides in 1X PBS for 1 minute and then removed the slides from buffer to dry briefly. Once partly dry, we placed a 2μl drop of Pro Long Diamond Antifade Mountant with DAPI on top of our sample along with a clean cover slip and left the mountant to cure for at least 24 hours.

### Light microscopy, imaging, and image analysis

For eight species we recorded videos of live sperm *in vitro* using a Leica Z6 lens and JVC KY-F75U camera in conjuction with Microvision’s Archimed software. Sperm were removed from beetles using our standard dissection procedure and placed on a slide in IX PBS under a coverslip. We recorded a total of 16 short movies of live ground beetle sperm conjugates from eight species at ambient temperature (Supporting Information MV1-MV16J. Although the videos are low quality, they give a coarse-grained view of how the conjugates of these species move and perhaps insight into how morphologically similar conjugates might move.

We visualized dead sperm using brightfield, darkfield, and fluorescence microscopy and differential inference contrast (DICJ on a Leica DM5500 compound microscope. We observed sperm and sperm conjugates at magnifications ranging between 100-400x depending on the size of the subject. Sperm heads were most easily visualized with fluorescence at 1000x as they are regularly about 1μm in length.

We used a Leica C425 camera paired with the Leica LAS software package to image our samples on a Leica DM5500 microscope. We chose to photograph sperm and sperm conjugates that were relatively isolated, in good condition, and easy to image or measure. We took a variable number of photographs per specimen and/or sperm preparation depending on the complexity and size of the subject matter, the quality of the preparation, and the sex of the beetle. For instance, sperm longer than 1mm frequently required taking more than one photo and stitching them together afterwards to fully capture the entire cell in a single image. We attempted to image at least five individual sperm cells and at least five sperm conjugates per preparation. We did not take measurements of sperm conjugates from females as the conjugates are modified by the female’s reproductive tract. We made qualitative observations of sperm conjugates from our female preparations and categorized conjugates by type.

We gathered morphometric data on sperm morphological variation in carabid beetles from these photographs using ImageJ (Rasband, 2012). We recorded data on the physical dimensions of individual sperm and the resulting conjugate when present. We studied 5.4 sperm on average per preparation across all preparations (n=397J. Sperm conjugation was observed in 147 of 177 species, and we studied 4.56 sperm conjugates on average per preparation among the 212 male preparations with conjugates. We gathered linear morphometric data for the following six traits: sperm length, head length, head width, spermatostyle length, spermatostyle width, conjugate length, and the length of the spermatostyle that is bare apically. We also directly counted or estimated the number of sperm found within a given conjugate. Carabid beetle sperm conjugates can frequently include hundreds to thousands of sperm, and when a direct count of sperm number was not an option, we used ImageJ and calculated the corrected total cell fluorescence of sperm heads or mitochondrial derivatives following McCloy et al., (2014) to estimate the number of sperm in a conjugate from our DAPI-stained sperm preparations.

We investigated the precision of our instruments and workflow in order to determine the number of significant figures we can reliably report in our measurements. To do this, we repeatedly measured identical subjects from photographs obtained from several rounds of imaging and microscope recalibration at different magnifications. Results from our test showed that we could reliably measure quantitative sperm traits down to two significant figures. For example, our instruments and workflow were precise to the nearest 1μm when measuring sperm heads between 10-20μm in length and were precise to the nearest 0.1 μm for sperm heads about lμm or less in width. Based on the results of our investigation, we rounded off our measurements to two significant figures.

### Data accessibility

Specimens dissected for this study and all resulting slides are stored in the personal research collection of RA Gomez and are available for examination upon request. The 6,499 light microscope images we captured are all available online through Morphobank (at http://morphobank.org/permalink/7P3123) and are organized by species and specimen code. We took a dorsal habitus photograph of one specimen of each species we studied for this project excluding *Pseudaptinus tenuicollis.* These photos are available online through Morphobank.

## 4. Results

### Overview of sperm form in carabid beetles

Our dataset includes new sperm data for 177 species of carabid beetle from throughout the group’s taxonomic breadth (Fig. 2) and reveals notable variation in ground beetle sperm, sperm conjugates, and sperm storage (Fig. 5-12). These data are summarized by species in Table 2 and by specimen in Supporting Information Spreadsheet S1. In advance of presenting taxon-by-taxon results (next section), we provide here an overview of our findings.

**Table 2.** Summarized sperm morphological data for 177 species of ground beetles. In order to limit column width, the term rod is used in place of spermatostyle. All measurements are reported in microns (μm) excluding the last column, which lists the average number of sperm found in a conjugate.

| Taxon | Conjugate type | Specimens examined (n) | Sperm length (avg) | Head length (avg) | Head width (avg) | Rod length (avg) | Rod width (avg) | Rod bare apex length (avg) | Sperm in conjugate (avg) |
| --- | --- | --- | --- | --- | --- | --- | --- | --- | --- |
| *Abacetus* sp. | Sheet | 4 | 590 | 1.5 | 0.3 | 2300 | 45 | 490 | 200 |
| *Abaris splendida* | Sheet | 3 | 170 | 1.5 | 0.4 | 190 | 7.9 | 12 | 24 |
| *Agonum muelleri* | Simple rod | 2 | 240 | 1.2 | 0.6 | 220 | 10 | 6.2 | 51 |
| *Agonum piceolum* | Simple rod | 5 | 260 |  |  | 160 | 8.1 | 16 | 66 |
| *Agra* sp. 1 |  | 1 | 74 |  |  |  |  |  |  |
| *Agra* sp. 2 | Simple rod | 1 | 140 |  |  | 90 | 5.2 | 19 |  |
| *Akephorus obesus* | Simple rod | 5 | 860 | 2.9 | 6.3 | 53 | 6.9 | 1.0 | 7.2 |
| *Amara aenea* | Simple rod | 8 | 940 | 1.4 | 0.4 | 1600 | 7.8 | 3.3 | 190 |
| *Amara farcta* |  | 1 | 540 | 2.4 | 0.5 |  |  |  |  |
| *Anatrichis minuta* |  | 1 | 720 | 4.3 | 0.5 |  |  |  |  |
| *Anillini* gen. nov. sp. nov. | Singleton | 4 | 290 |  |  |  |  |  |  |
| *Anisodactylus alternans* | Simple rod | 3 | 410 | 2.2 | 0.4 | 3700 | 21 | 60 | 120 |
| *Anisodactylus anthracinus* | Simple rod | 2 | 730 | 2.3 | 0.4 | 3900 | 17 | 94 | 880 |
| *Anisodactylus similis* | Simple rod | 1 | 380 |  |  | 3300 | 19 |  | 120 |
| *Anthia (Termophilum)* sp. |  | 1 | 1200 |  |  | 5800 | 13 |  |  |
| *Apenes lucidula* | Simple rod | 1 | 200 | 2.3 | 0.5 | 470 | 4.6 | 12 | 31 |
| *Apotomus* sp. | Singleton | 4 | 2700 |  |  |  |  |  |  |
| *Ardistomis obliquata* | Simple rod | 3 | 990 |  |  | 1400 | 21 |  | 220 |
| *Ardistomis schaumii* | Simple rod | 5 | 660 |  |  | 840 | 11 | 72 | 77 |
| *Aspidoglossa subangulata* | Simple rod | 2 | 790 |  |  | 6600 | 29 | 190 | 160 |
| *Badister ferrugineus* | Simple rod | 1 | 660 | 3.5 | 0.3 | 3800 | 19 |  |  |
| *Bembidion incrematum* | Singleton | 3 | 300 |  |  |  |  |  |  |
| *Bembidion iridescent* | Aggregate | 2 | 290 |  |  |  |  |  |  |
| *Bembidion kuprianovi* #2 | Singleton | 1 | 520 |  |  |  |  |  |  |
| *Bembidion obliquulum* | Singleton | 2 | 860 |  |  |  |  |  |  |
| *Bembidion sejunctum* | Aggregate | 1 | 760 |  |  |  |  |  | 6.8 |
| *Bembidion* sp.nr. *transversale* | Singleton/<br>Aggregate | 4 | 710 |  |  |  |  |  | 6.9 |
| *Bembidion zephyrum* | Singleton | 3 | 420 |  |  |  |  |  |  |
| *Blethisa oregonensis* | Simple rod | 2 | 51 | 11 | 0.8 | 260 | 11 | 9 | 220 |
| *Brachinus elongatulus* | Simple rod | 5 | 130 | 0.8 | 0.3 | 17 /<br>350 | 5.2 /<br>5.1 | 9.8 / ? | 51 / ? |
| *Brachinus ichabodopsis* | Simple rod | 1 | 120 |  |  | 640 |  |  |  |
| *Bradycellus* sp. 1 | Simple rod | 1 | 170 | 0.9 | 0.4 | 4900 | 11 |  | 900 |
| *Bradycellus* sp. 2 | Simple rod | 1 | 130 | 1.2 | 0.4 | 2200 | 8.9 | 72 | 2100 |
| *Brasiella wickhami* | Singleton | 1 | 110 |  |  |  |  |  |  |
| *Broscodera insignis* | Singleton<br>with rod | 2 | 1500 | 73 | 0.6 | 92 | 1.8 | 11 |  |
| *Calathus peropacus* | Simple rod | 3 | 210 | 2.4 | 0.5 | 3800 | 24 | 130 | 2900 |
| *Calleida bella* | Simple rod | 1 | 190 |  |  | 77 | 4.1 | 1.9 |  |
| *Calleida decora* | Sheet | 1 | 160 | 2.1 | 0.4 |  |  |  |  |
| *Calleida jansonii* | Sheet | 2 | 220 |  |  | 230 | 8.8 |  |  |
| *Calosoma peregrinator* | Simple rod | 2 | 78 | 18 | 0.8 | 44 | 11 | 2.9 | 48 |
| *Carabus nemoralis* | Simple rod | 1 | 72 | 11 | 0.6 | 46 | 17 | 26 | 47 |
| *Carabus taedatus* | Simple rod | 1 | 81 | 11 | 0.7 | 51 | 14 | 4.7 | 120 |
| *Catapiesis* sp. |  | 1 | 280 | 1.0 | 0.3 |  |  |  |  |
| *Cerapterus* sp. | Singleton | 1 | 390 | 26 | 0.8 |  |  |  |  |
| *Chlaenius cumatilis* | Simple rod | 2 | 480 | 1.7 | 0.4 | 660 | 6.3 | 15 | 77 |
| *Chlaenius glaucus* | Simple rod | 1 | 1200 | 2.9 | 0.4 | 910 | 8.5 | 82 |  |
| *Chlaenius harpalinus* |  | 2 | 510 | 3.7 | 0.5 | 490 | 5.3 |  |  |
| *Chlaenius leucoscelis* | Simple rod | 2 | 340 | 3.0 | 0.5 | 49 | 13 | 4.3 | 81 |
| *Chlaenius prasinus* | Simple rod | 1 | 360 | 3.0 | 0.4 | 17 | 8.6 | 3.2 | 56 |
| *Chlaenius ruficauda* | Sheet | 2 | 1000 | 6.7 | 0.4 | 5000 | 8.1 | 79 |  |
| *Chlaenius sericeus* | Simple rod | 1 | 330 | 1.4 | 0.5 | 450 | 5.1 | 22 | 130 |
| *Chlaenius tricolor* | Simple rod | 1 | 250 | 3.5 | 0.7 | 250 | 3.9 | 13 |  |
| *Cicindela haemorrhagica* | Singleton | 3 | 97 |  |  |  |  |  |  |
| *Clinidium* sp. nr. *guatemalenum* | Simple rod | 2 | 270 | 16 | 0.5 |  |  |  |  |
| *Clivina fossor* | Sheet | 3 | 91 |  |  | 3100 | 37 | 2600 | 370 |
| *Colliuris pensylvanica* | Simple rod | 3 | 330 | 2.1 | 0.4 | 840 | 18 | 8.1 | 340 |
| *Cychrus tuberculatus* | Simple rod | 1 | 50 | 9.1 | 0.7 |  |  |  |  |
| *Cycloloba* sp. |  | 1 | 260 |  |  | 4400 | 15 |  |  |
| *Cyclotrachelus dejeanellus* | Sheet | 3 | 480 | 4.3 | 0.4 | 6200 | 10 | 130 |  |
| *Cymindis basipunctata*-group sp. | Sheet | 1 | 240 |  |  | 800 | 3.5 |  |  |
| *Cymindis punctifera* | Simple rod | 1 | 240 | 1.6 | 0.3 | 1100 | 16 | 6.1 | 85 |
| *Cymindis punctigera* | Simple rod | 1 | 210 | 1.6 | 0.3 | 550 | 10 | 59 | 130 |
| *Cyrtomoscelis* cf. *dwesana* | Sheet | 3 | 700 |  |  |  | 20 |  |  |
| *Dicaelus suffusus* | Simple rod | 3 | 490 |  |  |  |  |  |  |
| *Diplochaetus planatus* | Singleton | 5 | 2200 | 46 | 0.4 |  |  |  |  |
| *Diplocheila nupera* | Simple rod | 1 | 570 |  |  | 13000 | 62 |  |  |
| *Diplous filicornis* | Simple rod | 1 | 73 | 1.8 | 0.6 |  |  |  |  |
| *Discoderus* sp. | Sheet | 1 | 300 | 1.8 | 0.4 | 860 | 6.9 | 3.8 | 34 |
| *Disphaericus* sp. | Simple rod | 2 | 790 |  |  | 600 | 11 |  | 54 |
| *Drypta* sp. |  | 3 | 180 |  |  |  |  |  |  |
| *Dyschirius dejeanii* | Simple rod | 1 | 790 | 3.4 | 1.3 | 1000 | 4.6 | 7.9 | 7.3 |
| *Dyschirius globosus* | Simple rod | 6 | 320 | 15 | 0.5 |  | 3.5 | 21 | 20 |
| *Dyschirius haemorrhoidalis* | Simple rod | 1 | 550 | 2.0 | 1.9 | 280 | 4 | 13 | 7.7 |
| *Dyschirius pacificus* | Simple rod | 2 | 990 | 3.5 | 1.1 |  |  |  | 7.2 |
| *Dyschirius thoracicus* | Simple rod | 4 | 360 | 4 | 4.3 | 40 | 7 | 3 | 7.5 |
| *Dyschirius tridentatus* | Simple rod | 1 | 820 | 4.0 | 4.5 | 1000 | 8.1 | 34 | 35 |
| *Ega sallei* |  | 1 | 1100 |  |  |  |  |  |  |
| *Elaphrus purpurans* | Simple rod | 5 | 73 | 13 | 0.9 | 83 | 4.9 | 3.5 | 54 |
| *Eucamaragnathus oxygonus* | Singleton | 3 | 62 | 8.5 | 1.7 |  |  |  |  |
| *Euryderus grossus* | Simple rod | 2 | 630 | 2.1 | 0.4 | 2900 | 13 | 84 | 1100 |
| *Galerita atripes* |  | 1 | 260 |  |  |  |  |  |  |
| *Galerita bicolor* |  | 1 | 300 |  |  |  |  |  |  |
| *Galerita forreri* | Sheet | 1 | 340 |  |  | 4800 | 41 | 76 |  |
| *Galerita lecontei* | Sheet | 2 | 280 |  |  | 3500 | 26 | 56 |  |
| *Gehringia olympica* | Singleton | 2 | 500 |  |  |  |  |  |  |
| *Goniotropis parca* | Simple rod | 3 | 48 | 12 | 0.6 | 100 | 9.2 | 5.3 | 94 |
| *Graphipterus* sp. | Simple rod | 1 | 960 | 0.5 | 0.4 | 2200 | 20 |  |  |
| *Haplotrachelus atropsis* | Simple rod | 1 | 470 | 0.6 | 0.2 | 120 | 13 |  | 96 |
| *Haplotrachelus cf. latesulcatus* | Simple rod | 2 | 420 | 0.8 | 0.3 | 180 | 18 | 23 | 130 |
| *Haplotrachelus sp.* | Simple rod | 2 | 410 | 0.9 | 0.3 | 140 | 16 | 11 | 75 |
| *Harpalus affinis* | Simple rod | 4 | 730 |  |  | 1800 | 12 | 510 |  |
| *Helluomorphoides papago* |  | 2 | 450 |  |  |  |  |  |  |
| *Hybothecus flohri* |  | 1 | 360 | 2.0 | 0.4 |  |  |  |  |
| *Lachnophorus elegantulus* | Simple rod | 2 | 290 |  |  | 710 | 22 | 25 | 130 |
| *Lachnophorus sp.* | Sheet | 1 | 290 |  |  | 610 | 13 |  |  |
| *Lebia deceptrix* | Sheet | 2 | 340 | 1.2 | 0.3 | 430 | 5.2 | 4.5 |  |
| *Lebia subgrandis* | Sheet | 1 | 370 | 2.0 | 0.3 | 520 | 6.5 | 3.2 |  |
| *Lebia viridis* | Simple rod | 2 | 340 | 0.8 | 0.4 | 800 | 14 | 31 |  |
| *Leptotrachelus sp.* | Sheet | 1 | 240 | 2.1 | 0.4 | 1100 | 63 | 120 |  |
| *Lionepha sp. nov.* | Aggregate | 3 | 400 |  |  |  |  |  | 6.7 |
| *Loricera decempunctata* | Simple rod | 3 | 130 | 12 | 0.9 | 1400 | 5.5 | 8.7 | 1500 |
| *Loricera foveata* | Simple rod | 2 | 120 | 13 | 0.9 |  |  |  |  |
| *Macrocheilus sp.* |  | 1 | 510 |  |  | 1400 | 8.0 |  |  |
| *Mastax sp.* | Simple rod | 4 | 170 | 0.8 | 0.3 | 20 | 2.7 | 3.5 | 21 |
| *Metrius contractus* | Simple rod | 3 | 150 | 7.8 | 1.0 | 84 | 7.2 | 5.0 | 220 |
| *Mioptachys flavicauda* | Singleton | 4 | 380 | 130 | 0.6 |  |  |  |  |
| *Morion sp.* |  | 1 | 140 |  |  |  |  |  |  |
| *Nebria brevicollis* | Singleton | 1 | 1400 | 26 | 0.6 |  |  |  |  |
| *Notiophilus sylvaticus* | Singleton | 3 | 2000 | 43 | 0.5 |  |  |  |  |
| *Omoglymmius hamatus* | Simple rod | 3 | 290 | 44 | 0.6 | 69 | 9.1 | 6.2 | 66 |
| *Omophron americanum* | Simple rod | 1 | 750 | 4.1 | 2.3 | 870 | 6.2 | 700 | 130 |
| *Omophron ovale* | Simple rod | 4 | 280 | 6.5 | 2.9 | 140 | 6.4 | 74 | 40 |
| *Omus audouini* | Singleton | 7 | 120 |  |  |  |  |  |  |
| *Omus dejeanii* | Singleton | 3 | 130 |  |  |  |  |  |  |
| *Oodes fluvialis* | Simple rod | 8 | 880 | 1.3 | 0.4 | 520 | 8.3 |  |  |
| *Opisthius richardsoni* | Singleton | 2 | 500 | 13 | 0.9 |  |  |  |  |
| *Ozaena* sp. |  | 1 | 86 |  |  |  |  |  |  |
| *Pachydesus* sp. | Singleton | 4 | 250 | 1.3 | 0.4 |  |  |  |  |
| *Pachyteles* sp. |  | 1 | 130 |  |  |  |  |  |  |
| *Panagaeus sallei* | Simple rod | 3 | 1400 | 6.7 | 0.5 | 1100 | 20 | 430 | 610 |
| *Paraclivina bipustulata* | Sheet | 2 | 540 |  |  | 1900 | 64 | 1100 | 370 |
| *Paratachys* sp. 1 | Singleton | 1 | 290 |  |  |  |  |  |  |
| *Paratachys* sp. 2 | Singleton | 2 | 450 | 29 | 0.5 |  |  |  |  |
| *Pasimachus californicus* | Simple rod | 3 | 390 | 0.8 | 0.3 | 86 | 16 | 24 | 66 |
| *Patrobus longicornis* | Simple rod | 4 | 76 | 1.3 | 0.3 | 74 | 4.8 | 0.6 | 23 |
| *Paussus (Bathypaussus)* sp. | Singleton | 1 | 420 | 22 | 0.5 |  |  |  |  |
| *Paussus cucullatus* | Singleton | 2 | 370 | 31 | 0.7 |  |  |  |  |
| *Pentagonica* sp. | Simple rod | 2 | 260 | 1.1 | 0.3 | 250 | 5.4 | 41 | 26 |
| *Perigona nigriceps* | Simple rod | 1 | 150 |  |  | 88 | 3.7 | 14 | 33 |
| *Perileptus* sp. | Singleton | 2 | 200 | 0.9 | 0.4 |  |  |  |  |
| *Pheropsophus* sp. 1 | Simple rod | 5 | 130 | 1.0 | 0.3 | 41000 | 7.8 | 8.5 |  |
| *Pheropsophus* sp. 2 | Simple rod | 1 | 120 |  |  |  | 5.2 |  |  |
| *Phloeoxena nigricollis* | Simple rod | 1 | 710 |  |  | 21 | 3.1 | 12 |  |
| *Poecilus laetulus* | Simple rod | 2 | 420 | 3.9 | 0.5 | 6100 | 140 |  |  |
| *Poecilus scitulus* | Simple rod | 1 | 420 | 2.2 | 0.4 |  |  |  |  |
| *Polpochila erro* | Simple rod | 2 | 320 |  |  | 810 | 34 | 100 |  |
| *Promecognathus laevisissimus* | Simple rod | 2 | 290 | 21 | 0.7 | 2000 | 35 | 58 | 420 |
| *Pseudaptinus horni* |  | 1 | 390 |  |  |  |  |  |  |
| *Pseudaptinus simplex* |  | 1 | 450 |  |  |  |  |  |  |
| *Pseudaptinus tenuicollis* |  | 1 | 400 |  |  | 1600 | 5.7 |  |  |
| *Pseudomorpha* sp. |  | 1 | 200 |  |  |  |  |  |  |
| *Psydrus piceus* |  | 1 | 260 | 13 | 0.8 |  |  |  |  |
| *Pterostichus infernalis* | Sheet | 5 | 330 | 2.6 | 0.4 | 2100 | 11 | 100 | 87 |
| *Pterostichus lama* | Sheet | 1 | 340 | 2.2 | 0.4 | 9600 | 27 | 110 |  |
| *Pterostichus melanarius* | Sheet | 1 |  |  |  |  |  |  |  |
| *Rhadine dissecta*-group sp. | Simple rod | 1 | 350 | 1.7 | 0.6 | 450 | 12 | 4.2 | 280 |
| *Scaphinotus marginatus* | Simple rod | 2 | 54 | 11 | 0.7 | 620 | 14 | 2.3 | 1500 |
| *Scarites (Distichus)* sp. | Simple rod | 1 | 260 | 1.0 | 0.4 | 160 | 3.9 |  | 19 |
| *Scarites (Parallelomorphus)* sp. | Simple rod | 2 | 330 |  |  | 40 | 7.1 |  | 64 |
| *Scarites marinus* | Simple rod | 3 | 200 |  |  | 18 | 3.2 | 10 | 17 |
| *Schizogenius litigiosus* | Sheet | 3 | 1000 |  |  | 960 | 4.3 | 44 | 7.2 |
| *Selenophorus* sp. |  | 1 | 260 | 2.3 | 0.4 |  |  |  |  |
| *Semiardistomis viridis* | Simple rod | 6 | 540 |  |  | 2200 | 3.1 | 150 | 42 |
| *Sericoda bembidioides* | Simple rod | 1 | 250 |  |  |  |  |  |  |
| *Sphaeroderus schaumii* | Simple rod | 2 | 61 | 15 | 0.7 | 1400 | 6.4 | 2.7 | 1100 |
| *Sphaeroderus stenostomus* | Simple rod | 5 | 57 | 16 | 0.6 | 1700 | 8.2 | 3.4 | 1500 |
| *Stenocrepis elegans* | Simple rod | 1 | 400 |  |  | 1900 | 31 |  | 1000 |
| *Stenognathus quadricollis* |  | 1 | 550 |  |  |  |  |  |  |
| *Stenolophus* sp. | Sheet | 1 | 500 | 1.4 | 0.3 | 5300 | 25 | 60 |  |
| *Stenomorphus convexior* | Sheet | 3 | 490 | 3.8 | 0.4 | 760 | 6.5 | 4.4 |  |
| *Stolonis intercepta* | Simple rod | 2 | 520 | 12 | 0.5 | 2000 | 21 | 260 | 1100 |
| *Stolonis* sp. | Simple rod | 1 | 370 |  |  | 590 | 21 |  |  |
| *Striganoviella vanhillei* | Simple rod | 6 | 410 | 2.0 | 1.9 | 220 | 4.0 | 21 | 8.3 |
| *Syntomus americanus* | Sheet | 2 | 190 | 2.1 | 0.3 | 340 | 8.5 | 35 | 55 |
| *Synuchus dubius* | Simple rod | 5 | 340 | 4.4 | 0.4 | 520 | 5.7 | 60 | 94 |
| *Tachyta inornata* | Mechanical | 2 | 3400 | 270 | 0.5 |  |  |  |  |
| *Tachyura rapax* | Mechanical | 1 | 2200 | 82 | 0.8 |  |  |  |  |
| *Tetracha carolina* | Singleton | 3 | 137 |  |  |  |  |  |  |
| *Tetragonoderus fasciatus* | Sheet | 5 | 250 | 1.2 | 0.4 | 2200 | 47 | 450 | 310 |
| *Tetragonoderus* sp. nr. *latipennis* | Sheet | 1 |  |  |  |  |  |  |  |
| *Thyreopterus flavosignatus* | Sheet | 1 | 250 | 1.5 | 0.3 | 330 | 25 | 12 | 270 |
| *Trachypachus inermis* | Simple rod | 1 | 530 | 8.5 | 1.5 | 87 | 9.4 | 12 | 12 |
| *Trachypachus slevini* | Simple rod | 4 | 840 | 12 | 1.9 |  |  |  |  |
| *Trechodes* sp. | Singleton | 1 | 100 |  |  |  |  |  |  |
| *Trechosiella scotti* | Singleton | 3 | 96 | 0.7 | 0.3 |  |  |  |  |
| *Trechus humboldti* | Singleton | 4 | 110 | 1.3 | 0.3 |  |  |  |  |
| *Zacotus matthewsii* | Singleton with rod | 4 | 550 | 18 | 0.7 | 17 | 4.3 |  |  |

New discoveries from our study include new instances of sperm conjugation, types of sperm conjugation previously unknown for the family, new occurrences of singleton sperm, newly documented sperm phenotypic variation, and the discovery that some female ground beetles store different parts of the sperm conjugate in different organs.

Sperm length frequently varies from one lineage to another, but sperm heads are almost always slender and narrow (Fig. 2B-C). Carabid beetle sperm range in total length from 48-3400μm whereas head width ranges from 0.2-6.3μm. Very few lineages of carabid beetles possess broad-headed sperm (Fig. 2C), but those that do, such as the genus *Omophron* (Fig. 6F-J) and many Dyschiriini, have very distinctive sperm that may be species-specific (Fig. 9I-N). Sperm head length varies much less than sperm total length and ranges from 0.5-270μm. Most ground beetles make sperm with heads that are shorter than 20μm in length. The sperm head is generally conspicuous as a single region of fluorescence following DAPI staining, but sperm in several lineages show two regions of fluorescence following DAPI staining: one faint region anteriorly approximately 1-2μm in length and a second prominent filament of much higher intensity fluorescence (Fig. 7B; Schubert et al., 2017). We attributed the short, faint region of fluorescence to the sperm’s nuclear DNA and the second, prominent region of fluorescence to the sperm’s mitochondrial DNA based on TEM observations of ground beetle sperm ultrastructure (Dallai et al., 2019; Witz, 1990; Gomez, unpublished data) and a recent study of carabid sperm with a similar staining pattern (Schubert et al., 2017). We considered the short, faint region of fluorescence the head because of its compact size, which is typical of heads of ground beetle sperm with this staining pattern (Schubert et al., 2017) and its location, as it is consistently located on the end of the sperm that is embedded in the spermatostyle.

Some male carabid beetles make singleton sperm, but the vast majority instead make sperm conjugates (Figs. 2-4), usually by joining variable numbers of sperm to a non-living structure called a spermatostyle. Sperm conjugation without a spermatostyle can be found in some trechite carabid beetles such as some Bembidiini, which make aggregate conjugates with few sperm (Figs. 7D-E). It is also found in some tachyine trechites such as *Tachyta inornata* and *Tachyura rapax,* whose sperm form haphazard groupings by grappling onto one another (Figs. 7-J). Singleton sperm were observed in several unrelated carabid lineages, including *Nebria* and near relatives (subfamily Nebriinae), *Gehringia olympica* (tribe Gehringiini), *Apotomus* sp. (tribe Apotomini), *Eucamaragnathus oxygonus* (tribe Hiletini), various tribes in the large clade Trechitae, tiger beetles (subfamily Cicindelinae), the ant parasite genus *Paussus* and near relatives (tribe Paussini), and the subfamily Broscinae (Fig. 3).

**Figure 4.**
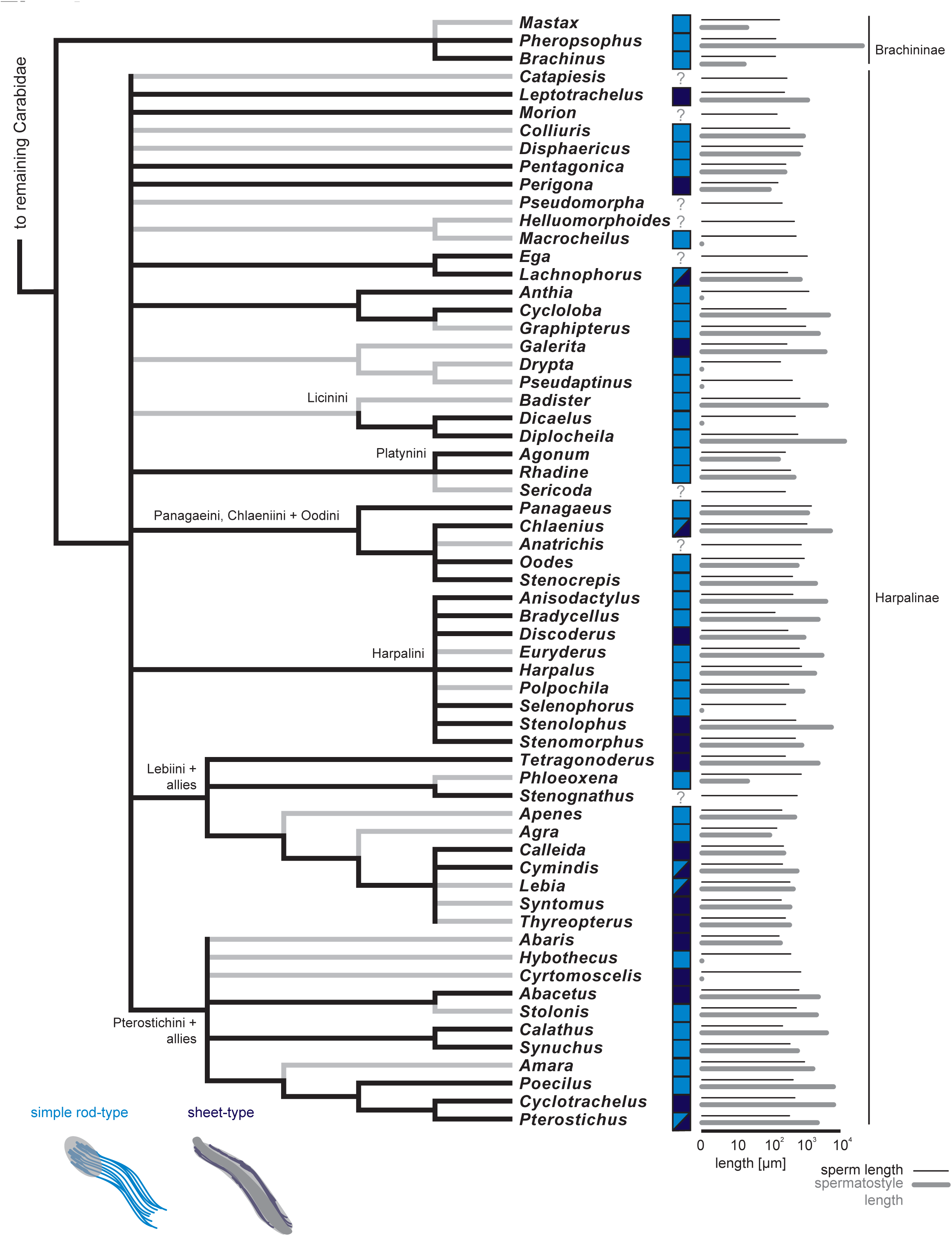
A genus-level phylogenetic visualization of ground beetle sperm data among higher-grade Carabidae (subfamilies Brachininae and Harpalinae). See Fig. 3 caption for more details.

The spermatostyle is present in nearly all ground beetles that make conjugates (Figs. 3-4). It is absent in all examined species that make singleton sperm except for two instances in the tribe Broscini indicating that sperm conjugation does not always follow spermatostyle production (Figs. 8A-C). We studied two Broscini, *Zacotus matthewsii* and *Broscodera insignis,* and both make singleton sperm joined to individual spermatostyles.

Several different aspects of the spermatostyle have been modified through evolutionary time, including size, overall shape, shape of the apex, placement or location of sperm on the spermatostyle, rigidity, thickness, and texture. Some carabid beetles make spermatostyles of varying sizes or two different size classes of similarly shaped spermatostyles (Takami and Sota, 2007). Males of the bombardier beetle *Brachinus elongatulus* make two distinct conjugates using two very different spermatostyles (Fig 11A-B).

The spermatostyle displays remarkable variation in length with relatively little variation observed in its width (Fig. 2D,E). As with sperm, the width of the spermatostyle tends to be fairly conserved across Carabidae. The widest spermatostyle we observed measured on average 140μm at its broadest. Most species, however, make spermatostyles that are much narrower in width, usually measuring between 2-20μm. The longest individual spermatostyle we observed was 5.8cm whereas the shortest individual spermatostyle we observed measured only 13μm, a more than 4,000-fold difference in length (Supporting Information Spreadsheet SI). The spermatostyle frequently varies in length between related species within major carabid lineages suggesting that spermatostyle length evolves rapidly and convergently (Figs. 2-4).

Spermatostyles are frequently rod-shaped, fusiform, or comet-shaped (broader anteriorly and attenuating to a narrow point posteriorly), but there are many exceptions. Some spermatostyles maintain this general shape but are helically shaped and rigid like a corkscrew (e.g., *Promecognathus laevissimus* spermatostyles; Fig. 8D-F) or compacted like a slinky (e.g., *Chlaenius ruficauda* spermatostyles; Fig. 11G). Others are cap-like and gelatinous (e.g., *Chlaenius prasinus* spermatostyles) or thin and ribbon-like (e.g., *Stenocrepis elegans* spermatostyles; Fig. 11I). Sperm can also be distributed along the spermatostyle in a variety of ways and in varying numbers. We recorded as few as five sperm in a conjugate to as many as a few thousand. The spermatostyle can include hyaline flanges (Figs. 6E, 8D) and channels or grooves that appear to be associated with sperm attachment or storage (Fig. 12D). The material surrounding sperm at their attachment point to the rod can have a different appearance compared to the remainder of the rod (Dallai et al., 2019; Hodgson et al., 2013). *Clivina* species make a capsule-like spermatostyle with a large sealed cavity that contains a mass of sperm (Fig. 9A). Sperm are usually densely distributed on all sides of a spermatostyle, but this trait is also variable. There are frequently extensive bare regions without attached sperm on the spermatostyles of many ground beetle conjugates (e.g., Figs. 6F, 9B, 9F, 12E). These bare regions are frequently found on the anterior end, but they commonly occur medially (Fig. 11K) or, in a few species, posteriorly. In *Aspidoglossa subangulata* sperm are attached to only one side of the spermatostyle, and although these large spermatostyles measure 6.6mm, less than 1mm of their length bear sperm (Fig. 9B). In *Dyschirius tridentatus,* the sperm are embedded in the spermatostyles via their heads in a single row with regular intervals between sperm (Figs. 9F-G). The spermatostyle commonly appears to be designed to accommodate sperm, particularly when the sperm heads are broad. Sperm will frequently be placed parallel to the longitudinal axis of the spermatostyle, but this is less common when sperm are broad-headed. For example, the broad-headed sperm of *Trachypachus* are placed within diagonally arranged slots on one side of the spermatostyle (Fig. 5G).

**Figure 5.**
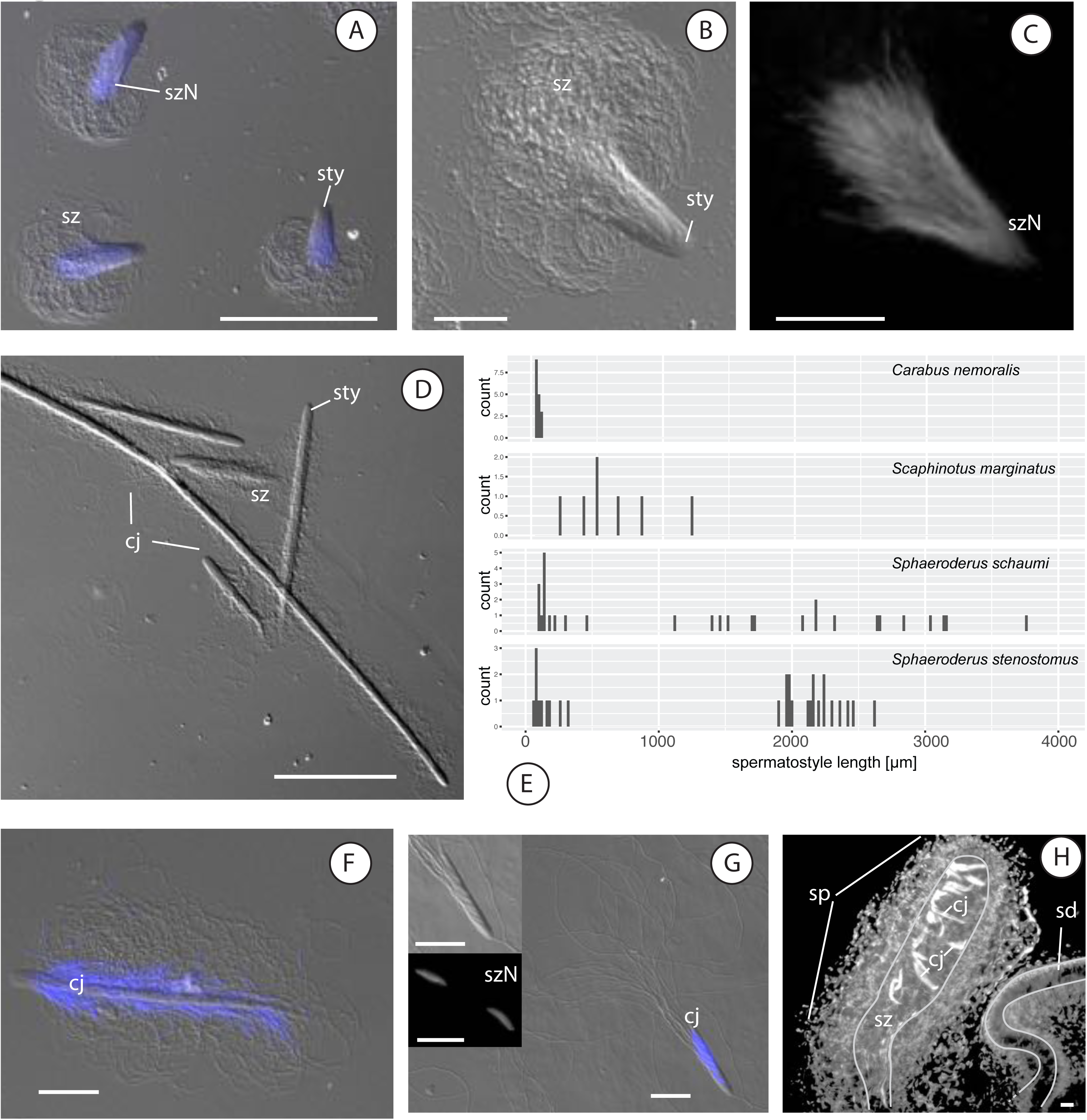
Sperm and sperm conjugate morphological variation in Carabinae (A-E), Elaphrinae (F), and Trachypachinae ground beetles (G-H). (A-C) rod conjugates of *Carabus nemoralis.* (D) rod conjugates of *Sphaeroderus stenostomus,* note conjugate size polymorphism. (E) histograms of sperm conjugate size variation in four Carabinae species. (F) rod conjugate of *Elaphrus purpurans.* (G) composite image of *Trachypachus inermis* sperm heads (lower inset) and sperm conjugates. (H) *Trachypachus slevini* female reproductive tract with stored sperm and several sperm conjugates with added thin white line to help visualize sperm storage organ and its adjoining duct. (A, F, G) stacked image of DIC and Fluoresecence microscopy images. (B, D, upper inset of G) DIC microscopy. (C, lower inset of G, H) Fluorescence images with only DAPI-stained structures visible, cj = conjugate, sd = spermathecal duct, sty = spermatostyle, sp = spermatheca, sz = spermatozoa, szN = sperm nuclei. Scale bars: 10 μm (G lower inset), 20 μm (B-C, F-H excluding lower inset of G), 100 μm (A, D).

**Figure 6.**
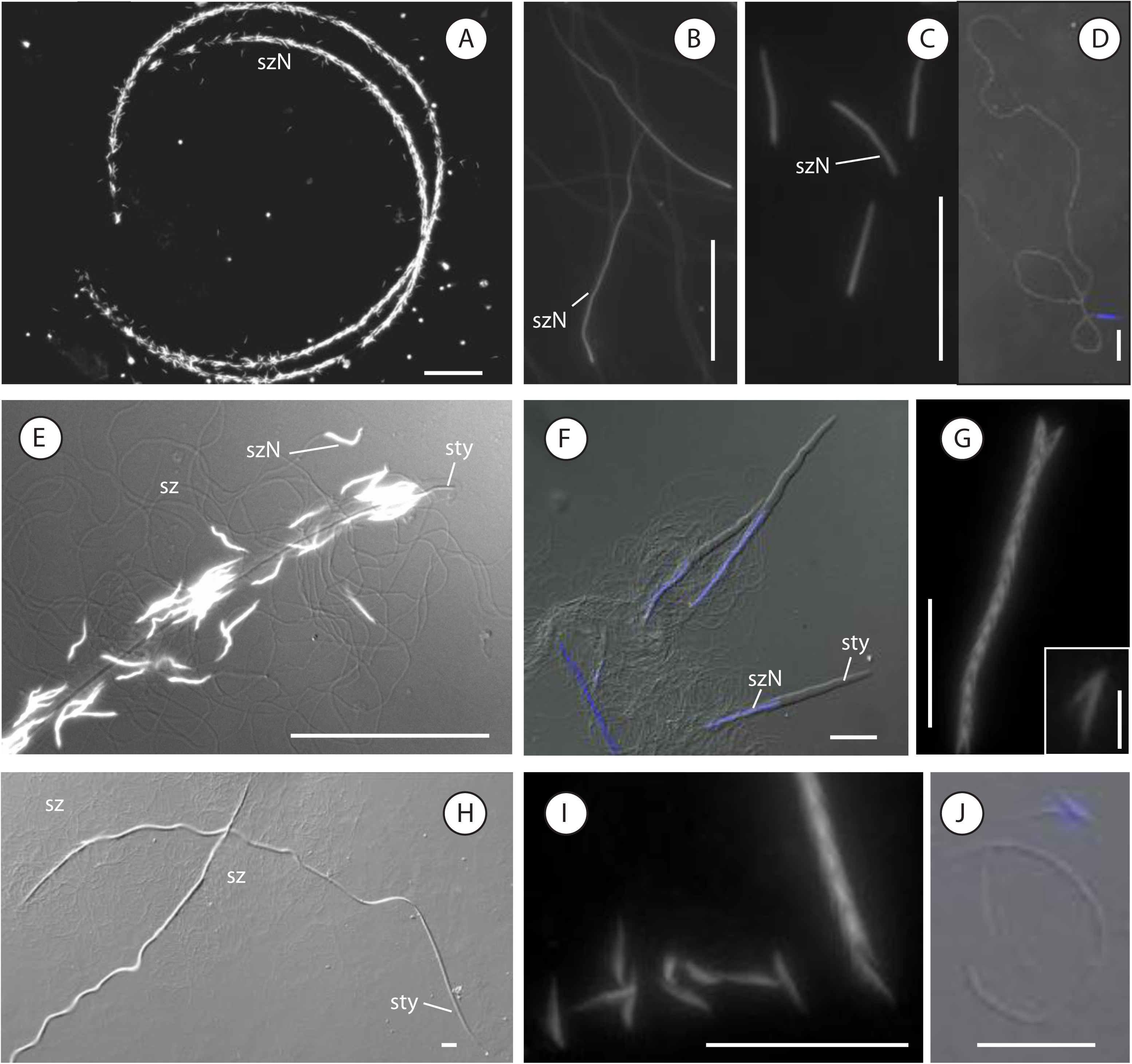
Sperm and sperm conjugate morphological variation in Loricerinae (A, E), Nebriinae (B-D), and Omophroninae (F-J) ground beetles. (A) large rod conjugates of *Loricera decempunctata* include approximately 1500 sperm. (B) slender and elongate sperm heads of *Notiophilus sylvaticus.* (C) sperm heads of *Opisthius richardsoni.* (D) singleton sperm of *Opisthius richardsoni.* (E) close-up of A. (F-1) complex rod conjugates of *Omophron.* (F) *Omophron ovale* rod conjugate, note the posterior placement of sperm in spermatostyle. (G) composite image of *O. ovale* sperm head and sperm conjugate, note the asymmetry of sperm heads and the stacking of heads. (H) *Omophron americanum* rod conjugate, note the prominent bare region of the spermatostyle anteriorly. (I) *O. americanum* sperm and sperm conjugate. (J) *O. americanum* spermatozoon, note the asymmetrical attachment of the flagellum. (A-C, G, I) Fluorescence images with only DAPI-stained structures visible. (D) stacked image of Darkfield and Fluoresecence microscopy images. (E-F, J) stacked image of DIC and Fluoresecence microscopy images. (H) DIC microscopy, sty = spermatostyle, sz = spermatozoa, szN = sperm nuclei. Scale bars: 5 μm (G inset), 20 μm (B-D, F-J excluding inset of G), 50 μm (E), 100 μm (A).

We dissected several female carabid beetles (Table 1), and we were usually able to recover sperm, indicating that most wild-caught females had been mated at least once. Of those preparations that were successful, we found that females stored sperm in their cul-de-sac type spermatheca and its adjoining duct. Conjugated sperm appear to become disassociated from each other in the spermatheca, and we consistently observed morphological differences in the spermatostyles recovered from our male and female slide preparations with spermatostyles recovered from female reproductive tracts appearing thinner, compressed, possibly digested, or completely absent in some of our female preparations (Figs. 12E-J). We frequently recovered some intact conjugates from our female preparations, but the spermathecae typically contained mostly individual sperm.

An unexpected sperm-female interaction from our study is the discovery that some females store different components of the male’s sperm conjugate in different storage organs (Figs. 12A-D). Females in the genus *Galerita* appear to store large quantities of spermatostyles in a balloon-shaped storage organ and individual sperm in a physically removed small spherical organ that had been thought to be glandular by Liebherr and Will, (1998). We surmise that female *Galerita* and relatives may have partially decoupled spermatostyle morphological evolution from sperm evolution by storing sperm and spermatostyles in different organs. It is clear from our study that sperm conjugation in Carabidae is widespread and variable, and females are interacting with conjugates, sperm, and spermatostyles. However, much research remains to be done to tease apart the nature and consequences of these sperm-female and conjugate-female interactions.

### Sperm form across major groups of carabid beetles

Subfamily Carabinae (Fig. 1A-E; 5A-E; 12I-J)

#### Species examined

(Table 1). Tribe Carabini: *Carabus nemoralis, Carabus taedatus,* and *Calosoma peregrinator.* Tribe Cychrini: *Cychrus tuberculatus, Scaphinotus marginatus, Sphaeroderus schaumii,* and *Sphaeroderus stenostomus*.

#### Sperm overview

The sperm of carabines tend to be among the shortest known sperm in carabid beetles (Table 2; Fig. 4A). The sperm thus far known are filamentous with slender heads that are visually indistinguishable from the rest of the cells (Fig. 5C). The sperm heads are obvious with DAPI staining (Figs. 5A, C).

All examined species of carabines make sperm conjugates with a spermatostyle (Table 2). The sperm conjugates of carabines all feature a spermatostyle that is either cap-like, short, and gelatinous in appearance (Figs. 5A-B) or rod-like, elongate, and stiff (Figs. 1A-E; 5D). Sperm are embedded in the spermatostyle via their heads although their flagella are unbounded (Figs. 1A-E). Species of *Calosoma* and *Carabus* (tribe Carabini) make sperm conjugates that are reminiscent of shuttlecocks with a short oblong spermatostyle. Longer spermatostyles with likely more sperm have been previously recorded from species of *Carabus* subgenus *Ohomoperus* (Takami and Sota, 2007). Species of *Sphaeroderus* and *Scaphinotus* (tribe Cychrini) make elongate conjugates composed of lengthy spermatostyles that include a large number of associated sperm.

#### Within-species variation

Carabines, particularly members of the genus *Carabus,* are among the best-studied carabid beetles for sperm morphology. Takami and Sota, (2007) studied several species of *Carabus* in the subgenus *Ohomopterus* and observed conjugate size polymorphism between specimens; many *Ohomopterus* make a single sperm form that is packaged into different size classes of conjugates (Takami and Sota, 2007). Takami and Sota, (2007) also found evidence for a positive correlation between risk of sperm competition and sperm conjugate polymorphism. If conjugates perform different roles depending on their size, different size classes of sperm conjugates would be expected.

Among the carabines we examined for this study, we found significant within-male conjugate size variation (Fig. 5E) in the species of Cychrini we studied, and minimal variation in *Carabus* and *Calosoma* (tribe Carabini). *Scaphinotus marginatus* and the two *Sphaeroderus* species we studied make a single short sperm morph, but males package their sperm into conjugates of different sizes (Table 2; Fig. 5E). The *Carabus* and *Calosoma* species we studied all make sperm conjugates with spermatostyles that vary less dramatically in length within males (Table 2; Fig. 5E).

#### Within-genera variation

Our sampling included two different species of the large cosmopolitan genus *Carabus,* which is split into numerous subgenera and two species of the eastern North American genus *Sphaeroderus. We* observed distinct differences in sperm length, placement of sperm in their conjugates, and number of included sperm between *C. (Tanaocarabus) taedatus* and *C. (Archicarabus) nemoralis. We* note that these species are likely not particularly closely related. Within *Carabus* subgenus *Ohomopterus,* Takami and Sota, (2007) observed variation in conjugate size polymorphism as well as minor variation in sperm length among several closely related species. Two *Sphaeroderus, S. schaumii* and *S. stenostomus,* possess morphologically similar sperm.

#### Reproductive tract observations

Male carabines tend to devote a considerable amount of intra-abdominal space to their testes and accessory glands. Perhaps because of the small size of their sperm and their generally large bodies, carabines consistently appear to make very large quantities of sperm.

We recovered partly bare spermatostyles from the spermathecae of female specimens of *Cychrus tuberculatus, Scaphinotus marginatus,* and *Sphaeroderus stenostomus* (Figs. 12I-J), confirming that the conjugates of conspecific males travel to the spermatheca.

#### Comments

Bouix, (1961; 1963) studied the sperm of several species of *Carabus* and reported finding dramatic instances of sperm polymorphism in DNA complement with some beetles having macrocephalic sperm with multiple sets of chromosomes. We did not investigate this topic systematically, but we found no instances of sperm polymorphism in DNA content in our samples. DAPI stained sperm heads within a species gave consistent fluorescent signals from cell to cell, which would not be expected if some sperm had more DNA. We suspect that Bouix, (1961; 1963) may have mistaken sperm conjugates for individual sperm cells, but this topic awaits further inquiry.

### Subfamily Elaphrinae (Figs. 5; 12G-H)

#### Species examined

(Table 1). Tribe Elaphrini: *Elaphrus purpurans* and *Blethisa oregonensis*.

#### Sperm overview

The sperm of elaphrines are short and filamentous (Table 2). The sperm heads are thin, tapered anteriorly, and are visually indistinguishable from the flagella (Fig. 5F). The heads are conspicuous with DAPI staining.

Both *Elaphrus* and *Blethisa* make sperm conjugates with moderately long rod­like spermatostyles (Table 2; Fig. 5F). The sperm are embedded in the spermatostyles via their heads with unbounded flagella, and the sperm are distributed more or less equally on all sides of the spermatostyles except for a short region anteriorly without attached sperm (Fig. 5F). The spermatostyles differ in size between *E. purpurans* and *B. oregonensis* but are similar in overall shape. The spermatostyles are narrowly rounded anteriorly and tapered posteriorly and resemble comets.

#### Within-species variation

Both male *B. oregonensis* studied showed high levels of size variation in spermatostyle length and the number of sperm in a conjugate with almost no variation between specimens in sperm size or variation in the density of sperm placement along the spermatostyle (Supporting Information Spreadsheet S1). The form of *B. oregonensis* sperm conjugates appears stable within males and within the species.

#### Reproductive tract observations

*We* recovered several largely intact conjugates from the spermatheca of one female *E. purpurans* and several completely bare spermatostyles from the spermatheca of a second female (Fig. 12G-H).

#### Sperm motility observations

The conjugates of *E. purpurans* move in the direction of the spermatosyle’s tapered slender end, which we considered posterior based on the anterior orientation of the sperm heads in the spermatostyle and histological studies of carabid conjugates (Hodgson et al., 2013; Schubert et al., 2017). It appears as though the sperm do not helically beat their flagella along their longitudinal axis but instead maintain a regular stroke pattern. The resulting movement of the conjugate is directional, similar to a rowboat (Supporting Information movies MV5-MV8). There does not appear to be any difference in swimming patterns between conjugates recovered from a female’s spermatheca and those found in a spermatophore.

### Subfamily Trachypachinae (Fig. 5G-H)

#### Species examined

(Table 1). Tribe Trachypachini: *Trachypachus inermis* and *Trachypachus slevini*.

#### Sperm overview

The sperm of *Trachypachus* is moderately long and filamentous (Table 2). Our measurements of sperm length in *T. slevini* vary somewhat across specimens (Supporting Information Spreadsheet S1) suggesting that sperm length in *T. slevini* is variable or that this variation is an artifact of our slide preparations for these samples. The sperm heads are slightly broader than the remainder of the cells (Fig. G). The heads are rod-shaped and appear narrowly rounded anteriorly (Fig. 5G).

The conjugates of *Trachypachus* are distinctive because of the asymmetrical arrangement of sperm in a conjugate, the small number of sperm in a conjugate, and the small size of the spermatostyle (Table 2; Fig. 5G). The spermatostyle is narrowly rounded anteriorly and attenuated posteriorly to a thin point. Sperm are located on only one side of the spermatostyle, and the heads are arranged diagonally relative to the longitudinal axis of the spermatostyle.

#### Within-genera variation

*Trachypachus inermis* sperm and their heads are slightly shorter than the sperm and heads of *T. slevini,* respectively.

#### Reproductive tract observations

*We* recovered two mostly complete conjugates from the spermatheca of a female *T. slevini* (Fig. 5H). The spermatostyles are asymmetrical with diagonally arranged slots and are similar to the spermatostyles of *T. inermis* sperm conjugates.

#### Sperm motility observations

*Trachypachus slevini* conjugates appear to move faster than individual sperm and seem able to change direction readily (Supporting Information MV15-MV16).

### Subfamily Loricerinae (Figs. 6A, E)

#### Species examined

(Table 1). Tribe Loricerini: *Loricera foveata* and *Loricera decempunctata*.

#### Sperm overview

*Loricera* sperm are short and filamentous (Table 2). The sperm heads are thin, tapered anteriorly, and are visually indistinguishable from the flagella (Fig. 6E). The heads are conspicuous with DAPI staining (Fig. 6E).

Gilson, (1884) first viewed the large conjugates of *Loricera* but mistook them for spermatophores. The sperm conjugates of *Loricera* include a long and thin spermatostyle with numerous sperm embedded via their heads with unbounded flagella (Fig. 6A). The sperm are distributed more or less equally on all sides of the spermatostyle along its entire length apart from a short region anteriorly (Fig. 6A). The spermatostyle is rod-like and narrows to a sharp point anteriorly and posteriorly. It is crescent-shaped and curved.

#### Within-genera variation

Sperm differ slightly in total length between *L.foveata* and *L. decempuncata,* but our data are limited.

#### Reproductive tract observations

The spermatheca of *Loricera* resembles a Gordian knot, and we found several sets of spermatostyles within the spermathecae of our specimens of *L.foveata* (Supporting Information MV9).

### Subfamily Nebriinae (Figs. 6B-D)

#### Species examined

(Table 1) Tribe Nebriini: *Nebria brevicollis.* Tribe Opisthiini: *Opisthius richardsoni.* Tribe Notiophilini: *Notiophilus sylvaticus*.

#### Sperm overview

Sperm in Nebriinae are generally long and filamentous (Table 2). Sperm heads in nebriines are either thin, tapered apically, and visually indistinct as in many other early-diverging carabid groups or rod-like and slightly broader than the remaining portions of the cells. *Opisthius richardsoni* sperm heads are rod-like and are slightly thickened (Fig. 6C-D). The sperm of *N. brevicollis* and *Notiophilus* are notable for having rather long heads that are visually indistinct when unstained (Fig. 6B).

All nebriines studied to date only make singleton sperm with no evidence of a spermatostyle (Table 2). Depending on the phylogenetic position of nebriines, this could represent an early loss of conjugation and the spermatostyle in the tree of Carabidae (Fig. 2) or singleton sperm could be the ancestral state of Carabidae.

### Subfamily Omophroninae (Figs. 6F-J)

#### Species examined

(Table 1). Tribe Omophronini: *Omophron americanum* and *Omophron ovale*.

#### Sperm overview

The sperm of *Omophron* are among the most distinctive sperm in carabid beetles (Table 2). The sperm heads are broad, asymmetrical and approximately V-shaped (Figs. 6G, I), and the flagellum joins the head asymmetrically on one of its sides (Fig. 6J).

The sperm of *Omophron* conjugates are arranged in a highly organized fashion inside of a rod-like spermatostyle (Fig. 6F-GJ). The sperm heads are paired together such that the side of the head bearing the flagellum is lateral (Fig. 6G,I). These pairs of sperm are radially stacked one inside of another in a row that is very reminiscent of the rouleaux stacking in diving beetle sperm (Higginson et al., 2012a; Pitnick et al., 2009a). Unlike the rouleaux stacking of diving beetles, the stacked grouping of *Omophron* sperm heads are embedded in a rod-like spermatostyle (Table 2). The spermatostyle is bare for approximately 80% of its length in *O. americanum* (Fig. 6H) and about 50% of its length in *O. ovale* with sperm located only in the posterior part of the conjugate (Fig. 6F).

#### Within-genera variation

Sperm are notably different between *O. americanum* and *O. ovale* with numerous morphological differences in their sperm. The sperm differ in total length, head size and shape, spermatostyle length, and the extent to which the spermatostyle lacks sperm.

#### Reproductive tract observations

*We* have been unable to recover sperm from the female reproductive tract of field-collected *Omophron* females despite at least six attempts to do so. In contrast, in other carabid genera we typically found sperm in a female’s spermatheca. *Omophron* is unusual in this regard, and we speculate that either our timing was bad or females are storing sperm in another location or using it in a non-typical way.

### Subfamily Trechinae, Tribe Patrobini (Figs. 7A-B)

#### Species examined

(Table 1). *Diplous filicornis* and *Patrobus longicornis*.

#### Sperm overview

Patrobine sperm are short and filamentous (Table 2; Figs. 7A-B). The heads are short and compact and visually indistinct. When stained with DAPI, patrobine sperm show two regions of fluorescence.

**Figure 7.**
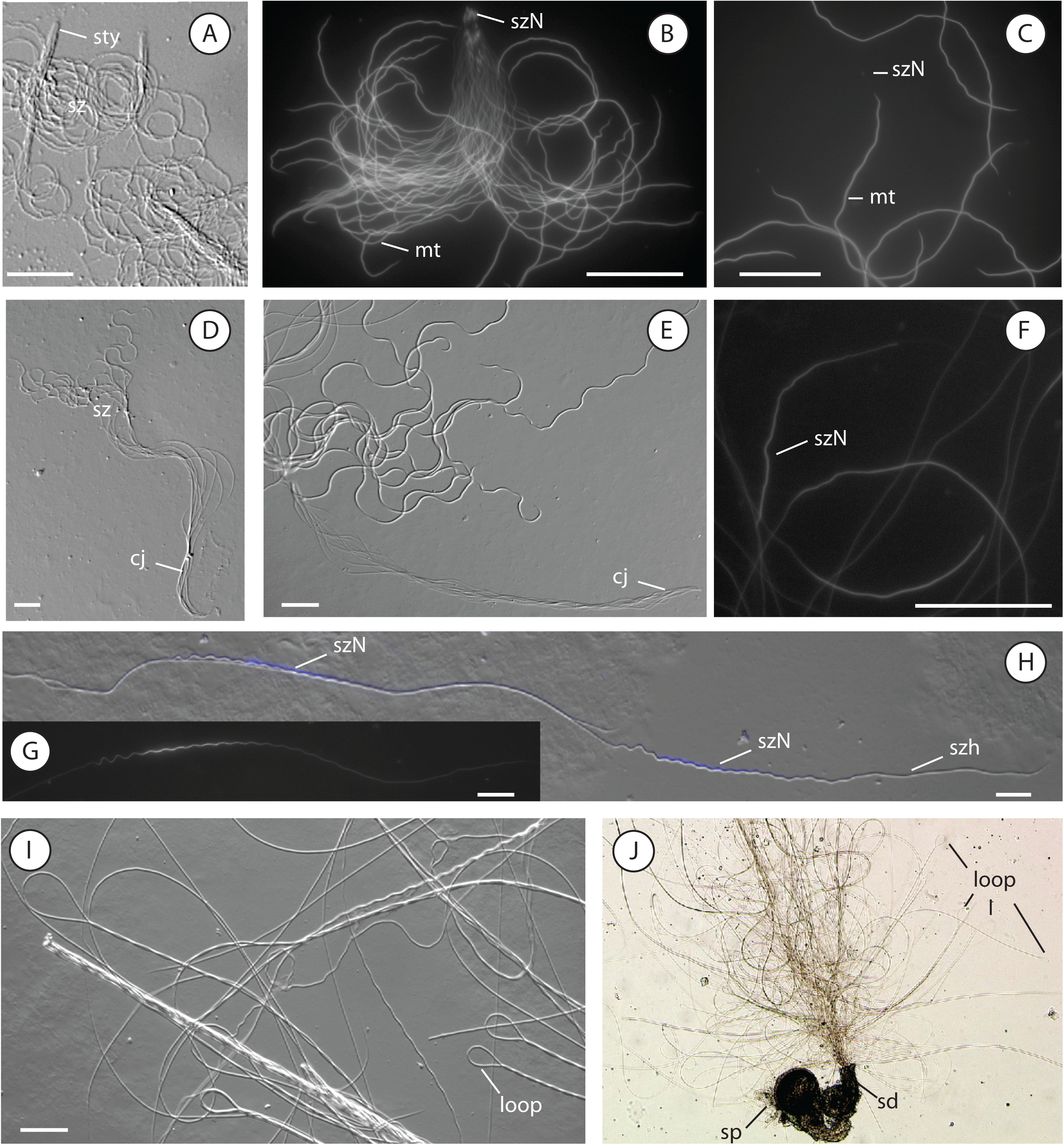
Sperm and sperm conjugate morphological variation in Trechinae ground beetles. (A) rod conjugates of *Patrobus longicornis.* (B) rod conjugate of *Diplous filicornis* recovered from the spermatheca of a female specimen. (C) *Trechus humboldti* sperm showing two regions of fluorescence corresponding to the minute nucleus and large mitochondrial derivatives. (D) aggregate conjugate with 9 sperm in an undescribed species of *Lionepha.* (E) aggregate conjugate of *Bembidion* sp. nr. *transversale,* note the lack of a spermatostyle. (F) elongate sperm heads of *Diplochaetus planatus.* (G-H) elongate sperm heads of *Tachyta inornata,* note the zig-zag shape, the extensive pre-nuclear area, and the interaction between two spermatozoa. (I) *Tachyta inornata* appear to form mechanical conjugates by forming hairpin loops with their flagella and grappling with adjacent sperm. (J) *Tachyta inornata* sperm recovered from a female spermatheca forming characteristic loops as they swim (Supporting Information MV12-MV14 of live *T. inornata* sperm). (A, D-E, H, I) DIC microscopy. (B-C, F-G) Fluorescence images with only DAPI-stained structures visible. (J) Brightfield microscopy, loop = flagellar loops, mt = mitochondrial derivatives, sd = spermathecal duct, sp = spermatheca, sty = spermatostyle, sz = spermatozoa, szh = sperm head, szN = sperm nuclei. Scale bars: 20 μm.

The sperm conjugates of patrobines include a simple rod-like spermatostyle with sperm embedded via their heads with unbounded flagella (Table 2; Figs. 7A-B). We did not gather morphometric data from the sperm conjugates recovered from our female specimen of *D.filicornis,* but it is clear that *D.filicornis* males make rod conjugates with generally 50 sperm or less embedded in a short and slender spermatostyle.

#### Reproductive tract observations

*We* recovered several seemingly intact sperm conjugates from the spermatheca of a female *D.filicornis*.

### Subfamily Trechinae, Supertribe Trechitae (Figs. 7C-J)

#### Species examined

(Table 1). Tribe Trechini, subtribe Trechodina: *Pachydesus* sp., *Perileptus* sp., *Trechodes* sp., *Trechosiella scotti.* Tribe Trechini, subtribe Trechina: *Trechus humboldti.* Tribe Anillini: an undescribed form from Oregon, USA. Tribe Bembidiini: *Bembidion incrematum, Bembidion iridescens,* one of two species under the name *Bembidion kuprianovi, Bembidion* sp. nr. *transversale, Bembidion sejunctum, Bembidion zephyrum,* and an undescribed species of *Lionepha.* Tribe Pogonini: *Diplochaetus planatus.* Tribe Tachyini: *Mioptachysflauvicauda, Paratachys* sp. 1, *Paratachys* sp. 2, *Tachyta inornata,* and *Tachyura rapax*.

#### Sperm overview

Trechitae sperm vary dramatically in length, and this variation appears to depend on conjugation state (Table 2; Fig. 2). Trechitae sperm tend to be short to very long when singletons, moderately long generally when part of a conjugate with cementing material, or very long when involved in mechanical conjugation. Sperm total length ranges from as short as 100μm in an unidentified species of *Trechodes* to the longest known sperm in Carabidae found in the tachyine *Tachyta inornata* with its 3400uin-long sperm. Sperm length across Trechitae tends to be shorter than 1mm, and sperm seem to have increased in length in the tribe Pogonini and some members of the tribe Tachyini. Sperm heads in Trechitae are generally thin, tapered anteriorly and filamentous (e.g., Fig. 7C). We were unable to consistently observe or confidently identify the heads of some trechite sperm following DAPI staining. Some trechite sperm show only one large region of fluorescence removed from either end of the sperm (e.g., *Mioptachys flauvicauda* and all studied *Bembidion* species), which we suspect corresponds to their mitochondrial derivatives (Fig. 7C). Based on the sperm heads that we could visualize, head length ranges from very short and patrobine-like in *Trechus* and trechodine trechines to long or very elongate in *Diplochaetus* and tachyines (Figs. 7G-H). The heads of *Tachyta inornata, Tachyura rapax,* and *Paratachys* spp. are unusual for their elongate size and zig-zag shape (Figs. 7G-H).

Sperm conjugation is either absent or present in Trechitae (Table 2). Singleton sperm are found in some *Bembidion,* an undescribed anilline, trechodine trechines, *Trechus humboldti, D. planatus, M. flauvicauda,* and the two *Paratachys* species we studied. The species that do make conjugates do so without an apparent spermatostyle. Conjugated sperm in the subfamily are either aggregates (Figs. 7D, E) or mechanical conjugates (Figs. 7G-J). Aggregate conjugates are found in some *Bembidion* and *Lionepha.* Within one species, *Bembidion* sp. nr. *transversale,* we had one specimen with evident aggregate conjugates, but in the other specimens we found only singleton sperm; the cause of these differences is not known (Supporting Information Spreadsheet SI). The heads of aggregate conjugates appear to be aligned in register and presumably are joined together via cementing material. Because these sperm are aligned parallel to one another and are joined together without a spermatostyle, the conjugate is approximately the same length as an individual sperm cell. Mechanically conjugated sperm were observed in *Tachyta inornata* and *Tachyura rapax,* whose sperm form conjugates haphazardly via grappling onto one another (see Sperm motility observations).

#### Within-genera variation

*We* studied several different species of the large and complex genus *Bembidion* and found that sperm differ in total length and, perhaps, presence or frequency of conjugation. Because we did not focus on a group of closely related *Bembidion,* our data cannot speak to the usefulness of sperm-level variation in species delimitation in *Bembidion*.

#### Reproductive tract observations

The spermatheca of many trechites is small, compact, and frequently well sclerotized unlike most other ground beetles (Liebherr and Will, 1998).

#### Sperm motility observations

The *Bembidion* sperm that we observed consisted of a mix of singleton sperm and conjugated sperm (Supporting Information MV3). *Bembidion* sperm move via helical klinotaxis, and their sperm conjugates swim notably faster than singleton sperm though we did not quantify this apparent difference in speed (Supporting Information MV3-MV4).

*Tachyta inornata* and *Tachyura rapax* sperm are singletons, but we observed them forming haphazard groups when released from the spermatheca or the male internal tract (Supporting Information MV12-MV14 for *T. inornata). We* observed the sperm of these beetles forming hairpin loops with their flagella (Figs. 7I-J) while undulating up and down and beating their flagella. Because of this motion and their long length, these sperm became net-like, and they began grappling onto adjacent sperm as they moved. It is difficult to fully characterize their behavior from our videos, but it appears as though sperm latch onto adjacent sperm and slide up their neighbor sperm. We observed live sperm of three male and one female *Tachyta inornata* and one male *Tachyura rapax.* Sperm in these species consistently formed hairpin loops leading to the formation of groups of sperm of varying size. Although the data are limited, we think that this is an example of secondary conjugation in Carabidae and a novel example of mechanical conjugation in animals. Mechanical conjugation is defined as a grouping of sperm that results from sperm haphazardly grappling onto one another and forming groups of variable size (Higginson and Pitnick, 2011), which is in keeping with our observations of sperm in these tachyines. Mechanical conjugation has been previously reported only from muroid rodents. Rodent sperm conjugates or trains have been the topic of much active research on the biomechanics of sperm (Fisher et al., 2014), and they possibly represent a case of sperm cooperation (e.g., Higginson and Pitnick, 2011; Immler et al., 2007; Moore, 2002; Pizzari and Foster, 2008).

### Subfamily Broscinae (Figs. 8A-C)

#### Species examined

(Table 1). *Broscodera insignis* and *Zacotus matthewsii*.

#### Sperm overview

Broscinae sperm are moderately long to long (Table 2). The sperm heads are visually indistinct from the remainder of the cell, but they are obvious with DAPI staining (Fig. 8C). The heads are filamentous and rod-like in *Z. matthewsii* (Fig. 8C) and slender, elongate, and wavy in *Broscodera insignis* (Fig. 8A).

**Figure 8.**
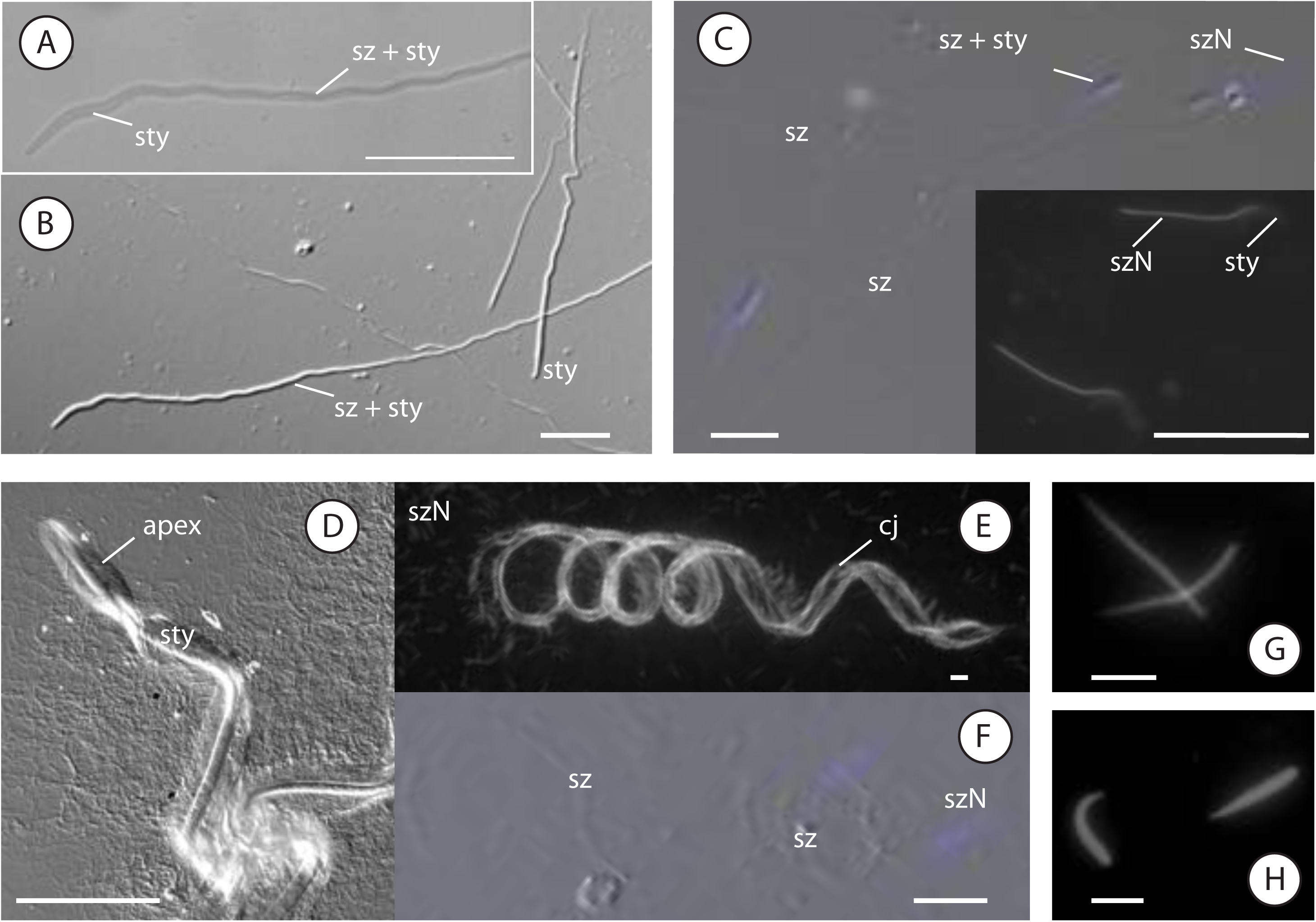
Sperm and sperm conjugate morphological variation in Broscinae (A-C) and ground beetles that we considered non-Harpalinae Carabidae of uncertain position (D-H). (A-B) *Broscodera insignis* sperm are singleton but are individually joined to a spermatostyle. (C) Composite image of *Zacotus matthewsii* sperm. Singleton sperm appear broad-headed but are filamentous and are paired with a short and broad spermatostyle (see inset). (D-E) Rod sperm conjugate of *Promecognathus laevissimus* with its large corkscrew-shaped spermatostyle. The apex of the spermatostyle is frequently variable, and in *P. laevissimus* the apex is spoon-shaped (D). (F) *Promecognathus laevissimus* spermatozoon. (G) *Psydrus piceus* sperm head. (H) *Eucamaragnathus oxygonus* sperm head. (A) Brightfield microscopy. (B, D) DIC microscopy. (C, F) stacked image of DIC and Fluoresecence microscopy images, (inset of C, E, G, H) Fluorescence images with only DAPI-stained structures visible, apex = apex of spermatostyle, cj = conjugate, sty = spermatostyle, sz = spermatozoa, szN = sperm nuclei. Scale bars: 5 μm (G, H), 20 μm (A-C, E-F), 100 μm (D).

Broscinae are notable for making a spermatostyle without conjugation (Table 2; Figs. 8A-C), which is a character combination that we have not observed outside of these two species. Broscinae make singleton sperm that are filamentous and are individually embedded in a cap-like or sleeve-like spermatostyle. The spermatostyle of *Z. matthewsii* is short, broad, and sperm-like in form (Fig. 8C) such that when sperm are joined to these spermatostyles, they resemble broad-headed sperm. The spermatostyle of *B. insignis* is sleeve-like and elongate (Fig. 8A-B).

#### Reproductive tract observations

The sperm of both species appear to become easily separated from their spermatostyles, and we were generally unable to find sperm joined to spermatostyles in our female preparations. We collected sperm from spermathecae of several female *Z. matthewsii* and found mostly sperm without spermatostyles.

### Non-Harpalinae Carabidae *incertae sedis* (Figs. 8D-H)

#### Species examined

(Table 1). Tribe Gehringiini: *Gehringia olympica.* Tribe Hiletini: *Eucamaragnathus oxygonus.* Tribe Apotomini: *Apotomus* sp. Tribe Promecognathini: *Promecognathus laevissimus.* Tribe Psydrini: *Psydrus piceus*.

#### Sperm overview

Sperm in these beetles are filamentous and variable in length (Table 2). We were unable to visualize the sperm heads of our *G. olympica* and *Apotomus* sp. sperm preparations. The heads of the remaining beetles are thin, tapered anteriorly and more-or-less indistinct from the rest of the cells (e.g., Figs. 8F-H). They are conspicuous with DAPI staining.

*Gehringia olympica, Eucamaragnathus oxygonus,* and *Apotomus* sp. all make singleton sperm (Table 2). We were unable to study the sperm of male *P. piceus,* and we found no evidence for conjugation in our preparation of a female *P. piceus. Promecognathus laevissimus* makes large conjugates by joining hundreds of sperm to a large corkscrew-shaped spermatostyle (Figs. 8D-E). The anterior end of the spermatostyle is spoon-shaped and without sperm (Fig. 8D). The spermatostyle appears to be composed of two parts: a central opaque rod with attached hyaline flanks (Fig. 8D). The sperm are more heavily distributed laterally on the hyaline flanks of the spermatostyle.

#### Reproductive tract observations

*We* recovered several intact and motile sperm conjugates from the spermatheca of our studied female *P. laevissimus* specimen.

#### Sperm motility observations

*Promecognathus laevissimus* conjugates move in the direction of their anterior end and spin in a helical fashion as they swim perhaps due to the shape of the corkscrew-shaped spermatostyle and the action of its hundreds of attached sperm (Supporting Information MV10-MV11).

### Tribes Clivinini and Dyschiriini (Fig. 9)

#### Species examined

(Table 1). Tribe Clivinini: *Ardistomis obliquata, Ardistomis schaumii, Aspidoglossa subangulata, Paraclivina bipustulata, Clivina fossor, Schizogenius litigiosus,* and *Semiardistomis viridis.* Tribe Dyschiriini: *Akephorus obesus, Dyschirius thoracicus, Dyschirius dejeanii, Dyschirius globosus, Dyschirius haemorrhoidalis, Dyschirius pacificus,* and *Dyschirius tridentatus*.

#### Sperm overview

Sperm in Clivinini and Dyschiriini are diverse (Table 2). Sperm length varies from moderately short to long. Sperm heads are filamentous (tribe Clivinini) or short and generally broad and distinctively shaped (tribe Dyschiriini). We were consistently unable to identify the heads of clivinine sperm, and we did not collect any morphometric data on their sperm heads. Sperm heads in Dyschiriini are typically broad and asymmetrical and possibly species-or lineage-specific in shape (Figs. 9I-N). The putative mitochondrial derivatives of Clivinini but not Dyschiriini are conspicuous with DAPI staining and frequently form complex loops that can be mistaken for sperm heads.

**Figure 9.**
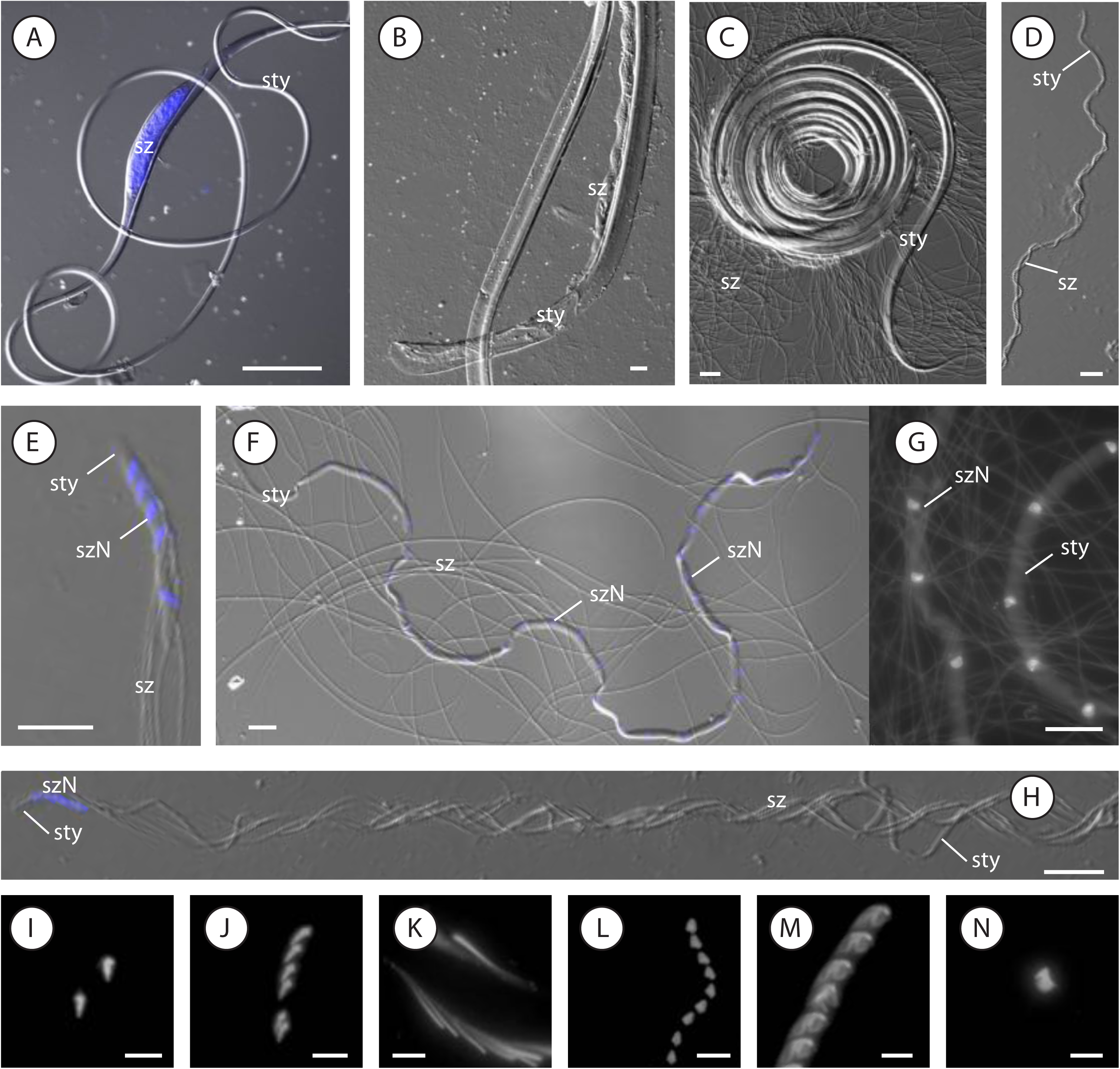
Sperm and sperm conjugate morphological variation in Clivinini (A-D) and Dyschiriini (E-N) (Scaritinae *partim)* ground beetles. (A) sheet conjugate of *Clivina fossor.* The spermatostyle of *C.fossor* contains a central cavity where sperm are housed. (B) large sheet conjugate of *Aspidoglossa subangulata,* note the asymmetrical attachment of sperm. (C) rod conjugate of *Ardistomis obliquata.* (D) sheet conjugate of *Schizogenius litigiosus,* note the wrapping of sperm around the spermatostyle. (E) rod conjugate of *Akephorus marinus,* note the broad and triangular sperm heads. (F-G) rod conjugate of *Dyschirius tridentatus,* note the regular distribution of sperm in the spermatostyle (G). (H) rod conjugate of *Dyschirius dejeanii.* Although the spermatostyle and sperm of *D. tridentatus* and *D. dejeanii* are similar, the arrangement of their sperm is very different. (I-N) *Dyschirius* sperm heads: (I) *D. dejeanii,* (J) *D. pacificus,* (K) *D. globosus,* (L) *D. haemorrhoidalis,* (M) D. *thoracicus,* (N) *D. tridentatus.* (A, E, F, H) stacked image of DIC and Fluoresecence microscopy images. (B-D) DIC microscopy. (G, I-N) Fluorescence images with only DAPI-stained structures visible. sty = spermatostyle, sz = spermatozoa, szN = sperm nuclei. Scale bars: 5 μm (I-N), 20 μm (B-H), 100 μm (A).

All clivinines and dyschiriines studied to date make sperm conjugates (Table 2). The sperm conjugates all include a spermatostyle, but there is notable variation within these groups at the level of the conjugate. The sperm conjugates of Clivinini tend to be either rod conjugates (Fig. 9C) or sheet conjugates [*cf.* Sasakawa 2007; Fig. 9B). The spermatostyle varies substantially in length between Clivinini species as does the number of sperm in a conjugate (Table 2). The rod conjugates of Dyschiriini generally include less than 35 embedded sperm paired to spermatostyles of varying lengths (Figs. 9E-H). The sperm conjugates of Dyschiriini are unusual among carabids because they include so few sperm in a conjugate that the sperm can be easily counted (e.g., Figs. 9E-H). Perhaps because of their typically broad size, the sperm heads of Dyschiriini are arranged in a neat row within the spermatostyle (Fig. 9E) and are never placed parallel to one another.

#### Unusual conjugate-level variation

The sperm conjugates of some Clivinini and Dyschiriini are particularly unusual in that they include more spermatostyle than sperm. For example, the sperm conjugate of A *subangulata* includes a large 6600μm spermatostyle, of which less than l/6^th^ of its length bears sperm; the rest of the spermatostyle is completely bare. Sperm in *A. subangulata* conjugates are distributed on only one side of the spermatostyle further biasing the conjugate towards spermatostyle and less towards sperm. The sperm conjugates of *Clivina* are unusual in that their spermatostyles feature an expanded cavity where the sperm are sealed (Fig. 9 A), and based upon observations during dissections, it appears as though the conjugates are not motile. Upon rupturing the apical portion of the spermatostyle of *Clivina* conjugates, sperm were released via a narrow internal duct subtending the cavity.

#### Within-species variation

Many of the sperm traits we recorded for Clivinini and Dyschiriini sperm show high degrees of variance between preparations. We suspect that these large variances are largely symptomatic of our preparation of the sperm and sperm conjugates of these beetles, which are lengthy and easily damaged.

#### Within-genera variation

*We* studied more than one species of *Dyschirius, Clivina,* and *Ardistomis.* Species of *Clivina* and *Ardistomis* have largely similar sperm conjugates that differ slightly in the size and shape of the spermatostyle as well as in the lengths of their sperm and mitochondrial derivatives. *Dyschirius,* however, appears to be an especially interesting group of Carabidae in which to study the evolution of sperm form, sperm-female morphological coevolution, and in which to explore the possibility of using sperm form for species delimitation. We have studied a handful of different species of *Dyschirius,* and it is clear that sperm, particularly head shape, evolves rapidly within this group. The sperm heads are frequently complex in shape and notably different from one lineage to the next. Understanding the extent to which sperm head shape varies within *Dyschirius* species was not a goal of this study, and we note that these data are still preliminary.

#### Reproductive tract observations

The large sheet conjugates of some clivinines such as *A. subangulata* appear to occupy a large amount of space in the male reproductive tract and are relatively few in number. These large conjugates can be particularly difficult to extract undamaged.

### Subfamily Scaritinae excl. the tribes Clivinini and Dyschiriini (Figs. 10A-D)

#### Species examined

(Table 1). Tribe Scaritini: *Haplotrachelus atropsis, Haplotrachelus cf. latesulcatus, Haplotrachelus* sp., *Scarites marinus, Scarites (Distichus)* sp., and *Scarites (Parallelomorphus)* sp. Tribe Pasimachini: *Pasimachus californicus*.

#### Sperm overview

Sperm in these groups are moderately short in total length (Table 2). The sperm are filamentous, and the heads are visually indistinct. This grouping of beetles make sperm with two regions of fluorescence following DAPI staining similar to Patrobini sperm and the sperm of the vast majority of Harpalinae that we studied (Fig. 10B-D). The sperm heads are inconspicuous and weakly fluorescent compared to the intensely fluorescent mitochondrial derivatives. The heads are small and compact (Figs. 10B-C), and the mitochondrial derivatives are significantly longer than the sperm heads and average between 70-78μm in *P. californiens* and *S. marinus* (Fig. 10D).

**Figure 10.**
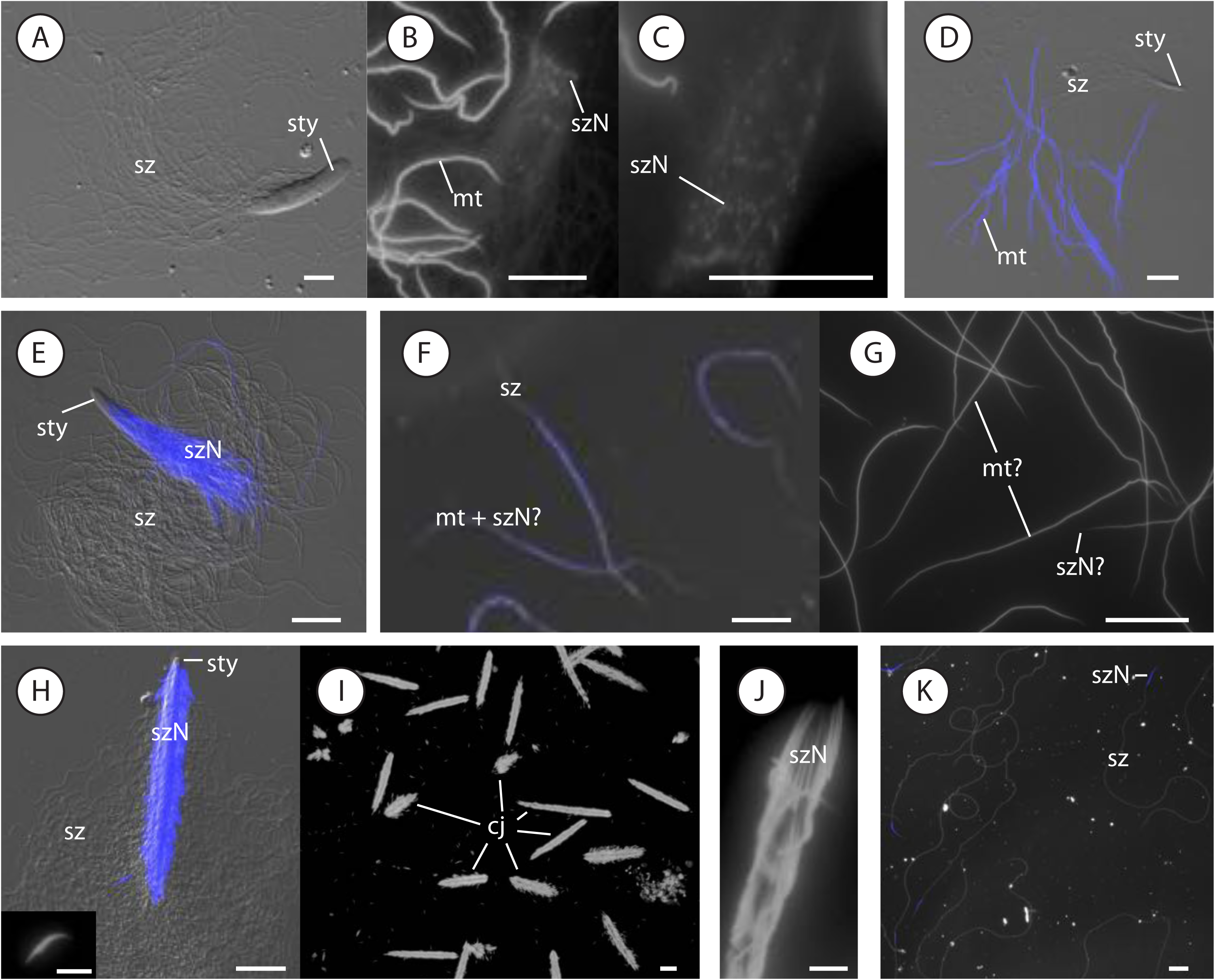
Sperm and sperm conjugate morphological variation in ground beetles of the subfamilies Scaritinae (excluding Clivinini and Dyschiriini) (A-D), Rhysodinae (E), Cicindelinae (F-G), and Paussinae (H-K). (A-C) rod conjugates of *Pasimachus californicus* with small weakly fluorescent sperm heads (B-C) and large intensely fluorescent mitochondrial derivates (B). (D) rod conjugate of *Scarites marinus,* note the conspicuous mitochondrial derivatives. (E) rod conjugates of *Omoglymmius hamatus* include sperm with only one obvious region of fluorescence following DAPI staining. (F-G) singleton sperm of *Brasiella wickhami,* note the small gap in fluorescence between the suspected nucleus and mitochondrial derivatives (G). (H) Composite image of *Metrius contractus* rod conjugate and slightly broad sperm head (inset). (I) conjugate size polymorphism in *Metrius contractus.* (J) closeup of rod conjugate of *Goniotropis parca,* note the linear arrangement of slender-headed sperm. (K) singleton sperm of an unidentified species of *Cerapterus.* (A) DIC microscopy. (D-E, H) stacked image of DIC and Fluoresecence microscopy images. (F, K) stacked image of Darkfield and Fluoresecence microscopy images. (B-C, G, inset of H, I-J) Fluorescence images with only DAPI-stained structures visible, cj = conjugate, mt = mitochondrial derivatives, sty = spermatostyle, sz = spermatozoa, szN = sperm nuclei. Scale bars: 5 μm (inset of H, J), 20 μm (A-G, I, K).

The species we studied in *Haplotrachelus, Pasimachus,* and *Scarites* all make sperm conjugates with small, cap-like spermatostyles or short rod-like spermatostyles (Table 2; Figs. 10A, D). Sperm are embedded in the spermatostyle via their small, compact heads; their flagella are unbounded. The spermatostyle is short and cap-like in *S. marinus* and in an unidentified species of *Scarites* subgenus *Parallelomorphus.* The remaining species that we studied all make rod spermatostyles that are more elongate. *Haplotrachelus* males make spermatostyles that are noticeably less rigid than the spermatostyles of other beetles in this group and their apices are flattened and spatulate.

#### Within-species variation

*We* found notable variation in sperm conjugate size between specimens of *P. californiens* (Supporting Information Spreadsheet SI). *Pasimachus californiens* sperm appear to be monomorphic, but their spermatostyles differ in average length between specimens. The spermatostyles also differ in shape with some spermatostyles appearing short and oblong or stretched posteriorly and elongated. The differences in spermatostyle size and shape influence the average number of sperm in a conjugate, and males show large variances in the average number of embedded sperm in their conjugates.

Sasakawa, (2009) studied the sperm of a Japanese species of *Scarites, S. terricola,* and found an unusual example of within-male variation in sperm. *Scarites terricola* males makes a short filamentous sperm morph that looks similar to the sperm of close relatives and is involved in conjugation, and a second sperm morph that is large and macrocephalic and always present as singletons. These sperm traits are distinct from other cases of sperm dimorphism in adephagan beetles like those seen in many diving beetles (Higginson et al., 2012a) because the two different sperm forms of *S. terricola* do not combine to make a conjugate.

#### Within-genera variation

Large-bodied Scaritinae are frequently known to be morphologically homogenous and taxonomically challenging (e.g., Jeannel, 1941; Nichols, 1988). We studied three different species of the Old World scaritine genus *Haplotrachelus* and three likely distantly related species of the cosmopolitan genus *Scarites. We* observed minor differences in sperm length between these species compared to their congeners. *Haplotrachelus* species all make remarkably similar spermatostyles that differ slightly in size. The conjugates of the *Scarites* species we examined differ in shape, size, and number of included sperm.

#### Sperm ultrastructure

Witz, (1990) studied the sperm form of two species of *Pasimachus, P. strenuus* and *P. subsulcatus.* He found that *Pasimachus* sperm include a small, electron dense nucleus with two adjacent large mitochondrial derivatives with a herringbone pattern of paracrystaline material in cross section (Witz, 1990). Their sperm have a typical 9+9+2 arrangement of microtubules in the axoneme, and he was unable to discern an acrosome in the mature sperm of *P. subsulcatus.* Witz, (1990) also found a series of small microtubules adjacent to the nucleus and developing mitochondria in *Pasimachus* spermatids and hypothesized that these are involved in organelle elongation in the mature sperm.

#### Reproductive tract observations

Males in these genera of Carabidae all have a blind sac termed a vesicula seminalis (Will et al., 2005) that branches off of the vas deferens prior to meeting the accessory glands. We ruptured the vesicula seminalis of our studied male beetles and consistently found it to contain numerous sperm conjugates.

### Subfamily Rhysodinae (Fig. 10E)

#### Species examined

(Table 1). Tribe Clinidiini: *Clinidium* sp. *nr. guatemalenum.* Tribe Omoglymmiini: *Omoglymmius hamatus*.

#### Sperm overview

Rhysodinae sperm are moderately short and filamentous (Table 2; Fig. 10E). The sperm heads are thin and filamentous and visually indistinct from the rest of the cells. The heads are conspicuous with DAPI, and the mitochondrial derivatives of rhysodine sperm are not visible following DAPI staining.

Rhysodines make sperm conjugates with a relatively short and oblong rod-like spermatostyle (Table 2; Fig. 10E). The sperm are embedded in the spermatostyle via their heads with unbounded flagella.

#### Reproductive tract observations

Male rhysodines also have a vesicula seminalis (Will et al., 2005). We ruptured the vesicula seminalis of our studied male beetles and recovered numerous sperm conjugates. The accessory glands of rhysodines are unusual among carabids for their very elongate tips that are compacted inside their bodies (Will et al., 2005).

### Subfamily Cicindelinae (Figs. 10F-G)

#### Species examined

(Table 1). Tribe Amblycheilini: *Omus audouini* and *Omus dejeanii.* Tribe Cicindelini: *Brasiella wickhami* and *Cicindela haemorrhagica.* Tribe Megacephalini: *Tetracha Carolina*.

#### Sperm overview

Sperm in Cicindelinae are short and filamentous with little variation in length across the group (Table 2; Fig. 4A). The heads are filamentous, tapered anteriorly, and visually indistinct from the remainder of the cells. DAPI staining typically reveals one large region of fluorescence nearly two-thirds of the length of the sperm or more, sometimes with a more or less isolated small, lanceolate region of weak fluorescence apically (Fig. 10F-G). Werner, (1965) studied the sperm of a European tiger beetle, *Cicindela campestris,* using TEM and discovered that the nucleus of their sperm runs parallel to the axoneme and the mitochondrial derivatives similar to some Harpalinae (Dallai et al., 2019). Werner, (1965) also found that the nucleus ends before the mitochondrial derivatives and other axonemal structures. We suspect that with DAPI staining and light microscopy, we are visualizing both the mitochondrial derivatives and the nucleus of tiger beetle sperm (Fig. 10F-G). Because tiger beetles may all possess a nucleus that runs parallel to their axoneme similar to *C. campestris* and because we could not easily identify a gap in fluorescence between the mitochondrial derivates and the nucleus, we did not record sperm head measurements of our tiger beetle sperm preparations. The mitochondrial derivatives are filamentous and intensely fluorescent following DAPI staining. They range in length from 71μm in *C. haemorrhagica* to 110μm in *O. dejeanii*.

All Cicindelinae studied to date make only singleton sperm, and we have seen no evidence of spermatostyle production in any cicindeline preparation (Table 2).

#### Within-genera variation

We studied two species of the North American genus *Omus* from the Pacific Northwest, *O. dejeanii* and *O. audouini.* The sperm of the two species differ very slightly in total length and length of their mitochondrial derivates.

#### Reproductive tract observations

Perhaps because of their small sperm and lack of conjugation, we consistently found the seminal vesicles of male Cicindelinae to be filled with very large quantities of individual sperm.

### Subfamily Paussinae (Figs. 10H-K)

#### Species examined

(Table 1). Tribe Metriini: *Metrius contractus.* Tribe Ozaenini: *Goniotropis parca, Ozaena* sp., and *Pachyteles* sp. Tribe Paussini: *Cerapterus* sp., *Paussus cucullatus,* and an unidentified species of *Paussus (Bathypaussus)*.

#### Sperm overview

Sperm in Paussinae (Table 2) are filamentous and short when conjugated (Fig. 10H-J) or filamentous and moderately long when singletons (Fig. 10K). The sperm of *Goniotropis parca* is currently the shortest known sperm for the family Carabidae (Table 2). The heads of paussine sperm are thread-like and tapered anteriorly or slightly thickened and relatively short as in *Metrius contractus* (Figs. 10H-K). The heads are conspicuous with DAPI staining as a single region of fluorescence. The mitochondrial derivatives are not obvious following DAPI staining.

Paussines either make sperm conjugates or singleton sperm (Table 2). *Metrius contractus* and *Goniotropis parca* package their sperm into sperm conjugates with a moderately short rod-like spermatostyle (Figs. 10H-J). Species of the tribe Paussini, which include many obligate ant nest parasites, were found to make only singleton sperm (Fig. 10K). We only sampled a single female of *Pachyteles* and *Ozaena,* and we were unable to determine if they make conjugates. The sperm of *M. contractus* and *G. parca* are embedded in the spermatostyle via their heads with their flagella unbounded. Sperm appear to be distributed on all sides of the spermatostyle in the conjugates of *M. contractus* (Fig. 10H), but in *Goniotropis parca,* the sperm are generally located laterally on the spermatostyle with a prominent bare region medially (Fig. 10J).

#### Within-species variation

The sperm conjugates of *Metrius contractus* show high levels of polymorphism in size between specimens (Fig. 10I), but their sperm appear to be monomorphic. Because sperm are distributed throughout the vast majority of the length of the spermatostyle, these conjugates also vary in the number of included sperm. *Goniotropis parca* males, similarly, have variation between specimens in conjugate size with monomorphic sperm, but this variation is smaller than what we have observed in *M. contractus*.

#### Within-genera variation

We studied two species of the obligate ant-parasite genus *Paussus, P. cucullatus* (subgenus *Hylotorus)* and an unidentified species of the subgenus *Bathypaussus.* The sperm of these two species differ slightly in total length and head length. We also observed rather different sperm lengths but not head lengths between our specimens of *P. cucullatus* from two different populations (Supporting Information Table S2).

#### Reproductive tract observations

Male paussines all have a blind sac termed a vesicula seminalis (Will et al., 2005) that joins their vas deferens prior to its meeting with the accessory glands. We ruptured the vesicula seminalis of our studied male beetles and recovered numerous sperm conjugates.

### Subfamily Brachininae (Figs. 11A-D)

#### Species examined

Tribe Brachinini: *Brachinus elongatulus, Brachinus ichabodopsis, Mastax* sp., *Pheropsophus* sp. 1, *Pheropsophus* sp. 2.

#### Sperm overview

Brachininae sperm are filamentous and short with little variation in sperm total length (Table 2). The sperm heads are generally short, tapered anteriorly, and visually indistinct from the rest of the cells. Following DAPI staining, Brachininae sperm show two regions of fluorescence: the large and intensely fluorescent mitochondrial derivatives and the notably fainter, small and compact sperm heads (Fig. 11B). The sperm heads are short and narrow, under 1μm in length and width. The mitochondrial derivatives vary slightly in length between species, but they are generally conspicuous following DAPI staining and measure two-thirds of the total length of the sperm or more (Fig. 11B).

**Figure 11.**
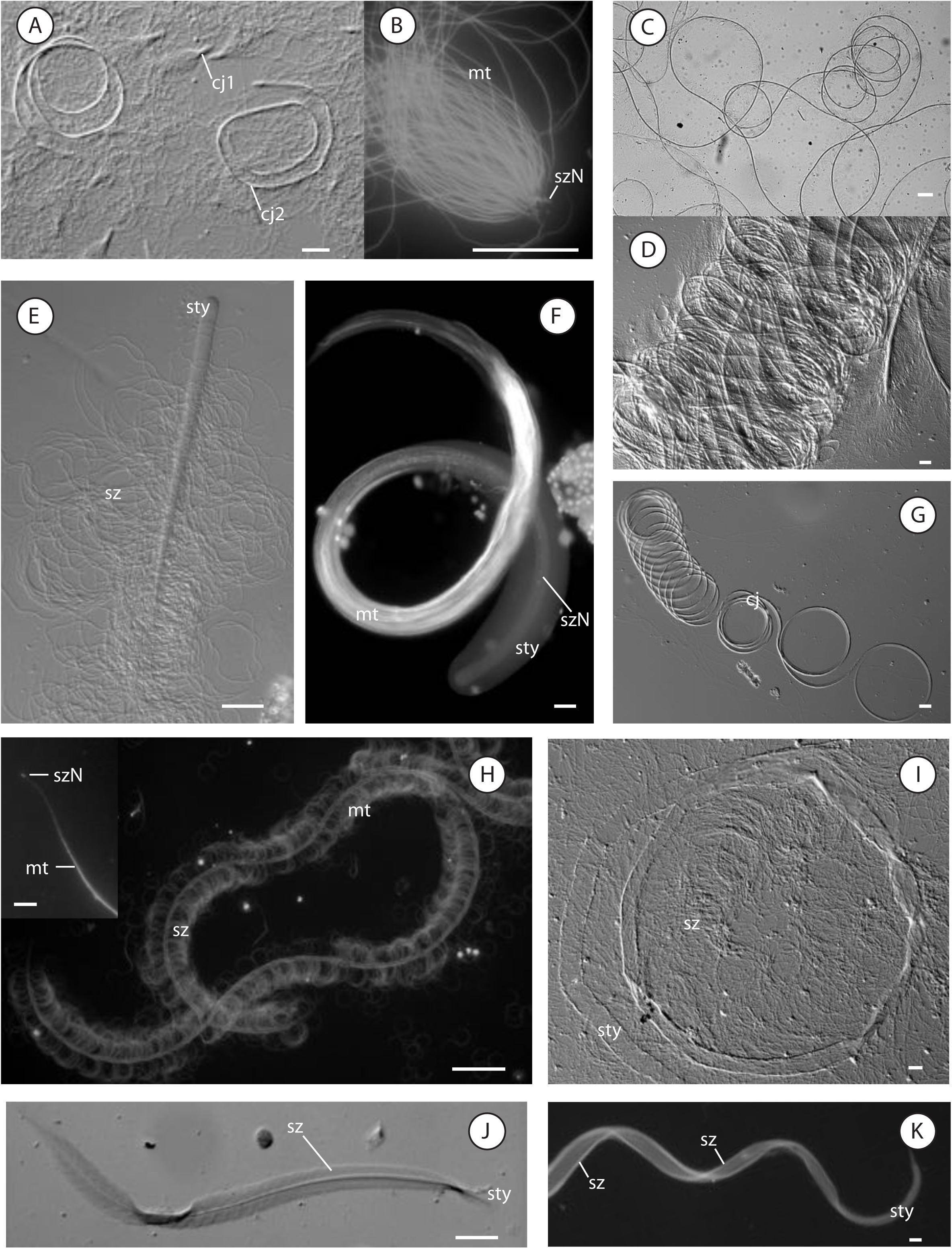
Sperm and sperm conjugate morphological variation in Brachininae (A-D) and Harpalinae (E-K) ground beetles. (A) *Brachinus elongatulus* males have monomorphic sperm but package them into two distinct rod conjugates. (B) closeup of small rod conjugate of *Brachinus elongatulus* showing the small weakly fluorescent sperm nuclei and the large intensely fluoresecent mitochondrial derivatives. (C-D) giant sperm conjugates of *Pheropsophus,* which reach up to 5.8 cm. (E) rod conjugate of *Agonum piceolum,* (F) sheet conjugate in an unidentified species of *Leptotrachelus.* (G) slinky-like sheet conjugate of *Chlaenius ruficauda.* (H) rod conjugate in an unidentified speices of *Bradycellus.* (I) rod conjugate of *Stenocrepis elegans,* note the thin, ribbon-like spermatostyle. (J) feather-like sheet conjugate of *Calleida jansoni.* (K) sheet conjugate of *Tetragonoderus fasciatus,* note the wavy spermatostyle and the bilateral attachment of sperm. (A, D, E, G, I-J) DIC microscopy. (B, F, H, K) Fluorescence images with only DAPI-stained structures visible. (C) Brightfield microscopy, cj = conjugate, mt = mitochondrial derivatives, sty = spermatostyle, sz = spermatozoa, szN = sperm nuclei. Scale bars: 5 μm (inset of H), 20 μm (A-B, D-G, I-K), 100 μm (C, H).

Brachininae all make sperm conjugates with either a short, cap-like spermatostyle and/or a slender and elongate rod-like spermatostyle (Figs. 11A-D). The sperm are embedded in the spermatostyle via their heads with unbounded flagella. *Brachinus elongatulus* is unusual because males make two distinct sperm conjugate morphs (Table 2; Fig. 11A). One of the sperm conjugate morphs of *B. elongatulus* includes between 30-70 sperm joined to a small, cap-like spermatostyle. The second conjugate morph of *B. elongatulus* is composed of hundreds of sperm, which we were unable to estimate accurately, joined to an elongate, ribbon-like spermatostyle. *Mastax* males make small conjugates with a cap-like spermatostyle that resemble the small conjugate morph of *B. elongatulus.* The remaining brachinines we studied make larger spermatostyles including the giant 41mm-long spermatostyles of the large-bodied bombardier genus *Pheropsophus. Pheropsophus* sperm conjugates currently hold the record for largest sperm conjugates in Carabidae and are likely among the largest sperm conjugates known. The giant spermatostyles of *Pheropsophus* are flexible and ribbon-like, forming numerous loops of varying sizes on the slide (Figs. 11C-D). The sperm of *Pheropsophus* conjugates are regularly distributed throughout the length of the spermatostyle. Although we cannot accurately estimate the number of sperm in these giant conjugates, they likely include thousands of sperm given the dense packing of sperm and the giant size of the spermatostyle.

#### Within-species variation

Our measurements of *Pheropsophus* sperm conjugates include a high amount of variance, between 4-5mm, in spermatostyle length between specimens. This variation in conjugate size may be accurate, but it may be an artifact of our preparations caused by the large size of these otherwise thin structures.

#### Within-genera variation

We studied two different, likely distantly related species of the large complex genus *Brachinus* and found obvious differences in sperm form. *Brachinus elongatulus* sperm differ from sperm in *B. ichabodopsis* in total length and the size of their mitochondrial derivatives. *Brachinus elongatulus* make two distinct sperm conjugates whereas *B. ichabodopsis* makes a single conjugate morph with a very slender elongate spermatostyle that does not resemble the spermatostyle of either conjugate morph in *B. elongatulus*.

### Subfamily Harpalinae (Figs. 11E-K; 12B-F)

#### Species examined

Tribe Abacetini: *Abacetus* sp., *Stolonis intercepta,* and *Stolonis* sp. Tribe Anthiini: *Anthia (Termophilum)* sp. Tribe Catapiesisini: *Catapiesis* sp. Tribe Chlaeniini: *Chlaenius cumatilis, Chlaenius glaucus, Chlaenius harpalinus, Chlaenius leucoscelis, Chlaenius prasinus, Chlaenius ruficauda, Chlaenius sericeus,* and *Chlaenius tricolor.* Tribe Ctenodactylini: *Leptotrachelus sp.* Tribe Cyclosomini: *Tetragonoderus fasciatus* and *Tetragonoderus* sp. nr *latipennis.* Tribe Dryptini: *Drypta* sp. Tribe Galeritini: *Galerita atripes, Galerita bicolor, Galerita forreri,* and *Galerita lecontei.* Tribe Graphipterini: *Cycloloba* sp. and *Graphipterus* sp. Tribe Harpalini: *Anisodactylus alternons, Anisodactylus anthracinus, Anisodactylus similis, Bradycellus* sp. 1, *Bradycellus* sp. 2, *Discoderus* sp., *Euryderus grossus, Harpalus affinis, Polpochila erro, Selenophorus* sp., *Stenolophus* sp., and *Stenomorphus convexior.* Tribe Helluonini: *Helluomorphoides papago* and *Macrocheilus* sp. Tribe Lachnophorini: *Eg a sallei, Lachnophorus* sp. nr. *elegantulus,* and *Lachnophorus elegantulus.* Tribe Lebiini: *Agra* sp. 1, *Agra* sp. 2, *Apenes lucidula, Calleida bella, Calleida jansoni, Calleida decora, Cymindis punctifera, Cymindis punctigera,* an unidentified member of the *basipunctata-* group of *Cymindis* subgenus *Pinacodera, Lebia deceptrix, Lebia subgrandis, Lebia viridis, Phloeoxena nigricollis, Stenognathus quadricollis, Syntomus americanus,* and *Thyreopterus flavosignatus.* Tribe Licinini: *Badister ferrugineus, Dicaelus suffusus,* and *Diplocheila nupera.* Tribe Morionini: *Morion* sp. Tribe Odacanthini: *Colliuris pensylvanica.* Tribe Oodini: *Anatrichis minuta, Oodes fluvialis,* and *Stenocrepis elegans.* Tribe Panagaeini: *Panagaeus sallei.* Tribe Peleciini: *Disphaericus* sp. Tribe Perigonini: *Perigona nigriceps.* Tribe Pentagonicini: *Pentagonica sp.* Tribe Platynini: *Agonum piceolum, Agonum muelleri,* an unidentified species of *Rhadine dissecta-group,* and *Sericoda bembidioides.* Tribe Pseudomorphini: *Pseudomorpha* sp. Tribe Pterostichini: *Abaris splendidula, Hybothecus flohri, Cyclotrachelus dejeanellus, Cyrtomoscelis* cf. *dwesana, Pterostichus (Morphnosoma) melanarius, Pterostichus (Hypherpes) lama, Pterostichus (Leptoferonia) infernalis, Poecilus laetulus,* and *Poecilus scitulus.* Tribe Sphodrini: *Calathus peropacus* and *Synuchus dubius.* Tribe Zabrini: *Amara aenea* and *Amara farcta.* Tribe Zuphiini: *Pseudaptinus horni, Pseudaptinus simplex,* and *Pseudaptinus tenuicollis*.

#### Sperm overview

The large subfamily Harpalinae, containing half of all carabid species, have sperm that are filamentous and vary widely in length from short to long (Table 2). Instances of both short and long sperm occur repeatedly throughout the subfamily. The sperm heads are typically inconspicuous and visually indistinct from the remainder of the cells. Following DAPI staining, Harpalinae sperm show one or two regions of fluorescence. The mitochondrial derivates are large and fluoresce intensely with DAPI staining (Fig. 11H) whereas the nuclei are weakly fluorescent, making morphological observation of the heads difficult. There are several specimens for which we were unable to clearly discern the head (41.5% of all Harpalinae preparations studied). Of the sperm heads that we could observe, our data show that Harpalinae sperm heads are short, commonly between 0.5-5.0μm in length (Table 2; Fig. 11H). The heads are generally tapered anteriorly or weakly asymmetrical and narrow, varying minimally in width.

Sperm conjugation seems to be the rule across Harpalinae with only few ambiguous exceptions (Table 2; Figs 3,11E-K). Sperm conjugation involves a spermatostyle, and the variation in conjugate shape and size across Harpalinae is striking, with numerous species making particularly elongate spermatostyles. The spermatostyle varies in total length from the short and cap-like spermatostyle of *Chlaenius prasinus* (Chlaeniini) to the enormous rod-like spermatostyles of *Pterostichus lama* (Pterostichini) and *Diplocheila nupera* (Licinini) that are among the largest spermatostyles in Carabidae (Figs. 3-4). The spermatostyle varies dramatically in shape with some species making corkscrew-shaped or spiral spermatostyles (e.g., *Anisodactylus alternons, Tetragonoderus fasciatus;* Fig. 11K), flat and ribbon-like spermatostyles (e.g., *Stenocrepis elegans;* Fig. 11I), slinky-shaped spermatostyles (e.g., *Chlaenius ruficauda;* Fig. 11G), or small and cap-like spermatostyles, in addition to variations on the more common simple rod-like spermatostyle (Fig. 11E). The apex of the spermatostyle shows a lot of variation and is frequently distinct in shape and/or width from the remainder of the spermatostyle. The apex is frequently simply tapered or gently expanded, but we have seen species with spermatostyles that are spoon­shaped or spatulate apically (e.g., *Pterostichus nigrita* Hodgson et al., 2013, *A. alternons, Cymindis punctigera)* or jagged and knife-like [*Poecilus* species).

Harpalinae make either rod or sheet conjugates (Fig. 4). In sheet conjugates, the sperm, including their flagella, are joined to the spermatostyle by hyaline material (Figs. 11F, J-K; Dallai et al., 2019; Sasakawa, 2007). We have observed sheet conjugates in various unrelated groups of ground beetles. Sheet conjugates occur in members of the following tribes: Abacetini (e.g., *Abacetus* sp.), Chlaeniini (e.g., some but not all *Chlaenius* we studied), Ctenodactylini (e.g., *Leptotrachelus* sp.), Cyclosomini (e.g., *Tetragonoderus* spp.), Lebiini (e.g., *Calleida jansoni, Lebia* spp., and *Syntomus americanus),* Lachnophorini (e.g., a Mexican species of *Lachnophorus),* and Harpalini (e.g., *Discoderus* sp., *Stenolophus* sp., and *Stenomorphus convexior).* Most of the Pterostichini we studied also make sheet conjugates. Males of the genus *Galerita* make sperm conjugates that we scored as sheet conjugates. *Galerita* conjugates are unusual in that the spermatostyles include a long groove that seems to be associated with sperm placement (Fig. 12D), reminiscent of the sperm conjugates of some *Clivina.* Typically sperm are distributed more or less evenly along the entire length of the spermatostyle in Harpalinae. Sometimes sperm are more densely distributed along the sides of the spermatostyle (common in sheet conjugates (Hodgson et al., 2013)) or along particular stretches of the spermatostyle resulting in prominent bare regions that are common posteriorly (e.g., the spermatostyle of *Euryderus grossus* is 2900μm long but only 200-300μm of its length bears sperm). The number of sperm in a conjugate varies dramatically within Harpalinae. Most of the Harpalinae we studied make conjugates with between 30-1000 sperm in a conjugate. We were not always able to estimate the number of sperm in Harpalinae conjugates particularly when the conjugates were very large (e.g., the 9mm conjugates of *P. lama)*.

**Figure 12.**
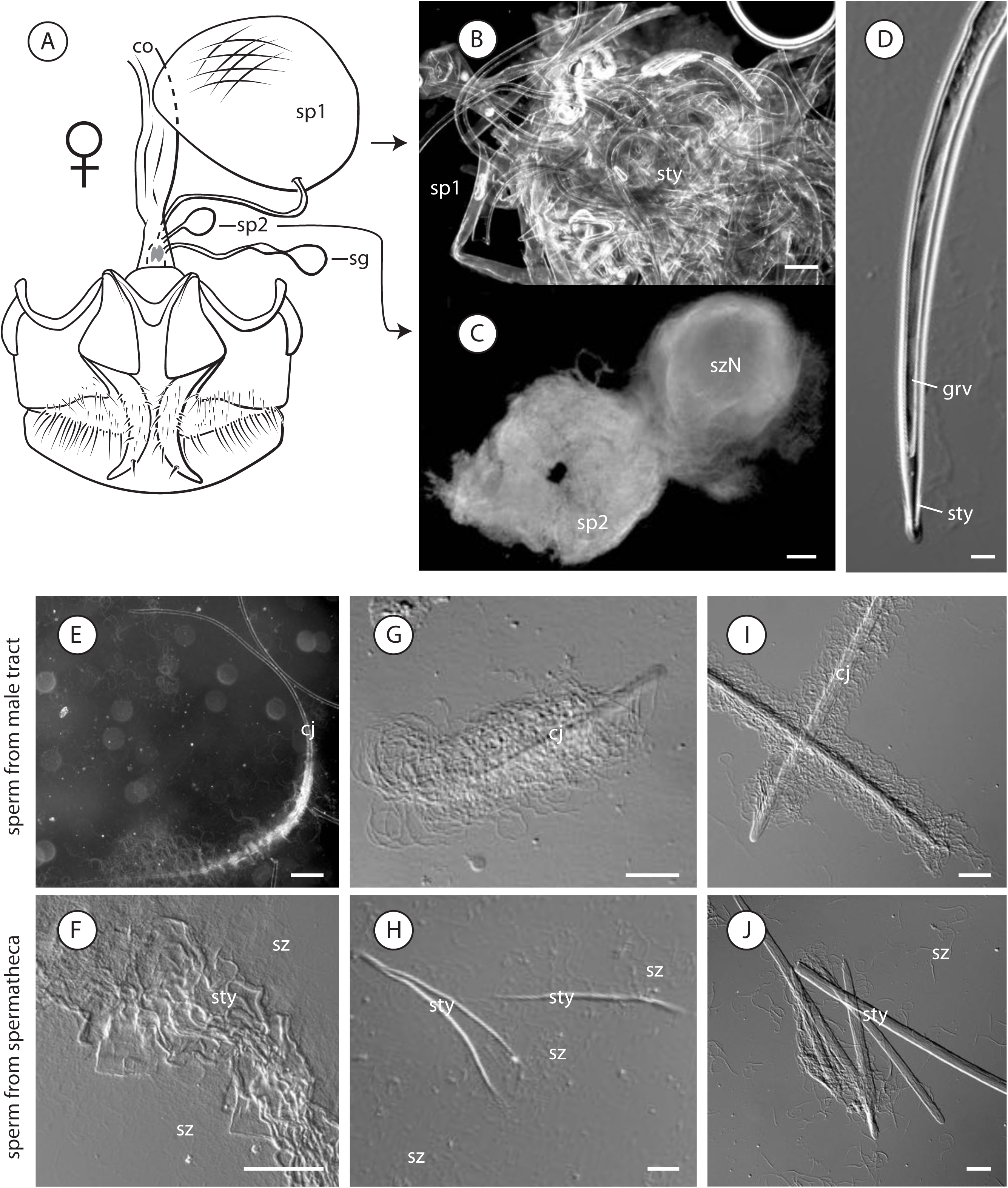
Sperm + female reproductive tract interactions observed in ground beetles. (A-C) *Galerita* sperm + female interactions. (A) *Galerita bicolor* female reproductive tracts include two sperm storage organs that store different parts of a male’s conjugate, redrawn from Liebherr and Will (1998). (B) A large mass of bare spermatostyles recovered from the large balloon-like spermatheca of *Galerita atripes.* (C) A large bolus of sperm recovered from the smaller spherical sperm storage organ (= secondary spermathecal gland of Liebherr and Will (1998)) of *Galerita bicolor.* (D) closeup of the spermatostyle of *Galerita forreri,* note the presence of a groove where we suspect sperm are attached. (E-J) sperm conjugates before and after storage in female reproductive tracts, note the dissociation of sperm from spermatostyles and morphological changes to spermatostyles. (E, G, I) sperm from male preparations. (F, H, J) sperm from female spermathecae. (E-F) *Harpalus affinis.* (G-H) *Elaphrus purpurans.* (I—J) *Sphaeroderus stenostomus.* bc = bursa copulatrix, cj = conjugate, co = common oviduct, grv = groove, sg = spermathecal gland, sp1 = spermatheca 1, sp2 = spermatheca2, sty = spermatostyle, sz = spermatozoa, szN = sperm nuclei. Scale bars: 20 μm (D, G-J), 100 μm (B-C, E-F).

We found no unambiguous evidence that conjugation is missing in any of the Harpalinae we studied. However, we were unable to discern some morphological details of conjugation in some of our lower quality preparations. In our *Anthia* and *Galerita* preparations, for example, it was difficult to identify whether sperm had simply become detached from the spermatostyle or were not physically associated with it in the first place.

#### Within-species variation

Male Harpalinae frequently make large conjugates with large spermatostyles. The spermatostyles are typically wider anteriorly than posteriorly and frequently possess a long and thin tail that is easily broken. In several of our harpaline preparations, we recorded large spermatostyle length variances between specimens of a given species. We suspect that some of the variation that we have observed is due to our damaging these spermatostyles during slide preparation.

#### Within-genera variation

We studied two or more species in several widely related Harpalinae genera. Harpalinae sperm conjugates tend to be more morphologically variable than sperm between species. Sperm frequently will differ slightly in total length between species. For example, *Agonum piceolum* and *Agonum muelleri* make sperm that differ slightly in length by nearly 20μm, but their conjugates are notably different. The sperm conjugates of A *muelleri* include a longer spermatostyle that is straight and rigid and includes a small bare region apically. We found a similar pattern in sperm and sperm conjugate variation among species of *Anisodactylus, Bradycellus, Calleida, Galerita, Cymindis,* and *Lebia*.

Conjugate type is typically stable within a genus, but some harpaline genera include species that make either rod or sheet conjugates (e.g., *Chlaenius,* present study; *Pterostichus,* Sasakawa, 2007). We had the opportunity to study several different North American species of the large cosmopolitan ground beetle genus *Chlaenius. We* studied eight different species of *Chlaenius* classified in 5 different subgeneric groupings. Most studied species of *Chlaenius* make rod conjugates, but *Chlaenius ruficauda* makes sheet conjugates. The spermatostyle is notably variable in size and shape between species, and the range of variation observed in spermatostyle length across *Chlaenius* is almost as extensive as the variation observed across Carabidae as a whole (Table 2). The sperm of *Chlaenius* varies significantly in total length and, to a lesser degree, in sperm head length. The monophyly of *Chlaenius* subgeneric groups remains an open question, but it appears that sperm, particularly sperm conjugates, evolve rapidly in *Chlaenius*.

#### Sperm ultrastructure

Recently, Dallai et al., (2019) studied the sperm ultrastructure of several *Pterostichus* species, *Amara aulica,* and *Demetrias atricapillus.* The sperm nuclei of these species are long, thin, and parallel to their axonemes like the nucleus of *Cicindela campestris* sperm (Dallai et al., 2019). If most Harpalinae sperm possess long and thin nuclei like these species, then perhaps it is the shape and size of their nuclei that explains the difficulty we had observing the sperm heads in many of our Harpalinae preparations (Fig. 11H). Their sperm have a typical 9+9+2 axoneme flanked by mitochondrial derivatives and small accessory bodies; their heads bear small, flat acrosomes (Dallai et al., 2019). Sperm in these species are packaged into sheet conjugates and are embedded laterally into the sidewall of the spermatostyle via their heads; their flagella are located in chambers that are joined to the spermatostyle by laminar extensions (Dallai et al., 2019).

#### Reproductive tract observations

We collected sperm from female Harpalinae for several of our slide preparations. We almost always found individual sperm and collections of spermatostyles or intact conjugates in the spermathecae of females from throughout our sampling. The spermatostyles of Harpalinae males are generally long, and they tend to be compacted within the female’s sperm storage organ. For example, male *Harpalus affinis* sperm conjugates include a long, slender, sickle-shaped spermatostyle (Fig. 12E), and we recovered large haphazard spermatostyle masses from the spermathecae of female *H. affinis* (Fig. 12F). Compacted masses of spermatostyles like these were commonly observed near the entrance of the spermatheca or the spermathecal duct with individual sperm predominately occupying the apical regions of the spermatheca. These masses were frequently difficult to break apart and mirrored the shape of the spermatheca.

Some female harpaline ground beetles use different storage organs for different parts of a male’s sperm conjugate. *Galerita* females appear to use one storage organ for sperm and another storage organ for the spermatostyles that males make (Figs. 12A-C). We dissected females of two different *Galerita* species, and discovered that the large balloon-shaped structure that has been called the spermatheca by Liebherr and Will, (1998) held large numbers of bare spermatosyles (Fig. 12A) and the notably smaller spherical structure termed a secondary spermathecal gland by Liebherr and Will, (1998) contained only individual sperm (Fig. 12C). These two storage sites are physically separated from one another and are connected via separate ducts to a larger common duct that joins the bursa copulatrix (Fig. 12 A; Liebherr and Will, 1998; Hunting, 2008). Nothing is known regarding sperm use by *Galerita* females, but we speculate that conjugates arrive to the balloon-shaped structure, dissociate or become dissociated from their sperm, and sperm travel or are moved to the functional spermatheca. Near relatives of *Galerita* possess similar female reproductive tract forms (Hunting, 2008), and it seems likely that this pattern of decoupled sperm and spermatostyle storage applies more broadly.

#### Sperm motility observations

*Agonum piceolum* sperm conjugates swim in a typical helical fashion, and they appear to swim faster than individual sperm (Supporting Information MV1-MV2).

## 5. Discussion

### Trends in carabid sperm evolution

One of the obvious ways in which sperm vary is in their length, a trait which may covary with conjugation. Long sperm have historically received much attention, and long sperm can be ornaments evolving under sexual selection (Lüpold et al., 2016). Various studies have shown that longer sperm are more costly to produce than shorter sperm (e.g., Pitnick, 1996). Considering only this factor, we would expect that through time sperm would evolve to become shorter (Parker, 1970; 1998). Sperm, instead, show a wide range in lengths in response to a variety of post-mating selection pressures (Lüpold and Pitnick, 2018). We found that carabid sperm vary in length from 48-3400μm, with variation in either direction towards long or short sperm occurring in several large groups of ground beetles, suggesting that sperm size is an evolutionarily labile trait (Figs. 2-4). Many species with singleton sperm tend to make long sperm of over 1mm in length, and perhaps sperm length is correlated with conjugation state or loss of the spermatostyle (Figs. 2-3; Higginson et al., 2012a). Because the spermatostyle competes for space with sperm in male and female reproductive tracts, it may be the case that loss of conjugation with a spermatostyle allows for more space for sperm, which might allow for either longer sperm or more numerous sperm. In diving beetles, sperm length does not correlate with conjugation (Higginson et al., 2012a), and conjugation may have more to do with occupying a site favorable for fertilization rather than conferring any motility advantages to sperm (Higginson et al., 2012b).

Drag and efficient sperm packaging for conjugation may explain why ground beetle sperm are rarely broad-headed. The vast majority of carabid beetles studied make filamentous sperm with visually indistinct heads that are usually no broader than the remainder of the cell (e.g., Fig. 8F). Sperm head length varies from 0.5-270μm (Fig. 4B), but most species have sperm heads that measure under 20μm (Table 2), suggesting that head length evolves much slower than sperm total length. Because sperm live in a low Reynolds number environment where viscous forces dominate over inertial forces (Vogel, 1994), drag is likely an important physical variable in sperm evolution (e.g., Ishimoto and Gaffney, 2015). If drag is an important variable in sperm evolution, you would expect sperm to be broad-headed only rarely (Humphries and Evans, 2008). At the same time, the physical joining of sperm to each other or to a spermatostyle might covary with head shape through time (Higginson and Pitnick, 2011). Higginson et al., (2012a) found that the gain or loss of broad-headed sperm was evolutionarily correlated with qualitative changes to sperm conjugate type. Unlike ground beetles, many diving beetles have broad-headed sperm, but diving beetles do not make sperm conjugates with a spermatostyle (Higginson et al., 2012a). Thick-or broad-headed sperm (1.0-6.3μm) are found in only a few ground beetles such as some Dyschiriiini (Figs. 9I-J, L-N), members of the genus *Omophron* (Figs. 6G, I), genus *Trachypachus* (Fig. 5G), *Eucamaragnathus oxygonus* (Fig. 8H), and *Metrius contractus* (Fig. 10H). Based on the taxonomic distribution of broad-headed sperm (Fig. 4C), it seems likely that broad-headed sperm evolved from slender-headed sperm only a few times in Carabidae. Sperm head width may be more constrained functionally or develoμmentally than other aspects of sperm form. The scope of sperm head width variation is limited, and head width generally varies little between closely related species with narrow-headed sperm. Perhaps broad sperm heads are difficult to produce or are evolutionarily unstable. Diversification models of diving beetle sperm suggest that being broad-headed and single is an evolutionarily unstable state for sperm (Higginson et al., 2012a). All ground beetle species with thickened or broad heads make rod conjugates except for the singleton sperm of *Eucamaragnathus oxygonus.* Sperm conjugate type may not covary with sperm head width in ground beetles, but we surmise that the number of sperm in a conjugate depends on the form of the sperm head. For instance, the asymmetrical and broad-headed sperm of *Omophron* must stack and pair up in such a way to allow for the side bearing the flagellum to be lateral (Figs. 6G, I). Smaller heads might offer less drag, and it may be possible to group together more sperm by their heads when they are small. Species that make small-headed sperm (sperm head 1 x w under 0,5μmχ 0.5μm) typically make conjugates with large numbers of sperm. Whether sperm head width is correlated with the number of sperm in a conjugate remains to be evaluated more thoroughly.

### Trends in carabid sperm conjugate evolution

We identified several different qualitative types of sperm conjugates in ground beetles (Figs. 2-3; Table 2). Within each conjugate type, we found variation in sperm form, spermatostyle form, sperm number, and arrangement of sperm in a conjugate. The different conjugate types therefore only capture a small amount of the continuum of variation found in ground beetle sperm conjugates.

Sperm conjugation with a spermatostyle appears to have been present early in the history of Carabidae. Several studied species make only singleton sperm, but most carabid beetles make sperm conjugates, and the distribution of sperm conjugates suggests an early origin. Most early diverging lineages of ground beetles such as Trachypachinae, Elaphrinae, and Carabinae (Maddison et al., 2009, Maddison et al., unpublished data) make rod conjugates with a spermatostyle and unbounded flagella (Figs. 2, 5A-H). The one exception is subfamily Nebriinae, which make singleton sperm (Figs. 6B-D). The exact position of Nebriinae is an outstanding question in carabid systematics (Arndt et al., 2005; Maddison et al., 1999; 2009), and it may be that the ancestor of all carabids made singleton sperm. However, wherever Nebriinae might be placed, the extent of conjugates throughout carabids outside of Harpalinae suggests that sperm conjugation was present early in the history of Carabidae, and is ancestral for a majority of the family.

Sperm conjugation in ground beetles almost always involves a spermatostyle, and carabids with conjugated sperm tend to make either rod conjugates or sheet conjugates (Figs. 2-3; Sasakawa, 2007). Sheet conjugates occur in several putatively unrelated tribes of Harpalinae (e.g., Figs. 11F-G, J-K) and the tribe Clivinini (Figs. 9A-B). If rod sperm conjugates are ancestral, this would mean that there have been many independent transitions to sheet conjugates.

The distribution of singleton and conjugated sperm across ground beetles suggests that conjugation has been lost at least three times independently. Our low-resolution phylogeny implies a loss of conjugation in Cicindelinae, Paussini, and at the base of Trechitae (Fig. 2). Additional occurrences of singleton sperm are known from other phylogenetically scattered lineages such as Apotomini, Hiletini, and Gehringiini (Fig. 2; Maddison et al., 1999; 2009), but the lack of phylogenetic resolution for these groups prohibits deeper insights into the gain or loss of conjugation in ground beetles.

The subfamily Trechinae is one clade in which patterns of sperm evolution appear evident, in part as there is a well-supported phylogenetic tree on which to examine our results (Maddison et al., 2019). Trechinae males vary in sperm conjugation presence and type (Figs. 2, 7). Some trechines make sperm conjugates with small spermatostyles and short sperm (Figs. 7A-B), some make conjugates without a spermatostyle (Figs. 7D-E), many make singleton sperm (Fig. 7C), and some make long sperm that grapple onto one another, forming haphazard groupings of sperm (Figs. 7G-J; mechanical conjugation). Ancestral Trechinae appear to have had sperm conjugates with short sperm and simple rod-like spermatostyles as seen in the Patrobini (Figs. 7A-B). We hypothesize that the spermatostyle and conjugation were lost in several Trechitae before sperm got longer. Once singleton sperm became longer, some Tachynini sperm gained a different type of conjugation that does not include a spermatostyle or cementing material: their long sperm form large loops (Figs. 7I-J) that turn while they swim, grabbing adjacent sperm in the process. Because mechanical conjugation is thus far known only from muroid rodents and the data we have are limited, further research is needed to confirm that this represents another example of this phenomenon. We note that this sperm evolution model assumes that the loss of sperm conjugation with a spermatostyle is more probable than its being gained and that the mechanical conjugates of some tachyine carabids are not homologous (as conjugates) with the rod conjugates of Patrobini or Bembidiini.

### The spermatostyle as an understudied example of biological novelty

Conjugation with a spermatostyle is an interesting phenomenon because it entails a trade-off between sperm and spermatostyles. The more resources in terms of space, energy, and nutrients a male dedicates to spermatostyle production, the fewer resources are available for sperm production. Reducing sperm production seemingly reduces direct opportunities for paternal DNA to be passed to the next generation. Thus, increasing spermatostyles could reduce potential fertilizations, unless spermatostyles increase the per-sperm probability of successful fertlization. This trade-off in resource utilization also extends to sperm storage in the female’s reproductive tract. In spite of this, most carabids make spermatostyles, and the spermatostyle is frequently large and elaborate among many different species. We know little about the chemical composition of the spermatostyle, but histological evidence suggests that it is a matrix of proteins and carbohydrates (e.g., Hodgson et al., 2013; Schubert et al., 2017). Males sometimes make sperm conjugates with bare regions that lack sperm (e.g., Figs. 6F, 9A, 12E) suggesting that the spermatostyle is more than just a device for joining together sperm.

Evolution has explored a vast amount of morphological space in spermatostyles in ground beetles, and our data suggest that spermatostyles evolve at a much faster rate than sperm. Spermatostyles vary along several axes including size, shape, texture, and thickness. These spermatostyle traits frequently vary between species in a given higher-level taxonomic group (Fig. 2), and we suspect that there is much divergence and convergence in spermatostyle phenotype across Carabidae. Spermatostyle production can also occur without sperm conjugation (Fig. 3), as in Broscinae. Broscinae sperm are singleton, but their sperm are joined to individual spermatostyles (Figs. 8A-C). If spermatostyles evolve faster than sperm, then closely related species should vary more in spermatostyle and conjugate form than sperm form. Our observations of multiple species within 20 genera *(Agonum, Agra, Amara, Anisodactylus, Ardistomis, Brachinus, Bradycellus, Calleida, Carabus, Chlaenius, Cymindis, Galerita, Haplotrachelus, Lachnophorus, Lebia, Pterostichus, Scarites, Sphaeroderus, Stolonis, Tetragonoderus)* indicate that closely related species are more likely to differ in spermatostyle form than sperm form similar to the findings of Takami and Sota, (2007).

We hypothesize that the spermatostyle can be an ornament evolving under postmating sexual selection and that because it is non-cellular, it has been freed from constraints that may be operating on sperm. Spermatostyles that contain large bare regions are particularly interesting from a post-mating sexual selection perspective. If sperm are like lottery tickets (Parker, 1970), this is akin to going to the racing downs and spending lots on beer and little on tickets. It may be the case that large conjugates with few sperm and large spermatostyles are similar to the exaggerated ornaments of some *Drosophila* sperm (Lüpold and Pitnick, 2018; Lüpold et al., 2016). Perhaps the spermatostyles modulate female mating behavior and are essential for successful fertilization like the anucleate parasperm of some butterflies (Cook and Wedell, 1999; Sakai et al., 2019).

Other insects make sperm conjugates (e.g., Higginson and Pitnick, 2011; Higginson et al., 2012a; 2015), but only carabids and some whirligig beetles are known to make sperm conjugates with spermatostyles (Breland and Simmons, 1970; Gustafson and Miller, 2017; Higginson et al., 2015). Whirligig beetles are close relatives of ground beetles (McKenna et al., 2015; Maddison et al., 2009; Zhang et al., 2018), but the exact phylogenetic position of whirligigs relative to ground beetles is unclear. Whirligigs are usually placed with other aquatic adephagan beetles in a clade that is sister to ground beetles (McKenna et al., 2015; Zhang et al., 2018) or are inferred to be the sister to all remaining Adephaga (Beutel and Roughley, 1988). Evidence from whirligig systematics suggests that the spermatostyle is a derived trait within the group (Gustafson, personal comm.; Gustafson and Miller, 2017; Higginson et al., 2015), which implies that the spermatostyle in carabid beetles and whirligig beetles is convergent.

### Insight into sperm-female interactions

The female reproductive tract can be considered a morphological representation of female sperm preference traits (Birkhead, 1998; Eberhard, 1996), and we found preliminary evidence that females exert pressure on different components of a male’s sperm conjugate. We recovered sperm from the spermathecae of several females, and sperm conjugates clearly arrive at the spermatheca before dissociating (Figs. 12E-J). This observation confirms that sperm compete with spermatostyles for storage within a female’s reproductive tract. This means that a male that invests more in spermatostyle size sacrifices spermathecal space for sperm. Spermatostyles recovered from female preparations were typically thinner or compacted compared to spermatostyles recovered from males’ seminal vesicles, and most conjugated sperm in our female preparations had separated from their cohort sperm (Figs. 12E-J). Because sperm conjugates arrive to the spermatheca but only individual sperm can fertilize eggs, sperm must dissociate at some point (Higginson and Pitnick, 2011). Because sperm dissociation occurs within the female, females may be able to exert control over how quickly a male’s conjugated sperm dissociate (Pitnick et al., 2009b). It is possible that males vary in how easily their conjugated sperm dissociate.

A more compelling piece of evidence for cryptic female choice operating on sperm conjugates in ground beetles is the discovery that some females have separate storage organs for different parts of the sperm conjugate. Some *Galerita* females have a large balloon-shaped organ that stores spermatostyles and a small spherical spermatheca for sperm (Figs. 12A-C). This discovery suggests that female *Galerita* may have partially decoupled sperm evolution from spermatostyle evolution in males. Very little is known regarding sperm use by *Galerita* females, but we suspect that conjugates arrive to the balloon-shaped structure, dissociate or become dissociated from their sperm, and the sperm travel or are moved to the functional spermatheca. Near relatives of *Galerita* also possess a similar configuration of female reproductive tract structures (Hunting, 2008), and it seems likely that this pattern of sperm use applies more broadly.

### Are sperm conjugates greater than the sum of their parts?

Closely related ground beetles generally differ most at the level of the conjugate rather than the sperm themselves. We found variation among ground beetle sperm at multiple levels; in their sperm, spermatostyles, and how these are joined, i.e., the sperm conjugates (Fig. 2; Table 2). Conjugate-level variation likely evolves rapidly, as indicated by the significant variation we observed at this level in our sampling. The sperm of closely related species frequently differ only at the level of the conjugate. If sperm conjugates change rapidly, you would predict to see within-species or within-male variation in sperm conjugates. We found several instances of intraspecies or intramale variation in sperm conjugate form throughout Carabidae (e.g., Fig. 5E), with males making monomorphic sperm but packaging them into sperm conjugates that overlap in size or with males making two sperm conjugate size classes (Takami and Sota, 2007). *Brachinus elongatulus* males make monomorphic sperm but package them into two morphologically distinctive conjugates (Fig. 11A). Conjugate size polymorphism in ground beetles commonly entails variation in spermatostyle size, but we also observed variation beyond conjugate size among closely related species.

Ground beetle conjugates vary between species in the number of included sperm, and conjugates include anywhere from a few to thousands of sperm (Table 2). Because we often had difficulty estimating the number of included sperm in very large conjugates, our data surely underreport the variation that is present in Carabidae. The largest conjugates we observed were all conjugates with large spermatostyles. The density and arrangement of sperm along the spermatostyle determines the number of sperm cells in a given conjugate. As spermatostyles get larger, there is more space for sperm attachment, which may correlate with sperm size and number of sperm embedded in a conjugate. We generally found that larger spermatostyles included more sperm and/or longer sperm, but there are many exceptions.

Variation at the conjugate-level is usually the result of sperm number and distribution patterns rather than orientation of sperm. In some closely related species the conjugates have similar numbers of included sperm but different arrangements of sperm. One such example comes from the sperm conjugates of *Dyschirius tridentatus* and *D. dejeanii.* Both species make long rod-like spermatostyles of similar length with sperm of similar length. However, in *D. tridentatus* about 35 sperm are distributed one at a time throughout the length of the spermatostyle (Figs. 9F-G) whereas in *D. dejeanii* sperm are all located on one end of the spermatostyle in a small cluster of about 7 sperm (Fig. 9H).

A likely functional consequence of variation in number and placement of sperm in a conjugate is variation in motility, but motility alone likely does not explain the variation observed in carabid conjugates (Pitnick et al., 2009a). Evidence from muroid rodents and diving beetles indicates that sperm motility varies with conjugate form (Fisher et al., 2014; Higginson et al., 2012a). Our preliminary data from *in vitro* observation of ground beetle sperm suggests that motility differs among conjugate forms (Supporting Information MV1-MV16). Although we did not systematically investigate this topic enough to warrant firm conclusions, we suspect that sperm conjugate motility is dependent on the composition and arrangement of their sperm in a conjugate (Fisher et al., 2014) as well as interactions with the female reproductive tract epithelium (Lüpold and Pitnick, 2018). Given the wide range of variation observed in conjugate form, it seems unlikely that selection for variation in motility is the only proximate mechanism behind the morphological diversification of ground beetle sperm conjugates.

Our data indicate that the ingredients used to make sperm conjugates likely evolve slower than the arrangement of sperm in a conjugate. This pattern has also been found in other animal groups with a history of sperm conjugation (Higginson and Pitnick, 2011; Higginson et al., 2012a; Immler et al., 2007). The conclusion that sperm and spermatostyles evolve slower than the joining of these two in a conjugate aligns well with research on emergent patterns (Maynard Smith and Szathmary, 1995; Michord, 2007; Parrish and Edelstein-Keshet, 1999; Turing, 1952). Evolutionary theory on emergence predicts that as a group becomes more inclusive, it will show more variation (Novikoff, 1945). The highest levels of organization in emergent patterns contain the most variation (Novikoff, 1945). Our data suggest that there are bottom-up trends in the evolution of emergent patterns where the parts (sperm and spermatostyles) drive change in the group (conjugate) as well as top-down trends where the group drives changes in the parts. Sperm conjugation evolution may follow trends common to other emergent patterns in nature like schooling in fish, flocking in birds, and colony-formation in unicellular organisms (Parrish and Edelstein-Keshet, 1999). If so, the study of sperm conjugation might yield new insights into the evolution and develoμment of complex novel traits.

### Concluding remarks

The variation we observed in ground beetle sperm, spermatostyles, and sperm conjugates hints at rapid evolution at the molecular level. Understanding how genetic and epigenetic networks shape phenotype is a focus in the study of organic evolution (Jablonka and Lamb, 2005) and has previously been identified as a high-priority research goal in the study of post-mating sexual selection (Birkhead and Pizzari, 2002; Lüpold and Pitnick, 2018). As genetic tools are ever improving and becoming more accessible to non-model taxa (e.g., Ellegren, 2014; Russell et al., 2017), studies that use ground beetles and their diverse sperm and genitalia to answer questions on the genetics and epigenetics of post-mating sexual selection will soon be viable.

There are, of course, gaps in our dataset as we were unable to study the sperm form of carabid beetles from every major split in the tree of Carabidae. Sperm morphology of unsampled early diverging carabids such as Southern Hemisphere carabines (beetles of the genus *Pamborus* and *Ceroglossus)* and members of the tribes Migadopini and Cicindini could be particularly valuable for inferring transitions in sperm conjugate evolution. We did not study any male Psydrini, and we were also unable to study the sperm of any Moriomorphini, which are the sister-group of the large clade comprised of Brachininae and Harpalinae (Maddison et al., 2009; Maddison and Ober, 2011). The study of Broscinae sperm may shed light on the transition from sperm conjugation (with a spermatostyle) to singleton sperm (without a spermatostyle). We suspect that the study of sperm in Broscinae worldwide could yield new insights into the intersection of evolution and develoμment of sperm, spermatostyles, and sperm conjugation. In addition, advances in ground beetles phylogenetics from genome and transcriptome sequencing stand to greatly improve the accuracy of our inferences about sperm diversification.

Our survey did not principally focus on female reproductive tract form, and we acknowledge that we focused on only one side of the story (Ah-King et al., 2014). We agree with Ah-King et al., (2014) that future studies that regularly incorporate female reproductive tract data will be essential to a more holistic understanding of morphological evolution in sperm. We look forward to future research that more fully incorporates female reproductive trait data in the study of post-mating sexual selection in ground beetles.

## Supporting information

Spreadsheet S1 SuppInfo

Table S1 SuppInfo

Table S2 SuppInfo

## Acknowledgements

This study is a component of the PhD research of RAG. It is the result of contributions from many people and organizations, and we earnestly hope that we have not forgotten anyone here. For their love and support over many years, RAG gives his deepest appreciation to his parents Roberto Gomez and Yvonne M. Soto-Gomez and his siblings Miguel C. Gomez and Gabriela I. Gomez. For specimens, which served as the bedrock of this study, we would like to thank Curt Harden, Olivia Boyd, John Sproul, Kojun Kanda, and James Pflug. For their camaraderie and assistance in the field near and far, we would like Kojun Kanda, James Pflug, Aaron Smith, Geovanni M. Rodríguez Mirón, Andrew Johnston, Brian Raber, Marcin Kamiński, Daniel and Jutah Bartsch, Ruth Müller, Danielle Mendez, Phil Schapker, Adam Chouinard, James LaBonte, Olivia Boyd, Jason Schaller, Alan Yanahan, John Palting, Colin Schoeman, Jason Denlinger, Dominique Gonçalves, Ana Gledis Miranda, Norina Vicente, the park staff at Parque Nacional da Gorongosa in Mozambique, Lance Morrison, and Dabao Lu. For their help facilitating the collection and/or study of material for this project, we would like to give our thanks to Paulina Cifuentes-Ruiz, Jane Scott, Richard Boyd, the Gomez-Soto family, Danielle Mendez, Alan Yanahan, Jason Schaller, Wendy Moore, Eugene Hall, Jason Denlinger, Piotr Naskrecki and Chris Marshall. For valuable input on taxon sampling, we would like to thank Kip Will and James Liebherr. For sharing their valuable time and taxonomic expertise, we thank Danny Shpeley, Achille Casale, Petr Burlisch, and Kip Will. For help with the staining, imaging, and slide preparation, we would like to thank Barbara Taylor, Anne-Marie Girard, Grey Gustafson, Stephen Baca, and Kelly Miller. For help with SEM and TEM we thank Peter and Rosemary at the Core Labs at OSU. For their friendship and support, RAG thanks the Integrative Biology graduate students. RAG offers deep thanks to his graduate committee at Oregon State University for their mentorship and patience: James Strother, Virginia Weis, Brian Sidlauskas, and Devlin Montfort. Scott Pitnick and Romano Dallai are thanked for sharing their extensive knowledge of sperm biology and their perspectives on the evolution of sperm form in ground beetles.

We gratefully acknowledge the following sources of funding that allowed for us to study carabid beetle sperm from throughout the tree by providing financial assistance for travel for field work: the American Museum of Natural History Theodore Roosevelt Memorial grant (Mexico 2016), Integrative Biology Zoology Research Funds (Florida and Georgia, USA 2016), National Geographic Society early career grant (Republic of South Africa 2018), and the Entomological Society of America Systematics, Biodiversity, and Evolution division student research travel award (Mozambique 2018).

This project was partly funded by a National Science Foundation Graduate Research Fellowship to RAG (DGE-1143953) and by the Harold E. and Leona M. Rice Endowment Fund at Oregon State University.

## Notes

http://morphobank.org/permalink/?P3123

## References

Ah-King, M., Barron, A. B., Herberstein, M. E. (2014). Genital Evolution: Why Are Females Still Understudied? PLoS Biology, 12: e1001851. doi: https://doi.org/10.1371/journal.pbio.1001851

Ardnt, E., Beutel, R. G., Will, K. (2005). 7.8. Carabidae Latreille, 1802. In Beutel, R.G., Leschen, R.A.B. (Eds.), Handbook of Zoology. Band/Volume IV Arthropoda: Insecta. Teilband/Part 38 Coleoptera, Beetles. Volume 1: Morphology and Systematics (Archostemata, Adephaga, Myxophaga, Polyphaga partim) (pp. 119–146). Walter de Gruyter: Berlin, New York.

Arnqvist, G., Rowe, L. (2005). Sexual Conflict. Princeton University Press, Princeton.

Beutel, R. G., Roughley, R. E. (1988). On the systematic position of the family Gyrinidae (Coleotpera: Adephaga). Journal of Zoological Systematics and Evolutionary Research, 26: 380–400.

Birkhead, T. R. (1998). Cryptic female choice: Criteria for estabilishing female sperm choice. Evolution, 52:1212–1218.

Birkhead, T. R., Montgomerie, R. (2009). Three centuries of sperm research. In Birkhead, T.R., Hosken, D.J., Pitnick, S. (Eds.), Sperm Biology, an Evolutionary Perspective (pp. 1–42). Academic Press, San Diego.

Birkhead, T. R., Pizzari, T. (2002). Postcopulatory sexual selection. Nature Reviews and Genetics, 3: 262–273.

Bouix, G. (1961). Atypisme et degenerescence des spermatozoids dans le genre *Carabus*. Comptes Rendus De L’Academie Des Sciences, 252: 329–330.

Bouix, G. (1963). Sur la spermatogenese des *Carabus*, modalite et frequence de la spermiogenese atypique. Comptes Rendus De L’Academie Des Sciences, 256: 2698–2701.

Bousquet, Y. (2012). Catalogue of Geadephaga (Coleoptera, Adephaga) of America, north of Mexico. Zookeys, 245:1–1722.

Breland, O. P., Simmons, E. (1970). Preliminary studies of the spermatozoa and the male reproductive system of some whirligig beetles (Coleoptera: Gyrinidae). Entomological News, 81: 101–110.

Cook, P. A., Wedell, N. (1999). Non-fertile sperm delay female remating. Nature, 397: 486.

Crowson, R. A. (1981). The biology of the Coleoptera. Academic Press, London, 802 pp.

Dallai, R., Mercati, D., Giglio, A., Lupetti, P. (2019). Sperm ultrastructure in several species of Carabidae beetles (Insecta, Adephaga) and their organization in spermatozeugmata. Arthropod Structure & Development, 51: 1–13.

Darwin, C. (1871). The descent of man and selection in relation to sex. Reprinted, New York: Modern Library.

Eberhard, W. G. (1985). Sexual Selection and Animal Genitalia. Harvard University Press, Cambridge, MA.

Eberhard, W. G. (1996). Female control: sexual selection by cryptic female choice. Princeton University Press, Princeton, NJ.

Ellegren, H. (2014). Genome sequencing and population genomics in non-model organisms. Trends in Ecology & Evolution, 29: 51–63.

Ferenz, H-J. (1986). Structure and formation of sperm bundles in the carabid beetle *Pterostichus nigrita*. In den Boer, P.J., Luff, M.L., Mossokowski, D., Weber, F. (Eds.), Carabid beetles, their adaptations and dynamics (pp. 147–155). Gustav Fischer, Stuttgart, New York.

Ferraguti, M., Grassi, G., Erséus, C. (1989). Different models oftubificid spermatozeugmata. Hydrobiologia, 180: 73–82.

Fisher, H. S., Giomi, L, Hoekstra, H. E., Mahadevan, L. (2014). The dynamics of sperm cooperation in a competitive environment. Proceedings of Royal Society. Series B, Biological sciences, 281: 20140296.

Gilson, G. (1884). Etude compareé de la spermatogénèse chez les arthropods. La Cellule. I: 84–94.

Gough, H. M., Duran, D. P., Kawahara, A. Y., Toussaint, E. F. A. (2018). A comprehensive molecular phylogeny of tiger beetles (Coleoptera, Carabidae, Cicindelinae). Systematic Entomology, 44: 305–321.

Gustafson, G. T., Miller, K. B. (2017). Systematics and evolution of the whirligig beetle tribe Dineutini (Coleoptera: Gyrinidae: Gyrininae). Zoological Journal of the Linnean Society, XX: 1–33.

Higginson, D. M., Pitnick, S. (2011). Evolution of intra-ejaculate sperm interactions: do sperm cooperate? Biological Reviews, 86: 249–270.

Higginson, D. M., Miller, K. B., Segraves, K. A., Pitnick, S. (2012). Convergence, recurrence and diversification of complex sperm traits in diving beetles (Dytiscidae). Evolution, 66:1650–1661.

Higginson, D. M., Miller, K. B., Segraves, K. A., Pitnick, S. (2012) Female reproductive tract form drives the evolution of complex sperm morphology. Proceedings of the National Academy of Sciences, 109: 4538–4543.

Higginson, D. M., Badyaev, A. V., Segraves, K. A., Pitnick, S. (2015) Causes of discordance between allometries at and above species-level: an example with aquatic beetles. The American Naturalist, 186:176–186. https://doi.org/10.1086/682049

Hinchliff, C. E., Smith, S. A., Allman, J. F., Burleigh, J. G., Chaudhary, R., Coghilll, L. M., Crandall, K. A., Deng, J., Drew, B. T., Gazis, R., Gude, K., Hibbett, D. S., Katz, L. A., Laughinhouse, H. D., McTavish, E. J., Midford, P. E., Owen, C. L., Ree, R. H., Rees, J. A., Soltis, D. E., Williams, T., Cranston, K. A. (2015). Synthesis of phylogeny and taxonomy into a comprehensive tree of life. Proceedings of the National Academy of Sciences, 112: 12764–12769. https://doi.org/10.1073/pnas.1423041112

Hodgson, A. N., Ferenz, H-J., Schneider, S. (2013). Formation of sperm bundles in *Pterostichus nigrita* (Coleotpera: Carabidae). Invertebrate Reproduction & Development, 57:120–131.

Holman, L., Snook, R. R. (2006). Spermicide, cryptic female choice and the evolution of sperm form and function. Journal of Evolutionary Biology, 19:1660–1670.

Hosken, D. J., Stockley, P. (2004). Sexual selection and genital evolution. Trends in Ecology and Evolution, 19: 87–93.

Humphries, S., Evans, J. P. (2008). Sperm competition: linking form to function, BMC Evolutionary Biology, 8: 319.

Hunting, W. (2008) Female reproductive system of the tribe Galeritini (Coleoptera: Carabidae): structural features and evolution. Annals of Carnegia Museum, 77(1): 229–242 doi: 10.2992/0097-4463-77.1.229

Immler, S., Moore, H. D., Breed, W. G., Birkhead, T. R. (2007). By Hook or by Crook? Morphometry, Competition and Cooperation in Rodent Sperm. PLoS ONE 2(1): e 170. https://doi.org/10.1371/journal.pone.0000170

Ishimoto, K., & Gaffney, E. A. (2015). Fluid flow and sperm guidance: a simulation study of hydrodynamic sperm rheotaxis. Journal of the Royal Society, Interface, 12(106): 20150172.

Jablonka, E., Lamb, M. (2006). Evolution in Four Dimensions, Cambridge, MA: MIT Press.

Jeannel, R. (1941). Coléoptères Carabiques. Faune de France Part 1, 39:1–571.

Liebherr, J. K., Will, K. W. (1998). Inferring phylogenetic relationships within Carabidae (Insecta, Coleoptera) from characters of the female reproductive tract. In Ball, G., Casale, A., Vigna Taglianti, A. (Eds.), Phylogeny and Classification of Caraboidea (Coleoptera: Adephaga) (pp. 107-170). Museo Regionale di Scienze Naturali, Torino.

Lorenz, W. (2005). Systematic list of extant ground beetles of the world (Insecta Coleoptera “Geadephaga”: Trachypachidae and Carabidae incl. Paussinae, Cicindelinae, Rhysodinae). Second edition. [Author], Titzing, Germany

Lorenz, W. (2018). CarabCat: Global database of ground beetles (version Oct 2017). In Roskov Y., Ower G., Orrell T., Nicolson D., Bailly N., Kirk P.M., Bourgoin T., DeWalt R.E., Decock W., De Wever A., Nieukerken E. van, Zarucchi J., Penev L, eds. (2018), Species 2000 & ITIS Catalogue of Life, 24th September 2018. Digital resource at www.catalogueoflife.org/col. Species 2000: Naturalis, Leiden, the Netherlands. ISSN 2405-8858.

Lüpold, S., Manier, M. K., Puniamoorthy, N., Schoff, C., Starmer, W. T., Buckley Luepold, S. H., Belote, J. M., Pitnick, S. (2016). How sexual selection can drive the evolution of costly sperm ornamentation. Nature, 533: 535–538.

Lüpold, S., Pitnick, S. (2018). Sperm form and function: what do we know about the role of sexual selection. Reproduction, 155: 229–243.

Maddison, D. R., Baker, M. D., Ober, K. A. (1999). Phylogeny of carabid beetles as inferred from 18S ribosomal DNA (Coleoptera: Carabidae). Systematic Entomology, 24:103–138.

Maddison, D. R., Ober, K. A. (2011). Phylogeny of minute carabid beetles and their relatives based upon DNA sequence data (Coleoptera, Carabidae, Trechitae). ZooKeys, 147: 229–260.

Maddison, D. R., Kanda, K., Boyd, O. F., Faille, A., Porch, N., Erwin, T. L., Roig-Juñent, S. (2019). Phylogeny of the beetle supertribe Trechitae (Coleoptera: Carabidae): Unexpected clades, isolated lineages, and morphological convergence. Molecular phylogenetics and Evolution, 132:151–176.

Maddison, D. R., Moore, W., Baker, M. D., Ellis, T. M., Ober, K. A., Cannone, J. J., Gutell, R. R. (2009). Monophyly of terrestrial adephagan beetles as indicated by three nuclear genes (Coleoptera: Carabidae and Trachypachidae). Zoologica Scripta, 38:43–62.

Martínez-Navarro, E. M., Galián, J., Serrano, J. (2005). Phylogeny and molecular evolution of the tribe Harpalini (Coleoptera, Carabidae) inferred from mitochondrial cytochrome-oxidase I. Molecular Phylogenetics and Evolution, 35:127–146.

Maynard Smith, J., Szathmary, E. (1995). The major transitions in evolution. Oxford University Press.

McCloy, R. A., Rogers, S., Caldon, C. E., Lorca, T., Castro, A., Burgess, A. (2014). Partial inhibition of Cdkl in G 2 phase overrides the SAC and decouples mitotic events. Cell Cycle, 13:1400–1412.

McKenna, D. D., Wild, A. L., Kanda, K., Bellamy, C. L., Beutel, R. G., Caterino, M. S., Farnum, C. W., Hawks, D. C., Ivie, M. A., Jameson, M. L, Leschen, R. A. B., Marvaldi, A. E., McHugh, J. V., Newton, A. F., Robertson, J. A., Thayer, M. K., Whiting, M. F., Lawrence, J. F., Slipihski, A., Maddison, D. R., Farrell, B. D. (2015) The beetle tree of life reveals that Coleoptera survived end-Permian mass extinction to diversify during the Cretaceous terrestrial revolution. Systematic Entomology, 40: 835–880.

Michord, R.E. (2007). Evolution of individuality during the transition from unicellular to multicellular life. Proceedings of the National Academy of Sciences, 104: 8613–8618.

Miller, K. B., Bergsten, J. (2014). Predaceous diving beetle sexual systems. In Yee, D.A. (Ed.), Ecology, Systematics, and Natural History of Predaceous Diving Beetles (Coleoptera: Dytiscidae) (pp. 199–234). Springer, New York.

Miller, G. T., Pitnick, S. (2002). Sperm-female coevolution in *Drosophila*. Science, 298: 1230–1233.

Moore, H., Dvoráková, K., Nicholas, J., Breed, W. (2002). Exceptional sperm cooperation in the wood mouse. Nature, 418:174–177.

Moore, W. (2008). Phylogeny of Western Hemisphere Ozaenini (Coleoptera: Carabidae: Paussinae) based on DNA sequence data. Annals of Carnegie Museum, 77: 79–92.

Nichols, S. (1988). Systematics and biogeography of West Indian Scaritinae (Coleoptera: Carabidae). (Unpublished Ph.D. thesis). Ithaca, New York: Cornell University.

Novikoff, A. B. (1945). The concept of integrative levels and biology. Science, 101: 209–215.

Ober, K. A. (2002). Phylogenetic relationships of the carabid subfamily Harpalinae (Coleoptera) based on molecular sequence data. Molecular Phylogenetics and Evolution, 24: 228–248.

Ober, K. A., Maddison, D. R. (2008). Phylogenetic relationships of tribes within Harpalinae (Coleoptera: Carabidae) as inferred from 28S ribosomal DNA and the wingless gene. Journal of Insect Science, 8:1–32.

Osawa, S., Su, Z.-H., Imura, Y. (2004). Phylogeny and distribution of the subfamily carabinae. In Osawa, S., Su, Z.-H., Imura, Y. (Eds.), Molecular phylogeny and evolution of carabid ground beetles (pp. 25–32). Tokyo: Springer.

Parker, G. A. (1970). Sperm competition and its evolutionary consequences. Biological Reviews, 45: 525–567.

Parker, G. A. (1979). Sexual selection and sexual conflict. In Blum, M.S., Blum, N.A. (Eds.), Sexual Selection and Reproductive Competition in Insects (pp. 123­166). Academic Press, London.

Parker, G. A. (1998). Sperm competition and the evolution of ejaculates: towards a theory base. In Birkhead, T.R., Moller, A.P. (Eds.), Sperm Competition and Sexual Selection (pp. 3–54). Academic Press, San Diego.

Parker, G. A. (2005). Sexual conflict over mating and fertilization: an overview. Philosophical Transactions of the Royal Society B, 361: 235–259.

Parrish, J. K., Edelstein-Keshet, L. (1999). Complexity, Pattern, and Evolutionary Trade-Offs in Animal Aggregation. Science, 284: 99–101.

Pitnick, S. (1996). Investment in Testes and the Cost of Making Long Sperm in *Drosophila*. The American Naturalist, 148: 57–80.

Pitnick, S., Hosken, D. J. (2010). Postcopulatory Sexual Selection. In Westneat, D.F., Fox, C.W. (Eds.), Evolutionary Behavioral Ecology (pp. 379–399). Oxford University Press, New York.

Pitnick, S., Hosken, D. J., Birkhead, T. R. (2009). Sperm morphological diversity. In Birkhead, T.R., Hosken, D.J., Pitnick, S. (Eds.), Sperm Biology, an Evolutionary Perspective (pp. 69–149). Academic Press, San Diego.

Pitnick, S., Wolfner, M. F., Suarez, S. (2009). Ejaculate-female and sperm-female interactions. In Birkhead, T.R., Hosken, D.J., Pitnick, S. (Eds.), Sperm Biology, an Evolutionary Perspective (pp. 247–304). Academic Press, San Diego.

Pizzari, T., Foster, K. R. (2008). Sperm Sociality: Cooperation, Altruism, and Spite. PLoS Biol 6(5): el30. https://doi.org/10.1371/journal.pbio.0060130

Rasband, W. S. (2012). ImageJ, version 1.43u. National Institutes of Health, Bethesda, MD.

Robertson, J. A., Moore, W. (2016). Dissecting the species groups of *Paussus* L. (Carabidae: Paussinae): unraveling morphological convergence associated with myrmecophilous life histories. Systematic Entomology. DOI: 10.111/sysen.l2205

Russell, J. J., Theriot, J. A., Sood, P., Marshall, W. F., Landweber, L. F., Fritz-Laylin, L., Polka, J. K., Oliferenko, S., Gerbich, T., Gladfelter, A., Umen, J., Bezanilla, M., Lancaster, M. A., He, S., Gibson, M. C., Goldstein, B., Tanaka, E. M., Hu, C.-K., Brunet, A. (2017). Non-model model organisms. BMC Biology, 15: 55.

Sakai, H., Oshima, H., Yuri, K., Gotoh, H., Daimon, T., Yaginuma, T., Sahara, K., Niimi, T. (2019). Dimorphic sperm formation by *Sex-lethal*. Proceedings of the National Academy of Sciences, 116: 10412–10417.

Sasakawa, K. (2007). Sperm bundle and reproductive organs of carabid beetles tribe Pterostichini (Coleoptera: Carabidae). Naturwissenschaften, 94: 384–391.

Sasakawa, K., Toki, W. (2008). A new record, sperm bundle morphology and preliminary data on the breeding type of the ground beetle *Jujiroa estriata* Sasakawa (Coleoptera: Carabidae: Platynini). Entomological Science, 11:415­417.

Sasakawa, K. (2009). Marked sperm dimorphism in the ground beetle *Scarites terricola:* a novel type of insect sperm polymorphism. Physiological Entomology, 34: 387–390.

Schärer, L, Littlewood, D. T. J., Waeschenbach, A., Yoshida, W., Vizoso, D. B. (2011). Mating behavior and the evolution of sperm design. Proceedings of the National Academy of Sciences, 108: 1490–1495.

Schmoll, T., Sanciprian, R., Klevne, O. (2016). No evidence for effects of formalin storage duration or solvent medium exposure on avian sperm morphology. Journal of Ornithology, 157: 647.

Schubert, L. F., Krüger, S., Moritz, G. B., Schubert, V. (2017). Male reproductive system and spermatogenesis of *Limodromus assimilis* (Paykull 1790). PLoS ONE, 12: e0180492. https://doi.org/10.1371/journal.pone.0180492

Sivinski, J. (1984). Sperm in competition. In R.L. Smith (Ed.), Sperm Competition and the Evolution of Animal Mating Systems (pp. 85–115). Academic Press, New York.

Taggart, D. A., Johnson, J. L, O’Brien, H. P., Moore, H. D. M. (1993). Why do spermatozoa of American Marsupials form pairs? A clue from the analysis of sperm-pairing in the epididymis of the Grey Short-Tailed Opossum, *Monodelphis domestica*. The Anatomical Record, 236: 465–478.

Takami, Y., Sota, T. (2007). Sperm competition promotes diversity of sperm bundles in *Ohomopterus* ground beetles. Naturwissenschaften, 94: 543–550.

Turing, A. M. (1952). The Chemical Basis of Morphogenesis. Philosophical Transactions of the Royal Society of London. Series B, Biological Sciences, 237: 37–72.

Thornhill, R., Alcock, J. (1983). The evolution of insect mating systems. Harvard University Press, Cambridge, MA.

van der Horst, G., Maree, L. (2010). SpermBlue®: A new universal stain for human and animal sperm which is also amenable to automated sperm morphology analysis, Biotechnic & Histochemistry, 84(**6**): 299–308. doi: 10.3109/10520290902984274

Vogel, S. (1994). Life in Moving Fluids: the Physical Biology of Flow. Princeton University Press.

Vogler, A. P., Pearson, D. L. (1996). A molecular phylogeny of tiger beetles (Cicindelinae): congruence of mitochondrial and nuclear rDNA data sets. Molecular Phylogenetics and Evolution, 6: 321–328.

Werner, G. (1965). Untersuchungen über Die Spermiogenese Beim Sandläufer, *Cicindela campestris* L.Z. *Zeitschriftfür Zellforschung und Mikroskopische Anatomie*. Abteilung Histochemie, 66: 255–275.

Will, K. W., Liebherr, J. K., Maddison, D. R., Galián, J. A. (2005). Absence asymmetry: the evolution of monorchid beetles (Insecta: Coleoptera: Carabidae). Journal of morphology, 264: 75–93.

Will, K. W., Gill, A. S. (2008). Phylogeny and classification of *Hypherpes auctorum* (Coleoptera: Carabidae: Pterostichini: *Pterostichus)*. Annals of Carnegie Museum, 77: 93–127.

Witz, B. W. (1990). Comparative ultrastructural analysis of spermatogenesis in *Pasimachus subsulcatus* and *P. strenuus* (Coleoptera: Carabidae). Invertebrate Reproduction and Develoμment, 18:197–203.

Zhang, S.-Q., Che, L.-H., Li, Y., Liang, D., Pang, H., Ślipiński, A., Zhang, P. (2018). Evolutionary history of Coleoptera revealed by extensive sampling of genes and species. Nature communications, 9: 205.

