## Supplementary material for "Novelty and emergent patterns in sperm: morphological diversity and evolution of spermatozoa and sperm conjugation in ground beetles (Coleoptera: Carabidae)": Table S1 SuppInfo

**Table S1**: A brief summary of the state of knowledge of carabid sperm form in the primary literature.

| **Species** | **Sperm polymorphism (0 absent/**  **1 present)** | **Conjugation (0 absent/**  **1 present)** | **Data reported^1^** | **References** |
| --- | --- | --- | --- | --- |
| **Carabinae**  **Carabini** |  |  |  |  |
| *Calosoma inquisitor* (Linnaeus, 1758) | 0 | 1 | AS, CS | Gilson (1884) |
| *Carabus* Linnaeus, 1758 spp. [8 species?] | 1 | ? | FL?, HL? | Bouix (1961,1963) |
| *Carabus (Archicarabus) alysidotus*^2^ Illiger, 1798 | 1 | ? | - | Bouix (1963) |
| *Carabus (Carabus) vanvolxemi* Putzeys, 1785 | 1 | 1 | RL, SL | Takami and Sota (2007) |
| *Carabus (Carabus) granulatus* Linnaeus, 1758 | 0 | 1 | AS, CS, SL, SU | Dallai et al. (2019) |
| *Carabus (Chrysocarabus) auronitens* Fabricius, 1792 | 1 | ? | 1^st^ form: ?  2^nd^ form: HL | Bouix (1961,1963); Gilson (1884) |
| *Carabus (Chrysocarabus) basilicus*^3^ Chevrolat, 1836 | 1 | ? | - | Bouix (1963) |
| *Carabus (Chrysocarabus) hispanus*^2^ Fabricius, 1787 | 1 | ? | - | Bouix (1963) |
| *Carabus (Chrysocarabus) punctatoauratus* Germar, 1824 | 1 | ? | - | Bouix (1963) |
| *Carabus (Chyrsocarabus) rutilans*^2^ Dejean, 1826 | 1 | ? | - | Bouix (1963) |
| *Carabus (Mesocarabus) lusitanicus*^3^ Fabricius, 1801 | 1 | ? | 1^st^ form: ?  2^nd^ form: HL | Bouix (1963) |
| *Carabus (Megodontus) purpurascens* Fabricius, 1787 | 0 | 1 | AS, CS | Gilson (1884) |
| *Carabus (Ohomopterus) arrowianus* (Breuning, 1934) | 0 | 1 | RL, SL | Takami and Sota (2007) |
| *Carabus (Ohomopterus) dehaanii* Chaudoir, 1848 | 0 | 1 | RL, SL | Takami and Sota (2007) |
| *Carabus (Ohomopterus) insulicola* Chaudoir, 1869 | 0 | 1 | RL, SL | Takami (2002); Takami and Sota (2007) |
| *Carabus (Ohomopterus) iwawakianus* (Nakane, 1953) | 0 | 1 | RL, SL | Takami and Sota (2007) |
| *Carabus (Ohomopterus) kimurai* (Ishikawa, 1966) | 0 | 1 | RL, SL | Takami and Sota (2007) |
| *Carabus (Ohomopterus) maiyasanus* Bates, 1873 | 0 | 1 | RL, SL | Takami and Sota (2007) |
| *Carabus (Ohomopterus) uenoi* (Ishikawa, 1960) | 0 | 1 | RL, SL | Takami and Sota (2007) |
| *Carabus (Ohomopterus) yamato* (Nakane, 1953) | 0 | 1 | RL, SL | Takami and Sota (2007) |
| *Carabus (Oreocarabus) preslii* Dejean, 1830 | 0 | 1 | AS, CS, RL, SL, SU | Dallai et al. (2019) |
| *Carabus (Procrustes) coriaceus* Linnaeus, 1758 | 0 | 1 | AS, CS | Gilson (1884) |
| *Carabus* (*Tachypus*) *auratus* Linnaeus, 1758 | 0 | 1 | AS, CS | Gilson (1884) |
| **Cicindelinae**  **Cicindelini** |  |  |  |  |
| *Cicindela campestris* Linnaeus, 1758 | 0 | 0 | SU | Werner (1965) |
| **Harpalinae** |  |  |  |  |
| **Lebiini** |  |  |  |  |
| *Demetrias atricapillus* (Linnaeus, 1758) | 0 | 1 | AS, CS, SU | Dallai et al. (2019) |
| **Loricerinae**  **Loricerini** |  |  |  |  |
| *Loricera* Latreille, 1802 sp. | 0 | 1 | AS, CS | Gilson (1884) |
| **Platynini** |  |  |  |  |
| *Jujiroa estriata* Sasakawa, 2006 | 0 | 1 | AS, CL, CS | Sasakawa and Toki (2008) |
| *Limodromus assimilis* (Paykull, 1790) | 0 | 1 | AS, CS, RL, SL | Schubert et al. (2017) |
| **Pterostichini** |  |  |  |  |
| *Amara aulica* (Panzer, 1796) | 0 | 1 | AS, CS, SU | Dallai et al. (2019) |
| *Lesticus magnus* (Motschulsky, 1860) | 0 | 1 | AS, CL, CS | Sasakawa (2007) |
| *Myas cuprescens* (Motschulsky, 1858) | 0 | 1 | AS, CL, CS | Sasakawa (2007) |
| *Percus strictus* Dejean, 1828 | 0 | 1 | AS, CS | Carpucino et al. (2002) |
| *Poecilus samurai* (Lutshnik, 1916) | 0 | 1 | AS, CL, CS | Sasakawa (2007) |
| *Poecilus versicolor* (Sturm, 1824) | 0 | 1 | AS, CL, CS | Sasakawa (2007) |
| *Pterostichus* (*Argutor*) *dulcis* (Bates, 1883) | 0 | 1 | AS, CL, CS | Sasakawa (2007) |
| *Pterostichus* (*Argutor*) *sulcitarsis* Morawitz, 1862 | 0 | 1 | AS, CL, CS | Sasakawa (2007) |
| *Pterostichus* (*Badistrinus*) *bandotaro* Tanaka, 1958 | 0 | 1 | AS, CL, CS | Sasakawa (2007) |
| *Pterostichus (Bothriopterus) adstrictus* Eschscholtz, 1823 | 0 | 1 | AS, CL, CS | Sasakawa (2007) |
| *Pterostichus (Bothriopterus) subovatus* (Motschulsky, 1860) | 0 | 1 | AS, CL, CS | Sasakawa (2007) |
| *Pterostichus (Eosteropus) fuligineus* Morawitz, 1862 | 0 | 1 | AS, CL, CS | Sasakawa (2007) |
| *Pterostichus (Eosteropus) karasawai* Tanaka, 1958 | 0 | 1 | AS, CL, CS | Sasakawa (2007) |
| *Pterostichus (Eosteropus) orientalis* (Motschulsky, 1844) | 0 | 1 | AS, CL, CS | Sasakawa (2007) |
| *Pterostichus (Eosteropus) prolongatus* Morawitz, 1862 | 0 | 1 | AS, CL, CS | Sasakawa (2007) |
| *Pterostichus (Euferonia) habui* Jedlička, 1962 | 0 | 1 | AS, CL, CS | Sasakawa (2007) |
| *Pterostichus (Euferonia) thunbergi* Morawitz, 1862 | 0 | 1 | AS, CL, CS | Sasakawa (2007) |
| *Pterostichus (Eurythoracana) haptoderoides* (Tschitschérine, 1889) | 0 | 1 | AS, CL, CS | Sasakawa (2007) |
| *Pterostichus (Eurythoracana) kajimurai* Habu and Tanaka, 1957 | 0 | 1 | AS, CL, CS | Sasakawa (2007) |
| *Pterostichus (Georgeballius) hoplites* (Bates, 1883) | 0 | 1 | AS, CL, CS | Sasakawa (2007) |
| *Pterostichus (Japeris) defossus* Bates, 1883 | 0 | 1 | AS, CL, CS | Sasakawa (2007) |
| *Pterostichus (Lianoe) mirificus* Bates, 1883 | 0 | 1 | AS, CL, CS | Sasakawa (2007) |
| *Pterostichus (Lyrothorax) amagisanus* Tanaka and Ishida, 1972 | 0 | 1 | AS, CL, CS | Sasakawa (2007) |
| *Pterostichus (Lyrothorax) yoritomus* Bates, 1873 | 0 | 1 | AS, CL, CS | Sasakawa (2007) |
| *Pterostichus* (*Melanius*) *aterrimus* (Herbst, 1784) [unsure if =*Feronea nigerrima* [sic]; Gilson (1884)] | 0 | 1 | AS, CS | Gilson (1884) |
| *Pterostichus (Melanius) noguchii* Bates, 1873 | 0 | 1 | AS, CL, CS | Sasakawa (2007) |
| *Pterostichus (Morphnosoma) melanarius* (Illiger, 1798) | 0 | 1 | AS, CS, SL, SU | Dallai et al. (2019) |
| *Pterostichus (Nialoe) tokejii* Yoshida and Tanaka, 1960 | 0 | 1 | AS, CL, CS | Sasakawa (2007) |
| *Pterostichus (Oreophilus) morio* (Duftschmid, 1812) | 0 | 1 | AS, CS, SL, SU | Dallai et al. (2019) |
| *Pterostichus (Phonias) longinquus* Bates, 1873 | 0 | 1 | AS, CL, CS | Sasakawa (2007) |
| *Pterostichus (Platysma) eschscholtzii (*Germar, 1824) | 0 | 1 | AS, CL, CS | Sasakawa (2007) |
| *Pterostichus (Pseudomaseus) anthracinum* (Illiger, 1798) [unsure if = *Feronea anthracina* [sic] Gilson (1884)] | 0 | 1 | AS, CS | Gilson (1884) |
| *Pterostichus nigrita* (Paykull, 1790) | 0 | 1 | AS, CS, CL, SL, SU | Ferenz (1986); Hodgson et al. (2013); Schneider and Ferenz (2012) |
| *Pterostichus (Pseudomaseus) rotundangulus* Morawitz, 1862 | 0 | 1 | AS, CL, CS | Sasakawa (2007) |
| *Pterostichus (Rhagadus) microcephalus* (Motschulsky, 1860) | 0 | 1 | AS, CL, CS | Sasakawa (2007) |
| *Pterostichus (Rhagadus) polygenus* Bates, 1883 | 0 | 1 | AS, CL, CS | Sasakawa (2007) |
| *Pterostichus (Rhagadus) takaosanus* Habu, 1958 | 0 | 1 | AS, CL, CS | Sasakawa (2007) |
| *Pterostichus (Steropus) melas* (Creutzer, 1799) | 0 | 1 | AS, CS, SL, SU | Dallai et al. (2019) |
| **Scaritinae** ***sensu lato***  **Scaritini** |  |  |  |  |
| *Pasimachus subsulcatus* Say, 1823 | 0 | 1 | SL, SU | Witz (1990) |
| *Pasimachus strenuus* LeConte, 1874 | 0 | 1 | SU | Witz (1990) |
| *Scarites terricola* Bonelli, 1813 | 1 | 1^st^ form: 1  2^nd^ form: 0 | 1^st^ form: AS, CL, CS  2^nd^ form: FL, HL, HW, SL | Sasakawa (2009) |

^1^ Attachment of sperm to spermatostyle or hyaline material (AS): whether the flagella are bounded by additional material or are unbounded. Conjugate length (CL): length of the conjugate with sperm attached, which may or may not be equal to length of a spermatozoon. Conjugate shape (CS): some aspect of conjugate shape reported, often whether the conjugate possesses a left- or right-handed spiral, etc. Flagellum length (FL). Head length (HL). Head width (HD). Rod length (RL): length of spermatostyle or hyaline cap ignoring attached sperm. Sperm length (SL). Sperm ultrastructure (SU).

^2^Now placed in a different subgenus.

^3^The name that was used in the original publication is a junior synonym, and the name here reflects the current classification.
