## Supplementary material for "Novelty and emergent patterns in sperm: morphological diversity and evolution of spermatozoa and sperm conjugation in ground beetles (Coleoptera: Carabidae)": Table S2 SuppInfo

**Table S2**. Locality data for specimens sampled. Abbreviated specimen codes are reported below. Complete specimen codes include each number shown below preceded by the string “RAGspcmn0000000”.

| **species** | **code** | **locality** |
| --- | --- | --- |
| *Abacetus* sp. | 576 | RSA: KwaZulu-Natal: Royal Natal National Park, 28.687°S 28.9539°E, 1404m. 18.i.2018. |
| *Abacetus* sp. | 578 | RSA: KwaZulu-Natal: Royal Natal National Park, 28.687°S 28.9539°E, 1404m. 18.i.2018. |
| *Abacetus* sp*.* | 615 | RSA: KwaZulu-Natal: Hluhluwe-iMfolozi Park, Bekapanzi Pan, 28.2795°S 31.8203°E, 151m. 29.i.2018. |
| *Abacetus* sp. | 616 | RSA: KwaZulu-Natal: Hluhluwe-iMfolozi Park, Bekapanzi Pan, 28.2795°S 31.8203°E, 151m. 29.i.2018. |
| *Abaris splendidula* | 390 | USA: AZ: Cochise Co., San Pedro River, 31.5501°N 110.1375°W, 1237m. 09.viii.2016. |
| *Abaris splendidula* | 396 | USA: AZ: Cochise Co., San Pedro River, 31.5501°N 110.1375°W, 1237m. 09.viii.2016. |
| *Abaris splendidula* | 398 | USA: AZ: Santa Cruz Co., Santa Cruz River, Santa Gertrudis Lane, 31.5621°N 111.0458°W. 10.viii.2016. |
| *Agonum muelleri* | 172 | USA: OR: Benton Co., Corvallis, Crystal Lake Sports Complex, 44.5497°N 123.2453°W. 09.iv.2016. |
| *Agonum muelleri* | 217 | USA: OR: Benton Co., Corvallis, Crystal Lake Sports Complex, 44.5497°N 123.2453°W. 09.iv.2016. |
| *Agonum piceolum* | 147 | USA: OR: Benton Co., Corvallis, Crystal Lake Sport Complex, 44.5497°N 123.2478°W. 29.iii.2016. |
| *Agonum piceolum* | 148 | USA: OR: Benton Co., Corvallis, Crystal Lake Sport Complex, 44.5497°N 123.2478°W. 29.iii.2016. |
| *Agonum piceolum* | 150 | USA: OR: Benton Co., Corvallis, Crystal Lake Sport Complex, 44.5497°N 123.2478°W. 29.iii.2016. |
| *Agonum piceolum* | 151 | USA: OR: Benton Co., Corvallis, Crystal Lake Sport Complex, 44.5497°N 123.2478°W. 29.iii.2016. |
| *Agonum piceolum* | 152 | USA: OR: Benton Co., Corvallis, Crystal Lake Sport Complex, 44.5497°N 123.2478°W. 29.iii.2016. |
| *Agra* sp. 1 | 107 | Mexico: Veracruz: Est. Biol. Los Tuxtlas, 18.5855°N 95.0752°W, 125m. 06-09.viii.2015. |
| *Agra* sp. 2 | 123 | Mexico: Chiapas: La Sepultura, road to Madre Mia, 16.1691°N 93.5087°W, 786m. 11.viii.2015. |
| *Akephorus obesus* | 239 | USA: OR: Lincoln Co., Moolack Beach, 44.7093°N 124.0605°W, 4m. 21.v.2016. |
| *Akephorus obesus* | 243 | USA: OR: Lincoln Co., Moolack Beach, 44.7093°N 124.0605°W, 4m. 21.v.2016. |
| *Akephorus obesus* | 244 | USA: OR: Lincoln Co., Moolack Beach, 44.7093°N 124.0605°W, 4m. 21.v.2016. |
| *Akephorus obesus* | 282 | USA: OR: Lincoln Co., Moolack Beach, 44.7093°N 124.0605°W, 4m. 21.v.2016. |
| *Akephorus obesus* | 466 | USA: OR: Lincoln Co., Moolack Beach, 44.7017°N 124.0621°W. 03.xi.2016. |
| *Amara aenea* | 170 | USA: OR: Benton Co., Corvallis, Crystal Lake Sports Complex, 44.5497°N 123.2453°W, 09.iv.2016. |
| *Amara aenea* | 171 | USA: OR: Benton Co., Corvallis, Crystal Lake Sports Complex, 44.5497°N 123.2453°W, 09.iv.2016. |
| *Amara aenea* | 173 | USA: OR: Benton Co., Corvallis, Crystal Lake Sports Complex, 44.5497°N 123.2453°W, 09.iv.2016. |
| *Amara aenea* | 174 | USA: OR: Benton Co., Corvallis, Crystal Lake Sports Complex, 44.5497°N 123.2453°W, 09.iv.2016. |
| *Amara aenea* | 175 | USA: OR: Benton Co., Corvallis, Crystal Lake Sports Complex, 44.5497°N 123.2453°W, 09.iv.2016. |
| *Amara aenea* | 189 | USA: OR: Benton Co., Corvallis, Crystal Lake Sports Complex, 44.5497°N 123.2453°W, 09.iv.2016. |
| *Amara aenea* | 190 | USA: OR: Benton Co., Corvallis, Crystal Lake Sports Complex, 44.5497°N 123.2453°W, 09.iv.2016. |
| *Amara aenea* | 313 | USA: OR: Benton Co., Corvallis, Crystal Lake Sports Complex, 44.5486°N 123.2485°W, 67m. 21.vi.2016. |
| *Amara farcta* | 293 | USA: OR: Lake Co., Christmas Valley Dunes, 43.35°N 120.3899°W, 1316m. 05.vi.2016. |
| *Anatrichis minuta* | 457 | USA: GA: Pulaski Co., forest nr Ocmulgee Public Fishing Area, 32.3815°N 83.4749°W, 86m. 23.ix.2016. |
| Anillini *gen. nov. sp. nov.* | 554 | USA: OR: Benton Co., Prarie Peak, 44.2837°N 123.5851°W. 15.xii.2017. |
| Anillini *gen. nov. sp. nov.* | 555 | USA: OR: Benton Co., Prarie Peak, 44.2837°N 123.5851°W. 15.xii.2017. |
| Anillini *gen. nov. sp. nov.* | 556 | USA: OR: Benton Co., Prarie Peak, 44.2837°N 123.5851°W. 15.xii.2017. |
| Anillini *gen. nov. sp. nov.* | 557 | USA: OR: Benton Co., Prarie Peak, 44.2837°N 123.5851°W. 15.xii.2017. |
| *Anisodactylus alternans* | 299 | USA: OR: Lake Co., Musser Reservoir, 43.5535°N 120.0074°W, 1230m. 05.vi.2016. |
| *Anisodactylus alternans* | 301 | USA: OR: Lake Co., Musser Reservoir, 43.5535°N 120.0074°W, 1230m. 05.vi.2016. |
| *Anisodactylus alternans* | 302 | USA: OR: Lake Co., Musser Reservoir, 43.5535°N 120.0074°W, 1230m. 05.vi.2016. |
| *Anisodactylus anthracinus* | 367 | USA: AZ: Pima Co., Canoa Ranch Rest Area, southbound I19, 31.7653°N 111.0351°W, 914m. 01-03.viii.2016. |
| *Anisodactylus anthracinus* | 368 | USA: AZ: Pima Co., Madera canyon visitor area, 31.7412°N 110.8871°W, 1346m. 02.viii.2016. |
| *Anisodactylus similis* | 300 | USA: OR: Lake Co., Musser Reservoir, 43.5535°N 120.0074°W, 1230m. 05.vi.2016. |
| *Anthia (Termophilum)* sp*.* | 621 | RSA: Limpopo: Ben Lavin Nature Reserve, 23.1247°S 29.9453°E, 867m. 31.i.2018. |
| *Apenes lucidula* | 399 | USA: AZ: Santa Cruz Co., Santa Cruz River, Santa Gertrudis Lane, 31.5621°N 111.0458°W. 10.viii.2016. |
| *Apotomus* sp. | 602 | RSA: KwaZulu-Natal: Highover Wildlife Sanctuary, 29.9136°S 30.1002°E, 547m. 26.i.2018. |
| *Apotomus* sp. | 627 | RSA: Limpopo: Makuya Nature Reserve, Mutale Falls Camp, 22.4268°S 31.0544°E, 301m. 02.ii.2018. |
| *Apotomus* sp. | 628 | RSA: Limpopo: Makuya Nature Reserve, Mutale Falls Camp, 22.4268°S 31.0544°E, 301m. 02.ii.2018. |
| *Apotomus* sp. | 650 | Mozambique: Sofala: Parque Nacional da Gorongosa, Chitengo camp, 18.9773°S 34.3515°E, 21m. 08-15.ii.2018. |
| *Ardistomis obliquata* | 439 | USA: GA: Pulaski Co., forest nr Ocmulgee Public Fishing Area, 32.3815°N 83.4749°W, 86m. 23.ix.2016. |
| *Ardistomis obliquata* | 448 | USA: GA: Pulaski Co., forest nr Ocmulgee Public Fishing Area, 32.3815°N 83.4749°W, 86m. 23.ix.2016. |
| *Ardistomis obliquata* | 522 | USA: Arkansas: Logan Co., Ozark Nat. Forest, Cove Lake Campground, 35.2256°N 93.6227°W, 318m. 29.viii.2017. |
| *Ardistomis schaumii* | 516 | USA: Arkansas: Logan Co., Ozark Nat. Forest, Cove Lake Campground, 35.2256°N 93.6227°W, 318m. 29.viii.2017. |
| *Ardistomis schaumii* | 517 | USA: Arkansas: Logan Co., Ozark Nat. Forest, Cove Lake Campground, 35.2256°N 93.6227°W, 318m. 29.viii.2017. |
| *Ardistomis schaumii* | 518 | USA: Arkansas: Logan Co., Ozark Nat. Forest, Cove Lake Campground, 35.2256°N 93.6227°W, 318m. 29.viii.2017. |
| *Ardistomis schaumii* | 520 | USA: Arkansas: Logan Co., Ozark Nat. Forest, Cove Lake Campground, 35.2256°N 93.6227°W, 318m. 29.viii.2017. |
| *Ardistomis schaumii* | 521 | USA: Arkansas: Logan Co., Ozark Nat. Forest, Cove Lake Campground, 35.2256°N 93.6227°W, 318m. 29.viii.2017. |
| *Aspidoglossa subangulata* | 304 | USA: AR: Newton Co., Shop Creek near Highway 327, 35.9511°N 93.2434°W. 16.vi.2016. |
| *Aspidoglossa subangulata* | 307 | USA: AR: Newton Co., Shop Creek near Highway 327, 35.9511°N 93.2434°W. 16.vi.2016. |
| *Badister ferrugineus* | 319 | USA: OR: Benton Co., Jackson-Frazier wetlands. 22.vii.2016. |
| *Bembidion incrematum* | 154 | USA: OR: Benton Co., Corvallis, Crystal Lake Sport Complex, 44.5497°N 123.2478°W. 29.iii.2016. |
| *Bembidion incrematum* | 160 | USA: OR: Benton Co., Corvallis, Crystal Lake Sport Complex, 44.5497°N 123.2478°W. 29.iii.2016. |
| *Bembidion incrematum* | 161 | USA: OR: Benton Co., Corvallis, Crystal Lake Sport Complex, 44.5497°N 123.2478°W. 29.iii.2016. |
| *Bembidion iridescens* | 155 | USA: OR: Benton Co., Corvallis, Crystal Lake Sport Complex, 44.5497°N 123.2478°W. 29.iii.2016. |
| *Bembidion iridescens* | 156 | USA: OR: Benton Co., Corvallis, Crystal Lake Sport Complex, 44.5497°N 123.2478°W. 29.iii.2016. |
| *Bembidion kuprianovi #2* | 163 | USA: OR: Benton Co., Corvallis, Crystal Lake Sport Complex, 44.5497°N 123.2478°W. 29.iii.2016. |
| *Bembidion obliquulum* | 215 | USA: OR: Benton Co., packed earth bank nr mouth of Marys river & Willamette river, 44.5561°N 123.2618°W, 61m. 30.iv.2016. |
| *Bembidion obliquulum* | 216 | USA: OR: Benton Co., packed earth bank nr mouth of Marys river & Willamette river, 44.5561°N 123.2618°W, 61m. 30.iv.2016. |
| *Bembidion sejunctum* | 256 | USA: OR: Lincoln Co., Ona Beach State Park, 44.5234°N 124.0736°W, 4m. 21.v.2016. |
| *Bembidion* sp. nr. *transversale* | 093 | USA: OR: Benton Co., Truax Island, 44.586°N 123.1858°W, 61m. 25.vii.2015. |
| *Bembidion* sp. nr. *transversale* | 094 | USA: OR: Benton Co., Truax Island, 44.586°N 123.1858°W, 61m. 25.vii.2015. |
| *Bembidion* sp. nr. *transversale* | 100 | USA: OR: Benton Co., Truax Island, 44.586°N 123.1858°W, 61m. 25.vii.2015. |
| *Bembidion* sp. nr. *transversale* | 235 | USA: OR: Benton Co., Irish Bend County Park, 44.3647°N 123.2206°W, 79m. 09.v.2016. |
| *Bembidion zephyrum* | 254 | USA: OR: Lincoln Co., Moolack Beach, 44.7093°N 124.0605°W, 4m. 21.v.2016. |
| *Bembidion zephyrum* | 255 | USA: OR: Lincoln Co., Moolack Beach, 44.7093°N 124.0605°W, 4m. 21.v.2016. |
| *Bembidion zephyrum* | 271 | USA: OR: Lincoln Co., Moolack Beach, 44.7093°N 124.0605°W, 4m. 21.v.2016. |
| *Blethisa oregonensis* | 311 | USA: WA: Lewis Co., Forst Borst Lake, Centralia, 46.7221°N 122.9782°W, 50m. 04.vi.2016 |
| *Blethisa oregonensis* | 320 | USA: OR: Benton Co., Jackson-Frazier wetlands. 22.vii.2016. |
| *Brachinus elongatulus* | 326 | USA: AZ: Pima Co., Madera Canyon, 31.7273°N 110.881°W, 1465m. 01.viii.2016. |
| *Brachinus elongatulus* | 355 | USA: AZ: Pima Co., Madera Canyon, 31.7273°N 110.881°W, 1465m. 01.viii.2016. |
| *Brachinus elongatulus* | 356 | USA: AZ: Pima Co., Madera Canyon, 31.7273°N 110.881°W, 1465m. 01.viii.2016. |
| *Brachinus elongatulus* | 357 | USA: AZ: Pima Co., Madera Canyon, 31.7273°N 110.881°W, 1465m. 01.viii.2016. |
| *Brachinus elongatulus* | 358 | USA: AZ: Pima Co., Madera Canyon, 31.7273°N 110.881°W, 1465m. 01.viii.2016. |
| *Brachinus elongatulus* | 369 | USA: AZ: Pima Co., Madera Canyon, 31.7273°N 110.881°W, 1465m. 01.viii.2016. |
| *Brachinus elongatulus* | 370 | USA: AZ: Pima Co., Madera Canyon, 31.7273°N 110.881°W, 1465m. 01.viii.2016. |
| *Brachinus ichabobopsis* | 456 | USA: FL: Liberty Co., Cotton Landing, 30.0512°N 85.0729°W, 33m. 13.ix.2016. |
| *Bradycellus* sp. 1 | 363 | USA: AZ: Pinal Co., Peppersauce Canyon nr campground, 32.5372°N 110.7195°W. 06.viii.2016. |
| *Bradycellus* sp. 2 | 308 | USA: AR: Newton Co., Shop Creek near Highway 327, 35.9511°N 93.2434°W. 16.vi.2016. |
| *Brasiella wickhami* | 343 | USA: AZ: Pima Co., Canoa Ranch Rest Area, southbound I19, 31.7653°N 111.0351°W, 914m. 01-03.viii.2016. |
| *Broscodera insignis* | 669 | USA: OR: Benton Co., Marys Peak, Parker Creek Falls, 44.4983°N 123.5688°W, 832m. 23.v.2018. |
| *Broscodera insignis* | 670 | USA: OR: Benton Co., Marys Peak, Parker Creek Falls, 44.4983°N 123.5688°W, 832m. 23.v.2018. |
| *Calathus peropacus* | 430 | USA: AZ: Cochise Co., Chiricahua Mtns, Rustler Park, 31.9068°N 109.2769°W, 2548m. 18.viii.2016. |
| *Calathus peropacus* | 431 | USA: AZ: Cochise Co., Chiricahua Mtns, Rustler Park, 31.9068°N 109.2769°W, 2548m. 18.viii.2016. |
| *Calathus peropacus* | 432 | USA: AZ: Cochise Co., Chiricahua Mtns, Rustler Park, 31.9068°N 109.2769°W, 2548m. 18.viii.2016. |
| *Calleida bella* | 117 | Mexico: Chiapas: roadside stop near Armando Zebadea pueblo, 16.9291°N 93.4737°W, 942m. 10.viii.2015. |
| *Calleida decora* | 376 | USA: AZ: Santa Cruz Co., Walker Canyon, 31.38°N 111.0671°W, 1207m. 08.viii.2016. |
| *Calleida jansoni* | 114 | Mexico: Veracruz: roadside beating tropical deciduous trees off of Highway 145D, 18.0165°N 94.2126°W, 119m. 09.viii.2015. |
| *Calleida jansoni* | 121 | Mexico: Chiapas: nr. Ocozocoautla, El Ocote Bio. Res., 16.8468°N 93.4946°W, 891m. 09.viii.2015. |
| *Calosoma peregrinator* | 335 | USA: AZ: Santa Cruz Co., Casa Blanca Canyon Rd, 31.6311°N 110.7566°W, 1355m. 03.viii.2016. |
| *Calosoma peregrinator* | 336 | USA: AZ: Santa Cruz Co., Casa Blanca Canyon Rd, 31.6311°N 110.7566°W, 1355m. 03.viii.2016. |
| *Carabus nemoralis* | 218 | USA: OR: Benton Co., Crystal Lake Sports Complex, 44.5486°N 123.2485°W, 67m. 30.iv.2016. |
| *Carabus taedatus* | 419 | USA: AZ: Cochise Co., Chiricahua Mtns, Rustler Park, 31.9068°N 109.2769°W, 2548m. 18.viii.2016. |
| *Catapiesis* sp. | 438 | Guatemala: Petén: Ixpanpajul, 200m. 16.8729°N 89.8148°W, 9-10.vii.2016. |
| *Cerapterus* sp. | 622 | RSA: Limpopo: Ben Lavin Nature Reserve, 23.1435°S 29.9497°E, 844m. 31.i.2018. |
| *Chlaenius cumatilis* | 383 | USA: AZ: Cochise Co., Charleston bridge, 31.6247°N 110.174°W, 1195m. 09.viii.2016. |
| *Chlaenius cumatilis* | 385 | USA: AZ: Cochise Co., Charleston bridge, 31.6247°N 110.174°W, 1195m. 09.viii.2016. |
| *Chlaenius glaucus* | 384 | USA: AZ: Cochise Co., Charleston bridge, 31.6247°N 110.174°W, 1195m. 09.viii.2016. |
| *Chlaenius harpalinus* | 287 | USA: OR: Lake Co., Musser Reservoir, 43.5535°N 120.0074°W, 1230m. 05.vi.2016. |
| *Chlaenius harpalinus* | 288 | USA: OR: Lake Co., Musser Reservoir, 43.5535°N 120.0074°W, 1230m. 05.vi.2016. |
| *Chlaenius leucoscelis* | 412 | USA: AZ: Cochise Co., San Pedro River nr Fairbank, AZ, 31.7189°N 110.1925°W, 1166m. 15.viii.2016. |
| *Chlaenius leucoscelis* | 413 | USA: AZ: Cochise Co., San Pedro River nr Fairbank, AZ, 31.7189°N 110.1925°W, 1166m. 15.viii.2016. |
| *Chlaenius prasinus* | 303 | USA: AR, Washington Co., Devil’s Den State Park, Lee Creek, 35.788398°N 94.246027°W. 17.vi.2016. |
| *Chlaenius ruficauda* | 381 | USA: AZ: Cochise Co., Charleston bridge, 31.6247°N 110.174°W, 1195m. 09.viii.2016. |
| *Chlaenius ruficauda* | 382 | USA: AZ: Cochise Co., Charleston bridge, 31.6247°N 110.174°W, 1195m. 09.viii.2016. |
| *Chlaenius sericeus* | 397 | USA: AZ: Santa Cruz Co., Santa Cruz River, Santa Gertrudis Lane, 31.5621°N 111.0458°W. 10.viii.2016. |
| *Chlaenius tricolor* | 411 | USA: AZ: Cochise Co., San Pedro River nr Fairbank, AZ, 31.7189°N 110.1925°W, 1166m. 15.viii.2016. |
| *Cicindela haemorrhagica* | 283 | USA: OR: Harney Co., Mickey Springs, 42.6782°N 118.3483°W, 1226m. 04.vi.2016. |
| *Cicindela haemorrhagica* | 284 | USA: OR: Harney Co., Mickey Springs, 42.6782°N 118.3483°W, 1226m. 04.vi.2016. |
| *Cicindela haemorrhagica* | 285 | USA: OR: Harney Co., Mickey Springs, 42.6782°N 118.3483°W, 1226m. 04.vi.2016. |
| *Clinidium* sp. nr. *guatemalenum* | 128 | Mexico: Oaxaca: La Cumbre, 16.4543°N 97.0033°W, 2196m. 15.viii.2015. |
| *Clinidium* sp. nr. *guatemalenum* | 130 | Mexico: Oaxaca: La Cumbre, 16.4543°N 97.0033°W, 2196m. 15.viii.2015. |
| *Clivina fossor* | 541 | USA: OR: Benton Co., Philomath, 44.5299°N 123.3716°W, 82m. 16.ix.2017. |
| *Clivina fossor* | 542 | USA: OR: Benton Co., Philomath, 44.5299°N 123.3716°W, 82m. 16.ix.2017. |
| *Clivina fossor* | 543 | USA: OR: Benton Co., Philomath, 44.5299°N 123.3716°W, 82m. 16.ix.2017. |
| *Colliuris pensylvanica* | 340 | USA: AZ: Santa Cruz Co., Casa Blanca Canyon Rd, 31.6311°N 110.7566°W, 1355m. 03.viii.2016. |
| *Colliuris pensylvanica* | 347 | USA: AZ: Pima Co., Canoa Ranch Rest Area, southbound I19, 31.7653°N 111.0351°W, 914m. 01-03.viii.2016. |
| *Colliuris pensylvanica* | 348 | USA: AZ: Pima Co., Canoa Ranch Rest Area, southbound I19, 31.7653°N 111.0351°W, 914m. 01-03.viii.2016. |
| *Cychrus tuberculatus* | 671 | USA: OR: Benton Co., Marys Peak, 44.5089°N 123.5565°W, 1088m. 23.v.2018. |
| *Cycloloba* sp. | 585 | RSA: KwaZulu-Natal: Lotheni Nature Reserve, 29.4365°S 29.5121°E, 1578m, 20-21.i.2018. |
| *Cyclotrachelus dejeanellus* | 452 | USA: FL: Marion Co., Ocala National Forest, Juniper Springs Recreation Area, 29.1824°N 81.708°W, 25m. 15.ix.2016. |
| *Cyclotrachelus dejeanellus* | 453 | USA: FL: Marion Co., Ocala National Forest, Juniper Springs Recreation Area, 29.1824°N 81.708°W, 25m. 15.ix.2016. |
| *Cyclotrachelus dejeanellus* | 454 | USA: FL: Marion Co., Ocala National Forest, Juniper Springs Recreation Area, 29.1824°N 81.708°W, 25m. 15.ix.2016. |
| *Cymindis basipunctata-group sp.* | 119 | Mexico: Chiapas: pullout on highway 85 nr. El Chapopote, 16.881°N 93.4591°W, 1072m. 10.viii.2015. |
| *Cymindis punctifera* | 365 | USA: AZ: Santa Cruz Co., Casa Blanca Canyon Rd, 31.6311°N 110.7566°W, 1355m. 03.viii.2016. |
| *Cymindis punctigera* | 339 | USA: AZ: Pima Co., Madera canyon visitor area, 31.7412°N 110.8871°W, 1346m. 02.viii.2016. |
| *Cyrtomoscelis cf. dwesana* | 624 | RSA: Eastern Cape: Mkhambathi Nature Reserve, forest behind large group cabin, 31.3172°S 29.9673°E, 30m. 24-25.i.2018. |
| *Cyrtomoscelis cf. dwesana* | 625 | RSA: Eastern Cape: Mkhambathi Nature Reserve, forest behind large group cabin, 31.3172°S 29.9673°E, 30m. 24-25.i.2018. |
| *Cyrtomoscelis cf. dwesana* | 626 | RSA: Eastern Cape: Mkhambathi Nature Reserve, forest behind large group cabin, 31.3172°S 29.9673°E, 30m. 24-25.i.2018. |
| *Dicaelus suffusus* | 422 | USA: AZ: Cochise Co., Chiricahua Mtns, Rustler Park, 31.9068°N 109.2769°W, 2548m. 18.viii.2016. |
| *Dicaelus suffusus* | 423 | USA: AZ: Cochise Co., Chiricahua Mtns, Rustler Park, 31.9068°N 109.2769°W, 2548m. 18.viii.2016. |
| *Dicaelus suffusus* | 424 | USA: AZ: Cochise Co., Chiricahua Mtns, Rustler Park, 31.9068°N 109.2769°W, 2548m. 18.viii.2016. |
| *Diplochaetus planatus* | 316 | USA: OR: Lake Co., Lake Abert, 42.6414°N 120.1837°W, 1298m. 10.vii.2016. |
| *Diplochaetus planatus* | 499 | USA: NM: Torrance Co., Laguna del Perro, 34.6003°N 105.9252°W, 1861m. 24.viii.2017. |
| *Diplochaetus planatus* | 500 | USA: NM: Torrance Co., Laguna del Perro, 34.6003°N 105.9252°W, 1861m. 24.viii.2017. |
| *Diplochaetus planatus* | 501 | USA: NM: Torrance Co., Laguna del Perro, 34.6003°N 105.9252°W, 1861m. 24.viii.2017. |
| *Diplochaetus planatus* | 502 | USA: NM: Torrance Co., Laguna del Perro, 34.6003°N 105.9252°W, 1861m. 24.viii.2017. |
| *Diplocheila nupera* | 437 | USA: FL: Monroe Co., Marathon, Florida Keys Wildlife and Environmental Area, 24.7302°N 81.0551°W, 7m. 18.ix.2016. |
| *Diplous filicornis* | 481 | USA: OR: Benton Co., Irish Bend County Park, 44.3627°N 123.2199°W, 80m. 29.iv.2017. |
| *Discoderus* sp. | 361 | USA: AZ: Pinal Co., Peppersauce Canyon nr campground, 32.5372°N 110.7195°W, 06.viii.2016. |
| *Disphaericus* sp. | 641 | Mozambique: Sofala: Parque Nacional da Gorongosa, Chitengo camp, 18.9773°S 34.3515°E, 21m. 08-15.ii.2018. |
| *Disphaericus* sp*.* | 642 | Mozambique: Sofala: Parque Nacional da Gorongosa, Chitengo camp, 18.9773°S 34.3515°E, 21m. 08-15.ii.2018. |
| *Drypta* sp. | 577 | RSA: KwaZulu-Natal: Royal Natal National Park, 28.687°S 28.9539°E, 1404m. 18.i.2018. |
| *Drypta* sp. | 579 | RSA: KwaZulu-Natal: Royal Natal National Park, 28.7117°S 28.9371°E, 1540m. 18.i.2018. |
| *Drypta* sp. | 584 | RSA: KwaZulu-Natal: Hlalanathi Drakensberg Resort, 28.6558°S 29.0345°E, 1264m. 19.i.2018. |
| *Dyschirius dejeanii* | 298 | USA: OR: Harney Co., Ten Cent Lake, 42.9893°N 118.284°W, 1230m. 05.vi.2016. |
| *Dyschirius globosus* | 482 | Germany: Mecklenburg: Rostocker Heide, "Speckingsbruch" near Torfbrucke, 54.2334°N 12.2192°E, 15m. 26.v.2017. |
| *Dyschirius globosus* | 483 | Germany: Mecklenburg: Rostocker Heide, "Speckingsbruch" near Torfbrucke, 54.2334°N 12.2192°E, 15m. 26.v.2017. |
| *Dyschirius globosus* | 484 | Germany: Mecklenburg: Rostocker Heide, "Speckingsbruch" near Torfbrucke, 54.2334°N 12.2192°E, 15m. 26.v.2017. |
| *Dyschirius globosus* | 485 | Germany: Mecklenburg: Rostocker Heide, "Speckingsbruch" near Torfbrucke, 54.2334°N 12.2192°E, 15m. 26.v.2017. |
| *Dyschirius globosus* | 490 | Germany: Mecklenburg: Rostocker Heide, "Speckingsbruch" near Torfbrucke, 54.2334°N 12.2192°E, 15m. 26.v.2017. |
| *Dyschirius globosus* | 491 | Germany: Mecklenburg: Rostocker Heide, "Speckingsbruch" near Torfbrucke, 54.2334°N 12.2192°E, 15m. 26.v.2017. |
| *Dyschirius haemorrhoidalis* | 529 | USA: Arkansas: Madison Co., Kings River public access, 36.1432°N 93.5941°W, 373m. 01.ix.2017. |
| *Dyschirius pacificus* | 274 | USA: OR: Lincoln Co., Moolack Beach, 44.7093°N 124.0605°W, 4m. 21.v.2016. |
| *Dyschirius pacificus* | 275 | USA: OR: Lincoln Co., Moolack Beach, 44.7093°N 124.0605°W, 4m. 21.v.2016. |
| *Dyschirius thoracicus* | 486 | Germany: Mecklenburg: Rostock-Markgrafenheide, 54.1748°N 12.1424°E, 0m. 23.v.2017. |
| *Dyschirius thoracicus* | 487 | Germany: Mecklenburg: Rostock-Markgrafenheide, 54.1748°N 12.1424°E, 0m. 23.v.2017. |
| *Dyschirius thoracicus* | 488 | Germany: Mecklenburg: Rostock-Markgrafenheide, 54.1748°N 12.1424°E, 0m. 23.v.2017. |
| *Dyschirius thoracicus* | 489 | Germany: Mecklenburg: Rostock-Markgrafenheide, 54.1748°N 12.1424°E, 0m. 23.v.2017. |
| *Dyschirius tridentatus* | 273 | USA: OR: Benton Co., Irish Bend County Park, 44.3647°N 123.2206°W, 79m. 15.v.2016. |
| *Ega sallei* | 440 | USA: GA: public boat ramp in Hawkinsville, GA, 32.2837°N 83.4625°W, 70m. 23.ix.2016. |
| *Elaphrus purpurans* | 188 | USA: OR: Benton Co., Corvallis, Crystal Lake Sports Complex, 44.5497°N 123.2453°W. 09.iv.2016. |
| *Elaphrus purpurans* | 204 | USA: OR: Benton Co., Corvallis, Crystal Lake Sport Complex, 44.5497°N 123.2478°W. 29.iii.2016. |
| *Elaphrus purpurans* | 157 | USA: OR: Benton Co., Corvallis, Crystal Lake Sport Complex, 44.5497°N 123.2478°W. 29.iii.2016. |
| *Elaphrus purpurans* | 158 | USA: OR: Benton Co., Corvallis, Crystal Lake Sport Complex, 44.5497°N 123.2478°W. 29.iii.2016. |
| *Elaphrus purpurans* | 223 | USA: OR: Benton Co., Corvallis, Crystal Lake Sports Complex, 44.5497°N 123.2453°W. 09.iv.2016. |
| *Eucamaragnathus oxygonus* | 588 | RSA: Eastern Cape: Mtentu River Lodge, 31.2448°S 30.0462°E, 33m. 23.i.2018. |
| *Eucamaragnathus oxygonus* | 589 | RSA: Eastern Cape: Mtentu River Lodge, 31.2448°S 30.0462°E, 33m. 23.i.2018. |
| *Eucamaragnathus oxygonus* | 590 | RSA: Eastern Cape: Mtentu River Lodge, 31.2448°S 30.0462°E, 33m. 23.i.2018. |
| *Euryderus grossus* | 416 | USA: AZ: Cochise Co., S Blue Sky Rd S of Wilcox, AZ, 32.2335°N 109.7774°W, 1269m. 17.viii.2016. |
| *Euryderus grossus* | 417 | USA: AZ: Cochise Co., S Blue Sky Rd S of Wilcox, AZ, 32.2335°N 109.7774°W, 1269m. 17.viii.2016. |
| *Galerita atripes* | 395 | USA: AZ: Santa Cruz Co., Harshaw Creek, 31.4387°N 110.7247°W, 1600m. 10.viii.2016. |
| *Galerita bicolor* | 455 | USA: FL: Liberty Co., Cotton Landing, 30.0512°N 85.0729°W, 33m. 13.ix.2016. |
| *Galerita forreri* | 324 | USA: AZ: Pima Co., Madera canyon visitor area, 31.7412°N 110.8871°W, 1346m. 02.viii.2016. |
| *Galerita lecontei* | 380 | USA: AZ: Cochise Co., Charleston bridge, 31.6247°N 110.174°W, 1195m. 09.viii.2016. |
| *Galerita lecontei* | 408 | USA: AZ: Cochise Co., San Pedro River nr Fairbank, AZ, 31.7189°N 110.1925°W, 1166m. 15.viii.2016. |
| *Gehringia olympica* | 679 | USA: MT: Broadwater Co., Confederate Gulch, 46.5641°N 111.465°W, 1332m. 22.viii.2018. |
| *Gehringia olympica* | 680 | USA: MT: Broadwater Co., Confederate Gulch, 46.5641°N 111.465°W, 1332m. 22.viii.2018. |
| *Goniotropis parca* | 377 | USA: AZ: Santa Cruz Co., Walker Canyon, 31.38°N 111.0671°W, 1207m. 08.viii.2016. |
| *Goniotropis parca* | 378 | USA: AZ: Santa Cruz Co., Walker Canyon, 31.38°N 111.0671°W, 1207m. 08.viii.2016. |
| *Goniotropis parca* | 379 | USA: AZ: Santa Cruz Co., Walker Canyon, 31.38°N 111.0671°W, 1207m. 08.viii.2016. |
| *Graphipterus* sp. | 658 | Mozambique: Sofala: Parque Nacional da Gorongosa, 18.9766°S 34.3509°E, 39m. 15.ii.2018. |
| *Haplotrachelus atropsis* | 592 | RSA: Eastern Cape: Mtentu River Lodge, 31.2448°S 30.0462°E, 33m. 23.i.2018. |
| *Haplotrachelus cf. latesulcatus* | 598 | RSA: Eastern Cape: Mkhambathi Nature Reserve, 31.2886°S 30.0103°E, 17m. 25.i.2018. |
| *Haplotrachelus cf. latesulcatus* | 599 | RSA: Eastern Cape: Mkhambathi Nature Reserve, 31.2886°S 30.0103°E, 17m, 25.i.2018. RAG18012504. |
| *Haplotrachelus* sp. | 591 | RSA: Eastern Cape: Mtentu River Lodge, 31.2448°S 30.0462°E, 33m. 23.i.2018. |
| *Haplotrachelus* sp*.* | 593 | RSA: Eastern Cape: Mtentu River Lodge, 31.2448°S 30.0462°E, 33m. 23.i.2018. |
| *Harpalus affinis* | 166 | USA: OR: Benton Co., Corvallis, Crystal Lake Sports Complex, 44.5497°N 123.2453°W, 09.iv.2016. |
| *Harpalus affinis* | 167 | USA: OR: Benton Co., Corvallis, Crystal Lake Sports Complex, 44.5497°N 123.2453°W, 09.iv.2016. |
| *Harpalus affinis* | 168 | USA: OR: Benton Co., Corvallis, Crystal Lake Sports Complex, 44.5497°N 123.2453°W, 09.iv.2016. |
| *Harpalus affinis* | 169 | USA: OR: Benton Co., Corvallis, Crystal Lake Sports Complex, 44.5497°N 123.2453°W, 09.iv.2016. |
| *Helluomorphoides papago* | 325 | USA: AZ: Pima Co., Madera canyon visitor area, 31.7412°N 110.8871°W, 1346m. 02.viii.2016. |
| *Helluomorphoides papago* | 374 | USA: AZ: Santa Cruz Co., Walker Canyon, 31.38°N 111.0671°W, 1207m. 08.viii.2016. |
| *Hybothecus flohri* | 409 | USA: AZ: Cochise Co., San Pedro River nr Fairbank, AZ, 31.7189°N 110.1925°W, 1166m. 15.viii.2016. |
| *Lachnophorus elegantulus* | 405 | USA: AZ: Santa Cruz Co., Pena Blanca Lake parking area, 31.3986°N 111.0891°W, 1169m. 13.viii.2016. |
| *Lachnophorus elegantulus* | 406 | USA: AZ: Santa Cruz Co., Pena Blanca Lake parking area, 31.3986°N 111.0891°W, 1169m. 13.viii.2016. |
| *Lachnophorus* sp. | 134 | Mexico: Oaxaca: stop off of Highway 125 nr. La Esperanza, 16.4893°N 98.1085°W, 87m. 17.viii.2015. |
| *Lebia deceptrix* | 400 | USA: AZ: Santa Cruz Co., Walker Canyon, 31.38°N 111.0671°W, 1207m. 08.viii.2016. |
| *Lebia deceptrix* | 401 | USA: AZ: Santa Cruz Co., Walker Canyon, 31.38°N 111.0671°W, 1207m. 08.viii.2016. |
| *Lebia subgrandis* | 338 | USA: AZ: Santa Cruz Co., Casa Blanca Canyon Rd, 31.6311°N 110.7566°W, 1355m. 03.viii.2016. |
| *Lebia viridis* | 342 | USA: AZ: Santa Cruz Co., Pena Blanca Lake parking area, 31.3986°N 111.0891°W, 1169m. 13.viii.2016. |
| *Lebia viridis* | 364 | USA: AZ: Pinal Co., Peppersauce Canyon nr campground, 32.5372°N 110.7195°W. 06.viii.2016. |
| *Leptotrachelus* sp. | 450 | Guatemala: Alta Verapaz: Las cuevas, nr. intersection rt 5/9, 15.8683°N 90.0968°W, 180m. 7.viii.2016. |
| *Lionepha sp. nov.* | 212 | USA: OR: Benton Co., Marys Peak, rock seeps nr Alder falls, 44.4744°N 123.5286°W, 679m. 24.iv.2016. |
| *Lionepha sp. nov.* | 213 | USA: OR: Benton Co., Marys Peak, rock seeps nr Alder falls, 44.4744°N 123.5286°W, 679m. 24.iv.2016. |
| *Lionepha sp. nov.* | 214 | USA: OR: Benton Co., Marys Peak, rock seeps nr Alder falls, 44.4744°N 123.5286°W, 679m. 24.iv.2016. |
| *Loricera decempunctata* | 096 | USA: OR: Benton Co., Truax Island, 44.586°N 123.1858°W, 61m. 25.vii.2015. |
| *Loricera decempunctata* | 185 | USA: OR: Benton Co., Corvallis, Crystal Lake Sports Complex, 44.5497°N 123.2453°W, 09.iv.2016. |
| *Loricera decempunctata* | 186 | USA: OR: Benton Co., Corvallis, Crystal Lake Sports Complex, 44.5497°N 123.2453°W, 09.iv.2016. |
| *Loricera foveata* | 191 | USA: OR: Benton Co., Corvallis, Crystal Lake Sports Complex, 44.5497°N 123.2453°W, 09.iv.2016. |
| *Loricera foveata* | 145 | USA: OR: Benton Co., Corvallis, Crystal Lake Sport Complex, 44.5497°N 123.2478°W. 29.iii.2016. |
| *Macrocheilus* sp. | 623 | RSA: KwaZulu-Natal: Ithala Game Reserve, Ntshondwe Lodge, 27.5435°S 31.2829°E, 430m. 30.i.2018. |
| *Mastax* sp*.* | 594 | RSA: Eastern Cape: river crossing, 31.1302°S 29.7559°E. 23.i.2018. |
| *Mastax* sp. | 655 | Mozambique: Sofala: Parque Nacional da Gorongosa, 18.9987°S 34.3578°E, 41m. 13.ii.2018. |
| *Mastax* sp. | 656 | Mozambique: Sofala: Parque Nacional da Gorongosa, 18.9987°S 34.3578°E, 41m. 13.ii.2018. |
| *Mastax* sp. | 657 | Mozambique: Sofala: Parque Nacional da Gorongosa, 18.9987°S 34.3578°E, 41m. 13.ii.2018. |
| *Metrius contractus* | 252 | USA: OR: Benton Co., Chip Ross Park, 44.6065°N 123.2825°W, 200m. 06.v.2016. |
| *Metrius contractus* | 253 | USA: OR: Benton Co., Chip Ross Park, 44.6065°N 123.2825°W, 200m. 06.v.2016. |
| *Metrius contractus* | 266 | USA: OR: Benton Co., Chip Ross Park, 44.6065°N 123.2825°W, 200m. 06.v.2016. |
| *Mioptachys flavicauda* | 442 | USA: GA: Pulaski Co., forest nr Ocmulgee Public Fishing Area, 32.3815°N 83.4749°W, 86m. 23.ix.2016. |
| *Mioptachys flavicauda* | 443 | USA: GA: Pulaski Co., forest nr Ocmulgee Public Fishing Area, 32.3815°N 83.4749°W, 86m. 23.ix.2016. |
| *Mioptachys flavicauda* | 444 | USA: GA: Pulaski Co., forest nr Ocmulgee Public Fishing Area, 32.3815°N 83.4749°W, 86m. 23.ix.2016. |
| *Mioptachys flavicauda* | 445 | USA: GA: Pulaski Co., forest nr Ocmulgee Public Fishing Area, 32.3815°N 83.4749°W, 86m. 23.ix.2016. |
| *Morion* sp. | 115 | Mexico: Chiapas: roadside stop near Armando Zebadea pueblo, 16.9291°N 93.4737°W, 942m. 10.viii.2015. |
| *Nebria brevicollis* | 441 | USA: OR: Benton Co., Crystal Lake Sports Complex, 44.5506°N 123.2498°W, 61m. 30.ix.2016. |
| *Notiophilus sylvaticus* | 267 | USA: OR: Benton Co., Chip Ross Park, 44.6065°N 123.2825°W, 200m, 27.v.2016. |
| *Notiophilus sylvaticus* | 268 | USA: OR: Benton Co., Chip Ross Park, 44.6065°N 123.2825°W, 200m, 27.v.2016. |
| *Notiophilus sylvaticus* | 269 | USA: OR: Benton Co., Chip Ross Park, 44.6065°N 123.2825°W, 200m, 27.v.2016. |
| *Omoglymmius hamatus* | 249 | USA: OR: Jefferson Co., pine forest nr Lower Bridge campground, 44.5597°N 121.6185°W, 851m. 14.v.2016. |
| *Omoglymmius hamatus* | 250 | USA: OR: Jefferson Co., pine forest nr Lower Bridge campground, 44.5597°N 121.6185°W, 851m. 14.v.2016. |
| *Omoglymmius hamatus* | 251 | USA: OR: Jefferson Co., pine forest nr Lower Bridge campground, 44.5597°N 121.6185°W, 851m. 14.v.2016. |
| *Omophron americanum* | 310 | USA: AR, Washington Co., Devil’s Den State Park, Lee Creek, 35.7844°N 94.2437°W. 17.vi.2016. |
| *Omophron ovale* | 236 | USA: OR: Benton Co., Irish Bend County Park, 44.3647°N 123.2206°W, 79m. 09.v.2016. |
| *Omophron ovale* | 237 | USA: OR: Benton Co., Irish Bend County Park, 44.3647°N 123.2206°W, 79m. 09.v.2016. |
| *Omophron ovale* | 247 | USA: OR: Benton Co., Irish Bend County Park, 44.3647°N 123.2206°W, 79m. 10.v.2016. |
| *Omophron ovale* | 248 | USA: OR: Lincoln Co., Moolack Beach, 44.7093°N 124.0605°W, 4m. 21.v.2016. |
| *Omus audouini* | 193 | USA: OR: Lane Co., S of Veneta, 6 mi S of Crow, private property off of route 36 Territorial Hwy. 16-17.iv.2016. |
| *Omus audouini* | 194 | USA: OR: Lane Co., S of Veneta, 6 mi S of Crow, private property off of route 36 Territorial Hwy. 16-17.iv.2016. |
| *Omus audouini* | 195 | USA: OR: Lane Co., S of Veneta, 6 mi S of Crow, private property off of route 36 Territorial Hwy. 16-17.iv.2016. |
| *Omus audouini* | 196 | USA: OR: Lane Co., S of Veneta, 6 mi S of Crow, private property off of route 36 Territorial Hwy. 16-17.iv.2016. |
| *Omus audouini* | 198 | USA: OR: Lane Co., S of Veneta, 6 mi S of Crow, private property off of route 36 Territorial Hwy. 16-17.iv.2016. |
| *Omus audouini* | 200 | USA: OR: Lane Co., S of Veneta, 6 mi S of Crow, private property off of route 36 Territorial Hwy. 16-17.iv.2016. |
| *Omus audouini* | 272 | USA: OR: Lane Co., S of Veneta, 6 mi S of Crow, private property off of route 36 Territorial Hwy. 16-17.iv.2016. |
| *Omus dejeanii* | 224 | USA: OR: Benton Co., McDonald-Dunn forest, Lewisburg saddle, 44.636°N 123.2972°W, 314m. 06.v.2016. |
| *Omus dejeanii* | 225 | USA: OR: Benton Co., McDonald-Dunn forest, Lewisburg saddle, 44.636°N 123.2972°W, 314m. 06.v.2016. |
| *Omus dejeanii* | 226 | USA: OR: Benton Co., McDonald-Dunn forest, Lewisburg saddle, 44.636°N 123.2972°W, 314m. 06.v.2016. |
| *Oodes fluvialis* | 462 | USA: Florida: Glades Co., Fisheating Creek WMA, 26.9306°N 81.2877°W, 8.4m. 16.ix.2016. |
| *Oodes fluvialis* | 463 | USA: FL: Liberty Co., Cotton Landing, 30.0512°N 85.0729°W, 33m. 13.ix.2016. |
| *Oodes fluvialis* | 495 | USA: Arkansas: Logan Co., Ozark Nat. Forest, Cove Lake Campground, 35.2256°N 93.6227°W, 318m. 29.viii.2017. |
| *Oodes fluvialis* | 504 | USA: Arkansas: Montgomery Co., Private campground nr Ouachita River, 34.6333°N 93.6481°W, 204m. 27.viii.2017. |
| *Oodes fluvialis* | 505 | USA: Arkansas: Montgomery Co., Private campground nr Ouachita River, 34.6333°N 93.6481°W, 204m. 27.viii.2017. |
| *Oodes fluvialis* | 506 | USA: Arkansas: Montgomery Co., Private campground nr Ouachita River, 34.6333°N 93.6481°W, 204m. 27.viii.2017. |
| *Oodes fluvialis* | 507 | USA: Arkansas: Montgomery Co., Private campground nr Ouachita River, 34.6333°N 93.6481°W, 204m. 27.viii.2017. |
| *Oodes fluvialis* | 508 | USA: Arkansas: Montgomery Co., Private campground nr Ouachita River, 34.6333°N 93.6481°W, 204m. 27.viii.2017. |
| *Opisthius richardsoni* | 245 | USA: OR: Benton Co., Irish Bend County Park, 44.3647°N 123.2206°W, 79m. 15.v.2016. |
| *Opisthius richardsoni* | 246 | USA: OR: Benton Co., Irish Bend County Park, 44.3647°N 123.2206°W, 79m. 15.v.2016. |
| *Ozaena* sp. | 108 | Mexico: Veracruz: Est. Biol. Los Tuxtlas, 18.5855°N 95.0752°W, 125m. 06-09.viii.2015. |
| *Pachydesus* sp*.* | 564 | RSA: KwaZulu-Natal: Royal Natal National Park, 28.6906°S 28.949°E, 1431m. 17.i.2018. |
| *Pachydesus* sp. | 565 | RSA: KwaZulu-Natal: Royal Natal National Park, 28.6906°S 28.949°E, 1431m. 17.i.2018. |
| *Pachydesus* sp. | 586 | RSA: KwaZulu-Natal: pullout beside creek on Lower Lotheni Rd, 29.5224°S 29.646°E, 1555m. 21.i.2018. |
| *Pachydesus* sp. | 587 | RSA: KwaZulu-Natal: pullout beside creek on Lower Lotheni Rd, 29.5224°S 29.646°E, 1555m. 21.i.2018. |
| *Pachyteles* sp. | 109 | Mexico: Veracruz: Est. Biol. Los Tuxtlas, 18.5855°N 95.0752°W, 125m. 06-09.viii.2015. |
| *Panagaeus sallei* | 327 | USA: AZ: Pima Co., Canoa Ranch Rest Area, southbound I19, 31.7653°N 111.0351°W, 914m. 01-03.viii.2016. |
| *Panagaeus sallei* | 328 | USA: AZ: Pima Co., Canoa Ranch Rest Area, southbound I19, 31.7653°N 111.0351°W, 914m. 01-03.viii.2016. |
| *Panagaeus sallei* | 352 | USA: AZ: Pima Co., Canoa Ranch Rest Area, southbound I19, 31.7653°N 111.0351°W, 914m. 01-03.viii.2016. |
| *Paraclivina bipustulata* | 387 | USA: AZ: Cochise Co., San Pedro River, 31.5501°N 110.1375°W, 1237m. 09.viii.2016. |
| *Paraclivina bipustulata* | 388 | USA: AZ: Cochise Co., San Pedro River, 31.5501°N 110.1375°W, 1237m. 09.viii.2016. |
| *Paratachys* sp. 1 | 640 | RSA: Limpopo: Mashovhela Bush Lodge, 22.944°S 29.8907°E, 512m. 04.ii.2018. |
| *Paratachys* sp. 2 | 648 | Mozambique: Sofala: Parque Nacional da Gorongosa, 18.9923°S 34.3505°E, 32m. 10.ii.2018. |
| *Paratachys* sp. 2 | 649 | Mozambique: Sofala: Parque Nacional da Gorongosa, 18.9923°S 34.3505°E, 32m. 10.ii.2018. |
| *Pasimachus californicus* | 359 | USA: AZ: Pima Co., Madera canyon visitor area, 31.7412°N 110.8871°W, 1346m. 02.viii.2016. |
| *Pasimachus californicus* | 360 | USA: AZ: Pima Co., Madera canyon visitor area, 31.7412°N 110.8871°W, 1346m. 02.viii.2016. |
| *Pasimachus californicus* | 329 | USA: AZ: Pima Co., Madera canyon visitor area, 31.7412°N 110.8871°W, 1346m. 02.viii.2016. |
| *Patrobus longicornis* | 494 | USA: Arkansas: Logan Co., Ozark Nat. Forest, Cove Lake Campground, 35.2256°N 93.6227°W, 318m. 29.viii.2017. |
| *Patrobus longicornis* | 496 | USA: Missouri: Boone Co., Columbia, MO, Grindstone Nature Area, 38.928°N 92.3217°W. 03.ix.2017. |
| *Patrobus longicornis* | 497 | USA: Missouri: Boone Co., Columbia, MO, Grindstone Nature Area, 38.928°N 92.3217°W. 03.ix.2017. |
| *Patrobus longicornis* | 498 | USA: Missouri: Boone Co., Columbia, MO, Grindstone Nature Area, 38.928°N 92.3217°W. 03.ix.2017. |
| *Paussus (Bathypaussus)* sp. | 580 | RSA: KwaZulu-Natal: Hlalanathi Drakensberg Resort, 28.6597°S 29.0322°E, 1273m. 19.i.2018. |
| *Paussus cucullatus* | 603 | RSA: KwaZulu-Natal: Highover Wildlife Sanctuary, 29.9135°S 30.0996°E, 546m. 27.i.2018. |
| *Paussus cucullatus* | 612 | RSA: KwaZulu-Natal: Hluhluwe-iMfolozi Park, Sontuli picnic area, 28.2297°S 31.845°E, 117m. 29.i.2018. |
| *Pentagonica* sp. | 353 | USA: AZ: Santa Cruz Co., Casa Blanca Canyon Rd, 31.6311°N 110.7566°W, 1355m. 03.viii.2016. |
| *Pentagonica* sp. | 366 | USA: AZ: Pima Co., Madera canyon visitor area, 31.7412°N 110.8871°W, 1346m. 02.viii.2016. |
| *Perigona nigriceps* | 449 | USA: FL: Monroe Co., Long Key State Park, 24.8105°N 80.829°W, 17m. 18.ix.2016. |
| *Perileptus* sp. | 595 | RSA: Eastern Cape: river crossing, 31.1302°S 29.7559°E, 283m. 23.i.2018. |
| *Perileptus* sp. | 596 | RSA: Eastern Cape: river crossing, 31.1302°S 29.7559°E, 283m. 23.i.2018. |
| *Pheropsophus* sp. 1 | 566 | RSA: KwaZulu-Natal: Royal Natal National Park, 28.687°S 28.9539°E, 1404m. 18.i.2018. |
| *Pheropsophus* sp. 1 | 567 | RSA: KwaZulu-Natal: Royal Natal National Park, 28.687°S 28.9539°E, 1404m. 18.i.2018. |
| *Pheropsophus* sp. 1 | 568 | RSA: KwaZulu-Natal: Royal Natal National Park, 28.687°S 28.9539°E, 1404m. 18.i.2018. |
| *Pheropsophus* sp. 1 | 569 | RSA: KwaZulu-Natal: Royal Natal National Park, 28.687°S 28.9539°E, 1404m. 18.i.2018. |
| *Pheropsophus* sp. 1 | 570 | RSA: KwaZulu-Natal: Royal Natal National Park, 28.687°S 28.9539°E, 1404m. 18.i.2018. |
| *Pheropsophus* sp. 2 | 597 | RSA: Eastern Cape: Mkhambathi Nature Reserve, 31.2886°S 30.0103°E, 17m. 25.i.2018. |
| *Phloeoxena nigricollis* | 110 | Mexico: Veracruz: Ruiz Cortines, San Andres Tuxtla, 18.5273°N 95.1372°W, 1072m. 08.viii.2015. |
| *Poecilus laetulus* | 289 | USA: OR: Harney Co., Mann Lake, 42.777°N 118.4432°W, 1230m. 05.vi.2016. |
| *Poecilus laetulus* | 291 | USA: OR: Harney Co., Mann Lake, 42.777°N 118.4432°W, 1230m. 05.vi.2016. |
| *Poecilus scitulus* | 290 | USA: OR: Harney Co., Mann Lake, 42.777°N 118.4432°W, 1230m. 05.vi.2016. |
| *Polpochila erro* | 404 | USA: AZ: Pima Co., Happy Valley, oak forest, 32.1559°N 110.4748°W, 1262m. 14.viii.2016. |
| *Polpochila erro* | 410 | USA: AZ: Cochise Co., San Pedro River nr Fairbank, AZ, 31.7189°N 110.1925°W, 1166m. 15.viii.2016. |
| *Promecognathus laevissimus* | 227 | USA: OR: Benton Co., McDonald-Dunn forest, Lewisburg saddle, 44.636°N 123.2972°W, 314m. 06.v.2016. |
| *Promecognathus laevissimus* | 228 | USA: OR: Benton Co., McDonald-Dunn forest, Lewisburg saddle, 44.636°N 123.2972°W, 314m. 06.v.2016. |
| *Pseudaptinus horni* | 373 | USA: AZ: Santa Cruz Co., Walker Canyon, 31.38°N 111.0671°W, 1207m. 08.viii.2016. |
| *Pseudaptinus simplex* | 414 | USA: AZ: Pinal Co., Winkelman, AZ, 32.9852°N 110.7676°W, 587m. 11.viii.2016. |
| *Pseudaptinus tenuicollis* | 372 | USA: AZ: Santa Cruz Co., Walker Canyon, 31.38°N 111.0671°W, 1207m. 08.viii.2016. |
| *Pseudomorpha* sp. | 371 | USA: AZ: Santa Cruz Co., Walker Canyon, 31.38°N 111.0671°W, 1207m. 08.viii.2016. |
| *Psydrus piceus* | 415 | USA: AZ: Cochise Co., Chiricahua Mtns, Rustler Park, 31.9068°N 109.2769°W, 2548m. 18.viii.2016. |
| *Pterostichus infernalis* | 207 | USA: OR: Benton Co., Marys Peak, trail nr boundary btwn forest and meadow, 44.5091°N 123.5526°W, 1141m. 19.iv.2016. |
| *Pterostichus infernalis* | 208 | USA: OR: Benton Co., Marys Peak, trail nr boundary btwn forest and meadow, 44.5091°N 123.5526°W, 1141m. 19.iv.2016. |
| *Pterostichus infernalis* | 209 | USA: OR: Benton Co., Marys Peak, trail nr boundary btwn forest and meadow, 44.5091°N 123.5526°W, 1141m. 19.iv.2016. |
| *Pterostichus infernalis* | 210 | USA: OR: Benton Co., Marys Peak, trail nr boundary btwn forest and meadow, 44.5091°N 123.5526°W, 1141m. 19.iv.2016. |
| *Pterostichus infernalis* | 222 | USA: OR: Benton Co., Marys Peak, trail nr boundary btwn forest and meadow, 44.5091°N 123.5526°W, 1141m. 19.iv.2016. |
| *Pterostichus lama* | 229 | USA: OR: Benton Co., McDonald-Dunn forest, Lewisburg saddle, 44.636°N 123.2972°W, 314m. 06.v.2016. |
| *Pterostichus melanarius* | 088 | USA: OR: Benton Co., Truax Island, 44.586°N 123.1858°W, 61m. 25.vii.2015. |
| *Rhadine dissecta-*group sp. | 315 | USA: OR: Harney Co., Alvord desert dunes nr. bluff, 42.6753°N 118.3695°W, 1230m. 04.vi.2016. |
| *Scaphinotus marginatus* | 176 | USA: OR: Benton Co., Corvallis, Crystal Lake Sports Complex, 44.5497°N 123.2453°W, 09.iv.2016. |
| *Scaphinotus marginatus* | 182 | USA: OR: Benton Co., Corvallis, Crystal Lake Sports Complex, 44.5497°N 123.2453°W, 09.iv.2016. |
| *Scarites (Distichus)* sp. | 651 | Mozambique: Sofala: Parque Nacional da Gorongosa, Chitengo camp, 18.9773°S 34.3515°E, 21m. 08-15.ii.2018. |
| *Scarites (Parallelomorphus)* sp. | 629 | RSA: Limpopo: Makuya Nature Reserve, 22.427°S 31.0532°E, 274m. 02.ii.2018. |
| *Scarites (Parallelomorphus)* sp. | 630 | RSA: Limpopo: Makuya Nature Reserve, 22.427°S 31.0532°E, 274m. 02.ii.2018. |
| *Scarites marinus* | 435 | USA: FL: Monroe Co., Long Key State Park, 24.8105°N 80.829°W, 17m. 18.ix.2016. |
| *Scarites marinus* | 436 | USA: FL: Monroe Co., Long Key State Park, 24.8105°N 80.829°W, 17m. 18.ix.2016. |
| *Scarites marinus* | 460 | USA: FL: Monroe Co., Long Key State Park, 24.8105°N 80.829°W, 17m. 18.ix.2016. |
| *Schizogenius litigiosus* | 231 | USA: OR: Benton Co., Irish Bend County Park, 44.3647°N 123.2206°W, 79m. 09.v.2016. |
| *Schizogenius litigiosus* | 232 | USA: OR: Benton Co., Irish Bend County Park, 44.3647°N 123.2206°W, 79m. 09.v.2016. |
| *Schizogenius litigiosus* | 233 | USA: OR: Benton Co., Irish Bend County Park, 44.3647°N 123.2206°W, 79m. 09.v.2016. |
| *Selenophorus* sp. | 362 | USA: AZ: Pinal Co., Peppersauce Canyon nr campground, 32.5372°N 110.7195°W. 06.viii.2016. |
| *Semiardistomis viridis* | 305 | USA: AR: Newton Co., Shop Creek near Highway 327, 35.9511°N 93.2434°W. 16.vi.2016. |
| *Semiardistomis viridis* | 317 | USA: AR: Montgomery Co., Ouachita River, Fulton Branch Recreation Area, 34.6265°N 93.6589°W. 17.vi.2016. |
| *Semiardistomis viridis* | 511 | USA: Louisiana: Webster Parish, Kisatchie Nat. Forest, Caney Lakes Recreation Area, 32.6758°N 93.2928°W, 69m. 28.viii.2017. |
| *Semiardistomis viridis* | 512 | USA: Louisiana: Webster Parish, Kisatchie Nat. Forest, Caney Lakes Recreation Area, 32.6758°N 93.2928°W, 69m. 28.viii.2017. |
| *Semiardistomis viridis* | 513 | USA: Louisiana: Webster Parish, Kisatchie Nat. Forest, Caney Lakes Recreation Area, 32.6758°N 93.2928°W, 69m. 28.viii.2017. |
| *Semiardistomis viridis* | 515 | USA: Arkansas: Washington Co., Devil's Den State Park, Lee Creek bank, 35.7817°N 94.2494°W, 310m. 30.viii.2017. |
| *Sericoda bembidioides* | 420 | USA: AZ: Cochise Co., Chiricahua Mtns, Rustler Park, 31.9068°N 109.2769°W, 2548m. 18.viii.2016. |
| *Sphaeroderus schaumii* | 667 | USA: VA: Fauquier Co., G. R. Thompson WMA, 38.966°N 78.017°W. 22.i.2018. |
| *Sphaeroderus schaumii* | 668 | USA: VA: Fauquier Co., G. R. Thompson WMA, 38.966°N 78.017°W. 22.i.2018. |
| *Sphaeroderus stenostomus* | 659 | USA: VA: Fauquier Co., G. R. Thompson WMA, 38.966°N 78.017°W. 22.i.2018. |
| *Sphaeroderus stenostomus* | 660 | USA: VA: Fauquier Co., G. R. Thompson WMA, 38.966°N 78.017°W. 22.i.2018. |
| *Sphaeroderus stenostomus* | 661 | USA: VA: Fauquier Co., G. R. Thompson WMA, 38.966°N 78.017°W. 22.i.2018. |
| *Sphaeroderus stenostomus* | 662 | USA: VA: Fauquier Co., G. R. Thompson WMA, 38.966°N 78.017°W. 22.i.2018. |
| *Sphaeroderus stenostomus* | 663 | USA: VA: Fauquier Co., G. R. Thompson WMA, 38.966°N 78.017°W. 22.i.2018. |
| *Stenocrepis elegans* | 391 | USA: AZ: Pinal Co., Winkelman, AZ, 32.9852°N 110.7676°W, 587m. 11.viii.2016. |
| *Stenognathus quadricollis* | 112 | Mexico: Veracruz: Est. Biol. Los Tuxtlas, nr. Catemaco and Montepio, 18.5855°N 95.0752°W, 125m. 06-09.viii.2015. |
| *Stenolophus* sp. | 309 | USA: OR: Lake Co., Musser Reservoir, 43.5535°N 120.0074°W, 1230m. 05.vi.2016. |
| *Stenomorphus convexior* | 402 | USA: AZ: Pima Co., Happy Valley, oak forest, 32.1559°N 110.4748°W, 1262m. 14.viii.2016. |
| *Stenomorphus convexior* | 403 | USA: AZ: Pima Co., Happy Valley, oak forest, 32.1559°N 110.4748°W, 1262m. 14.viii.2016. |
| *Stenomorphus convexior* | 418 | USA: AZ: Cochise Co., S Blue Sky Rd S of Wilcox, AZ, 32.2335°N 109.7774°W, 1269m. 17.viii.2016. |
| *Stolonis intercepta* | 330 | USA: AZ: Pima Co., AZ DOT rest stop, northbound I19, 31.7653°N 111.0351°W, 914m. 01-03.viii.2016. |
| *Stolonis intercepta* | 331 | USA: AZ: Pima Co., Canoa Ranch Rest Area, southbound I19, 31.7653°N 111.0351°W, 914m. 01-03.viii.2016. |
| *Stolonis* sp. | 116 | Mexico: Chiapas: roadside stop near Armando Zebadea pueblo, 16.9291°N 93.4737°W, 942m. 10.viii.2015. |
| *Striganoviella vanhillei* | 605 | RSA: Eastern Cape: Mkhambathi Nature Reserve, Gwegwe, estuary, 31.2878°S 30.0102°E, 12m. 25.i.2018. |
| *Striganoviella vanhillei* | 606 | RSA: Eastern Cape: Mkhambathi Nature Reserve, Gwegwe, estuary, 31.2878°S 30.0102°E, 12m. 25.i.2018. |
| *Striganoviella vanhillei* | 607 | RSA: Eastern Cape: Mkhambathi Nature Reserve, Gwegwe, estuary, 31.2878°S 30.0102°E, 12m. 25.i.2018. |
| *Striganoviella vanhillei* | 608 | RSA: Eastern Cape: Mkhambathi Nature Reserve, Gwegwe, estuary, 31.2878°S 30.0102°E, 12m. 25.i.2018. |
| *Striganoviella vanhillei* | 609 | RSA: Eastern Cape: Mkhambathi Nature Reserve, Gwegwe, estuary, 31.2878°S 30.0102°E, 12m. 25.i.2018. |
| *Striganoviella vanhillei* | 611 | RSA: Eastern Cape: Mkhambathi Nature Reserve, Gwegwe, estuary, 31.2878°S 30.0102°E, 12m. 25.i.2018. |
| *Syntomus americanus* | 205 | USA: OR: Benton Co., Marys Peak, 44.5102°N 123.5509°W, 1153m. 19.iv.2016. |
| *Syntomus americanus* | 206 | USA: OR: Benton Co., Marys Peak, 44.5102°N 123.5509°W, 1153m. 19.iv.2016. |
| *Synuchus dubius* | 425 | USA: AZ: Cochise Co., Chiricahua Mtns, Rustler Park, 31.9068°N 109.2769°W, 2548m. 18.viii.2016. |
| *Synuchus dubius* | 426 | USA: AZ: Cochise Co., Chiricahua Mtns, Rustler Park, 31.9068°N 109.2769°W, 2548m. 18.viii.2016. |
| *Synuchus dubius* | 427 | USA: AZ: Cochise Co., Chiricahua Mtns, Rustler Park, 31.9068°N 109.2769°W, 2548m. 18.viii.2016. |
| *Synuchus dubius* | 428 | USA: AZ: Cochise Co., Chiricahua Mtns, Rustler Park, 31.9068°N 109.2769°W, 2548m. 18.viii.2016. |
| *Synuchus dubius* | 429 | USA: AZ: Cochise Co., Chiricahua Mtns, Rustler Park, 31.9068°N 109.2769°W, 2548m. 18.viii.2016. |
| *Tachyta inornata* | 258 | USA: OR: Jefferson Co., pine forest burn site off of road NF-14, 44.5358°N 121.6219°W, 1159m. 14.v.2016. |
| *Tachyta inornata* | 263 | USA: OR: Jefferson Co., pine forest burn site off of road NF-14, 44.5358°N 121.6219°W, 1159m. 14.v.2016. |
| *Tachyura rapax* | 264 | USA: OR: Benton Co., Irish Bend County Park, 44.3647°N 123.2206°W, 79m. 10.v.2016. |
| *Tetracha carolina* | 392 | USA: AZ: Pinal Co., Winkelman, AZ, 32.9852°N 110.7676°W, 587m. 11.viii.2016. |
| *Tetracha carolina* | 393 | USA: AZ: Pinal Co., Winkelman, AZ, 32.9852°N 110.7676°W, 587m. 11.viii.2016. |
| *Tetracha carolina* | 394 | USA: AZ: Pinal Co., Winkelman, AZ, 32.9852°N 110.7676°W, 587m. 11.viii.2016. |
| *Tetragonoderus fasciatus* | 332 | USA: AZ: Pima Co., AZ DOT rest stop, northbound I19, 31.7653°N 111.0351°W, 914m. 01-03.viii.2016. |
| *Tetragonoderus fasciatus* | 334 | USA: AZ: Pima Co., AZ DOT rest stop, northbound I19, 31.7653°N 111.0351°W, 914m. 01-03.viii.2016. |
| *Tetragonoderus fasciatus* | 344 | USA: AZ: Pima Co., Canoa Ranch Rest Area, southbound I19, 31.7653°N 111.0351°W, 914m. 01-03.viii.2016. |
| *Tetragonoderus fasciatus* | 345 | USA: AZ: Pima Co., Canoa Ranch Rest Area, southbound I19, 31.7653°N 111.0351°W, 914m. 01-03.viii.2016. |
| *Tetragonoderus fasciatus* | 346 | USA: AZ: Pima Co., Canoa Ranch Rest Area, southbound I19, 31.7653°N 111.0351°W, 914m. 01-03.viii.2016. |
| *Tetragonoderus* sp. nr. *latipennis* | 133 | Mexico: Oaxaca: stop off of Highway 125 nr. La Esperanza, 16.4893°N 98.1085°W, 87m. 17.viii.2015. |
| *Thyreopterus flavosignatus* | 604 | RSA: KwaZulu-Natal: Highover Wildlife Sanctuary, 29.9135°S 30.0996°E, 546m. 27.i.2018. |
| *Trachypachus inermis* | 238 | USA: OR: Lincoln Co., Moolack Beach, 44.7093°N 124.0605°W, 4m. 21.v.2016. |
| *Trachypachus slevini* | 240 | USA: OR: Lincoln Co., Moolack Beach, 44.7093°N 124.0605°W, 4m. 21.v.2016. |
| *Trachypachus slevini* | 241 | USA: OR: Lincoln Co., Moolack Beach, 44.7093°N 124.0605°W, 4m. 21.v.2016. |
| *Trachypachus slevini* | 294 | USA: OR: Lincoln Co., Moolack Beach, 44.7093°N 124.0605°W, 4m. 21.v.2016. |
| *Trachypachus slevini* | 295 | USA: OR: Lincoln Co., Moolack Beach, 44.7093°N 124.0605°W, 4m. 21.v.2016. |
| *Trechodes* sp. | 639 | RSA: Limpopo: Mashovhela Bush Lodge, 22.944°S 29.8907°E, 512m. 04.ii.2018. |
| *Trechosiella scotti* | 581 | RSA: KwaZulu-Natal: Royal Natal National Park, 28.7117°S 28.9371°E, 1540m. 18.i.2018. |
| *Trechosiella scotti* | 582 | RSA: KwaZulu-Natal: Royal Natal National Park, 28.7117°S 28.9371°E, 1540m. 18.i.2018. |
| *Trechosiella scotti* | 583 | RSA: KwaZulu-Natal: Royal Natal National Park, 28.7117°S 28.9371°E, 1540m. 18.i.2018. |
| *Trechus humboldti* | 559 | USA: OR: Benton Co., Prarie Peak, 44.284°N 123.593°W. 15.xii.2017. |
| *Trechus humboldti* | 560 | USA: OR: Benton Co., Prarie Peak, 44.284°N 123.593°W. 15.xii.2017. |
| *Trechus humboldti* | 561 | USA: OR: Benton Co., Prarie Peak, 44.284°N 123.593°W. 15.xii.2017. |
| *Trechus humboldti* | 562 | USA: OR: Benton Co., Prarie Peak, 44.284°N 123.593°W. 15.xii.2017. |
| *Zacotus matthewsii* | 219 | USA: OR: Benton Co., Marys Peak, trail nr boundary btwn forest and meadow, 44.5091°N 123.5526°W, 1141m. 19.iv.2016. |
| *Zacotus matthewsii* | 220 | USA: OR: Benton Co., Marys Peak, trail nr boundary btwn forest and meadow, 44.5091°N 123.5526°W, 1141m. 19.iv.2016. |
| *Zacotus matthewsii* | 221 | USA: OR: Benton Co., Marys Peak, trail nr boundary btwn forest and meadow, 44.5091°N 123.5526°W, 1141m. 19.iv.2016. |
| *Zacotus matthewsii* | 230 | USA: OR: Benton Co., McDonald-Dunn forest, Lewisburg saddle, 44.636°N 123.2972°W, 314m. 06.v.2016. |
